# Applying nutritional ecology to optimize diets of crickets raised for food and feed

**DOI:** 10.1101/2024.05.23.595553

**Authors:** Matthew J Muzzatti, Sarah J. Harrison, Emily R. McColville, Caelyn T. Brittain, Hunter Brzezinski, Sujitha Manivannan, Cassandra C. Stabile, Heath A. MacMillan, Susan M. Bertram

## Abstract

Increasing yield is a primary goal of mass insect rearing for food and feed, and diet shapes insect life history traits important to yield, such as survival, development time, and body size at adulthood. Little is known about how developmental macronutrient intake impacts survival, growth, and adult body size of mass reared insects. Here, we applied the nutritional geometry framework and reared individual tropical house crickets (*Gryllodes sigillatus*) from hatch to adulthood on a wide range of protein:carbohydrate diets. We measured weekly food consumption, survival, development time to adulthood, and adult body size and mass, and calculated a yield metric to extrapolate our individual-level results and predict how diet influences yield at the mass rearing level. Yield was maximized on a 3_P_:1_C_ diet, as crickets fed this diet were most likely to develop into adults and grew maximum mass and body size. When provided with a choice between diets, crickets selected a relatively balanced 1.05_P_:1_C_ diet throughout development, but males consumed 17% more protein than females. Our results represent a crucial first step towards determining the optimal standard feed formulation required to maximize cricket farming yield.

## 1.0 Introduction

Food security is a major global concern for the future. The current global agricultural system does not produce enough protein to meet global nutritional demands [1], and with a rising population expected to reach 9.7 billion people by 2050 [2], there is an urgent need for more protein. Insects, a sustainable and nutritious alternative protein source, are one potential solution to this problem. Insects are generally high in protein, vitamins, and minerals [3–6], and their production requires considerably less resources than other protein sources such as beef or chicken [7,8]. Crickets are one of the most common insects raised for food and feed [9], and are used principally by the pet food and aquafeed markets [10,11]. Increasing yield is a primary goal of agricultural research, and upscaling insect production to achieve higher yields is one way to meet world demand for protein [12]. Yield is a measurement of product harvested per unit area, and cricket farms struggle with how best to increase yields at a scale of billions of crickets required for a farming environment without the costs of additional labour. Techniques for increasing insect farm yields are not well understood [13], so an integrative approach linking animal ecology and physiology with industrial application may prove useful [14].

To a biologist, yield can be thought of as a product of life history traits; specifically, the body size or mass of individuals at harvest, the development time required to reach the desired life stage, and the proportion of individuals that survive to harvest. Extensive research has examined the regulation of insect body size and development time allowing for predictions of how natural selection shapes these life history traits [15–22]. Most of what we know about insect body size and development time has been derived from experiments using *Manduca sexta* as a model organism. *Manduca sexta* is a holometabolous insect that begins development as a larva, which must proceed through multiple larval feeding stages separated by molting events to reach a critical weight prior to pupating and undergoing complete metamorphosis to reach adulthood [18,19]. Final adult body size in holometabolous insects is determined by the interactions between body size achieved at the end of larval development, steroid hormones, and nutritional signalling [23,24]. In comparison, hemimetabolous insects do not pupate or metamorphose, but instead develop to adulthood by continuously secreting, digesting, and reassembling their cuticle through multiple nymphal instars before achieving their final size [25,26]. It is unclear whether the rules governing holometabolous insect body size and developmental time hold for hemimetabolous insects. What is clear is that diet is a strong determinant of insect fitness and the proportion of proteins and carbohydrates are especially important for optimizing insect life history traits [27–31]. Most early life nutrition experiments have been conducted using oligidic diets, which are diets that differ in the amounts of several natural ingredients. As these natural ingredients also differ in vitamins, minerals, and lipids, it is impossible to ascertain which nutrients are impacting early life history traits (e.g., carbohydrate and protein levels, or vitamin, mineral, or lipid levels).

The nutritional geometry framework [32,33] has been successfully applied to many insects using holidic diets (artificial diets containing chemically-defined ingredients; Dadd, 1960) to test how different proportions of key dietary nutrients influence life history traits. This approach graphically represents the nutritional requirements of an organism and considers how the consumption of two nutrients influences a life history trait, resulting in a ‘nutrient landscape’. Holidic diets are diluted with non-nutritional cellulose to ensure that some individuals consume low levels of nutrients to allow resolution in the lower-intake regions of the nutritional landscape to spread out nutrient intake along ‘nutritional rails’ [32,33]. A nutritional rail is a line in the nutritional landscape, the slope of which represents the ratio of nutrients in a particular diet. This approach has been particularly successful in examining how diet shapes cricket life history responses. For example, diets higher in protein can maximize female egg production and overall fecundity, while diets higher in carbohydrate regulate male reproductive fitness and behaviour [29,35–40]. Both carbohydrate and protein intake influence adult cricket weight gain and survival, with protein intake regulating adult weight gain and carbohydrate intake regulating adult lifespan [29,41]. Given the profound effect diet has on insect life history in the laboratory setting, the ratio of proteins to carbohydrates could strongly influence cricket yield in mass production farm settings. Therefore, ascertaining the ideal protein to carbohydrate ratio (P:C) that minimizes development time and maximizes body size, mass and survival has important implications for producing more edible protein [42,43].

Experiments using oligidic diets have demonstrated that early life nutrition and lifetime nutrition can be extremely important at influencing insect life history [39,44–48], but less research exists that teases out the effects of individual nutrients on early life history using holidic diets. It was long assumed that holidic diets could not be used to successfully rear insects throughout development [39]. As a result, most holidic diet experiments have been conducted using adults or late-stage juveniles, which overlooks the relative importance of early life and/or lifetime nutrient acquisition and allocation [35,41,49–57]. This assumption has since been invalidated, as a few studies have demonstrated that when fed from hatch to adulthood, diets proportionally higher in protein compared to carbohydrates can result in heavier crickets (*Acheta domesticus*; Gutiérrez et al., 2020) and more fecund caterpillars (*Heliothis virescens*; Roeder and Behmer, 2014) and crickets (*Gryllus bimaculatus*; Han and Dingemanse, 2017). Research testing the developmental effects of holidic diets spanning a wide P:C can be applied to mass reared crickets to better understand what proportion of nutrients can influence, and optimize, life history traits important to yield.

*Gryllodes sigillatus* (Gryllidae: Orthoptera) is a prominent species farmed for food and feed in North America with world-wide distribution [58]. The geometric framework has previously been applied to adult *G. sigillatus* in experiments that tested short feeding intervals from 10-21 days on artificial diets spanning a 5_P_:1_C_ – 1_P_:8_C_ range and four different nutrient concentrations [35,41,52,56]. These short feeding intervals were enough to elicit changes in adult immunity and reproductive life history traits. A high intake of nutrients in a relatively balanced diet maximized male reproductive traits, mating success, and female egg production [35,41,52]. High protein diets increased mortality, whereas high carbohydrate diets maximized calling effort and adult lifespan [41,56]. When given a choice between high carbohydrate and high protein diets, both adult sexes ate significantly more carbohydrate than protein and regulated their nutrient intake at ∼1_P_:2_C_ [41,52]. These results clearly demonstrate that diet at adulthood can strongly influence *G. sigillatus* life history and behaviour, but it remains unclear how early life nutrient acquisition influences juvenile *G. sigillatus* development and survival.

Here, we ran two independent experiments on *G. sigillatus* from hatch to adulthood. In the first experiment, we asked: (a) what proportion of proteins and carbohydrates maximizes cricket yield (i.e., results in the shortest development time, largest body size, heaviest mass, and greatest survival), and (b) are these responses sex-specific? We hypothesized that higher protein diets would result in larger body size and heavier mass, and higher carbohydrate diets would result in longer development time. We also hypothesized the effects would be sex specific as the same dietary P:C can have a different influence on males and females, possibly indicative of sexual conflict [37,46,52,55,59–61]. In the second experiment, we asked (a) how juvenile *G. sigillatus* regulates P:C intake when given a choice between pure carbohydrate and pure protein diets, b) whether P:C intake regulation changes throughout development, and (c) whether these responses are sex-specific. We hypothesized that the intake target of *G. sigillatus* would be different throughout development. An intake target range of ∼1_P_:2_C_ has been reported for *G. sigillatus* adults [41,52], but hemimetabolous insects such as crickets undergo multiple resource-intensive moults throughout development and experience changing physiological needs between moulting and foraging, and so macronutrient requirements may vary throughout development. We also hypothesized that, as in *Gryllus veletis* [29] and *Teleogryllus commodus* [37], dietary preference will not match with the diet that maximizes growth rate and would differ between the sexes. To our knowledge, our study is the first to test how a wide range of dietary P:C ratios and nutrient concentrations, fed from hatch to adulthood, influence life history traits in a mass reared hemimetabolous insect.

## 2.0 Methods

All experimental crickets were received as eggs from a commercial cricket supplier (Entomo Farms, Norwood, Ontario, Canada). The colony was fed ad libitum water and food (Earth’s Harvest Organic Cricket Grower; Earth’s Harvest, Oxford Mills, Ontario, Canada) that reflects a P:C of approximately 1_P_:1.9_C_. The eggs were laid in a medium of peat moss and maintained inside an incubator (Thermo Fisher Scientific Inc., Massachusetts, United States) at 32°C and 60% relative humidity. A 14:10-hour light:dark cycle was maintained using an LED installed alongside the incubator interior. These conditions were maintained for all experiments. Different batches of eggs from different batches of adults were used for each trial in these experiments. While parental contributions can drive variation in resource acquisition [62], egg-laying adults were always fed and treated in the exact same manner.

### 2.1 No choice of P:C

Experimental holidic diets were created following established protocols [29,34]. Diets consisted of seven different P:C ratios (8_P_:1_C_, 5_P_:1_C_, 3_P_:1_C_, 1_P_:1_C_, 1_P_:3_C_, 1_P_:5_C_, 1_P_:8_C_), each diluted to three cellulose levels (14, 45, 76%), for a total of 21 unique diets to spread-out nutrient intake along ‘nutritional rails’ and enable resolution in the lower-intake regions of the nutritional landscape [32,33]. Within 48 hours of egg emergence, 420 cricket nymphs were each placed into individual rearing containers consisting of 96.1-ml plastic condiment cups with aerated lids, a 14-mm-wide polyethylene push-in cap for a food dish, and a 0.75-uL polypropylene test tube (lid removed) for a water vial stoppered with 38-mm-wide saturated dental cotton wick (Healifty, Guangdong Province, China). Each cricket was pseudorandomly assigned to a diet, resulting in N=20 individuals per diet. Diets were fed from hatch to adulthood. Weekly, crickets were provided with fresh food; protein and carbohydrate intakes were calculated for each individual from total food intake (difference in dry weight of food dish before and after consumption) and known food compositions. Food was dried in an oven containing dishes of blue indicating silica gel desiccant beads (Dry & Dry, Brea, California, USA) at 30°C for 96 h before crickets were provided with food and again after one week of consumption (prior to weighing). Crickets were monitored three times a week for survival and time to adulthood. Individuals were either observed until their natural death or until 1-week post eclosion to adulthood when they were euthanized. Upon adult eclosion crickets were weighed using a Mettler Toledo model AB135-S analytical scale and photographed using a Zeiss Stemi 305 Stereo Microscope with an Axiocam 208 camera for body size measurements using ImageJ, version 1.48 software (National Institutes of Health, Bethesda, Maryland, United States of America). Body size measurements included head width, pronotum width, and pronotum length, and were averaged across three independent measurers to reduce measurement bias. Cricket nymphs are fragile, and some of these diets are stressful, so if a nymph died within the first two weeks of life it was documented and replaced with a new hatchling. Including replacement, a total of 928 total crickets were used in this experiment. From these 928 individuals, 87 females and 61 males were used in the linear models and nutritional landscapes because we only included individuals of known sex who also reached adulthood. An additional female that reached adulthood, but was missing body size and mass measurements, was included in the development time models and landscapes. All 928 individuals were used in the cox proportional hazards models because these models can handle censored individuals of unknown sex.

### 2.2 Statistical analyses – no choice of P:C

All statistical analyses were performed in R version 4.2.2 [63] and RStudio version 2022.07.2 [64]. Principal component analysis was used to extract orthogonal vectors from head width, pronotum width, and pronotum length at adulthood to quantify adult body size into a single measurement (PC1). The first principal component explained 95.4% of the variation (eigenvalue = 2.86) and had equal loadings for all variables. We used a multivariate surface-response approach to estimate and visualize the linear, quadratic, and correlational effects of amount of protein and carbohydrate eaten on body mass, body size PC1, and development time; response surfaces were visualized using non-parametric thin-plate splines using the ‘fields’ package [65]. The ‘OptimaRegion’ package [66] was used to generate 95% confidence regions (CR) and the global maximum for each response surface [41,67]. The default termination criterium was used for each 95% CR, and 500 simulations were generated for each plot. Due to a limitation of the ‘OptimaRegion’ package, for those response surfaces whose intake data did not exactly conform to a triangle shape in the nutritional landscape, it was necessary to visually extrapolate the vertices of the triangle to estimate 95% CR. Global maxima were mapped to the response surfaces, and all CR plots can be found in the Supplementary Material.

We used general linear models from the ‘stats’ package [63] to analyse how linear, quadratic, and correlational terms for protein and carbohydrate intake influenced adult mass, adult body size, and development time to adulthood. The false discovery rate B-Y method (FDR_BY_) was used to adjust alpha for each general linear model [68]. Male and female data were pooled and analysed together, as these pooled data are directly relevant to a farm environment because harvest (and final yield) combines both males and females. We also investigated the effects of diet on males and females separately because sex-specific life-history differences in response to diet are well documented in insects, including in crickets [29,37,39,54,55].

Cox proportional hazard models were applied using the ‘survival’ package [69,70] to determine how dietary P:C and cellulose concentration influenced development time to adulthood and survival time. Censored individuals were those that were lost during the experiment, or those juveniles or adults who did not die prior to the end of the experiment; censored status data are provided in Table S1. The predictor terms for the most complex survival model were P:C (categorical), cellulose concentration (categorical), and their interaction. To satisfy the assumption of proportional hazards for the survival time model, it was necessary to estimate the coefficient for cellulose concentration at three different time intervals by stratifying the data by survival times of 0-7 days, 7-14 days, and 14-153 days. We used the ‘MuMIn’ package [71] to perform model selection and identify the ‘best approximating model’ for our development time and survival time data. We then used Akaike’s Information Criterion, and calculated ΔAIC values by finding the difference between each model’s AIC score and that of the top ranked (lowest AIC) model [72,73]. We established our penultimate confidence set by rejecting models with ΔAIC < 4 (Tables S5 and S7), and established our final confidence set by further rejecting models that were more complex variants of any model with a lower AIC value (nesting approach; Richards, 2008; Richards et al., 2011). Our top models for both development time and survival time had unequivocal support (Akaike weights = 1).

We calculated a yield metric for each P:C using the following formula:

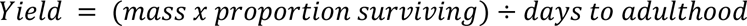

where mass and days to adulthood represented the average values of adults within a P:C, and proportion surviving was representative of both juveniles and adults within a P:C. Standard error (σ) for yield was calculated using the error propagation formula:

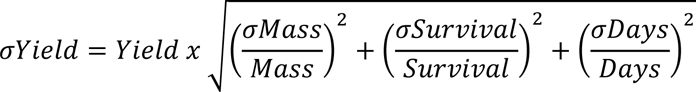

A 95% confidence interval for each yield metric was calculated using the following formula:

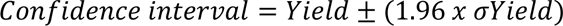

where 1.96 represents the critical value associated with a 95% confidence interval.

### 2.3 Choice of P:C

To test for preferences in dietary protein and carbohydrates, we used experimental holidic diets consisting of three different protein P:C ratios (1_P_:0_C_, 1_P_:1_C_, 0_P_:1_C_) that were each diluted to a 45% cellulose level. A 45% cellulose level was selected based on other insect diet choice experiments that used a similar level of nutrient dilution (46%; Harrison et al., 2014; 42%; Ng et al., 2019). Upon emergence, 120 crickets were distributed evenly across four group-living containers and were fed the 1_P_:1_C_ diet to prevent the high levels of mortality recorded in the no-choice experiment. Crickets were provided with fresh food twice during the first week, and daily feed intake was calculated as the difference between fresh and eaten-on feed dishes / 30 crickets / number of days spent feeding. Food was dried as in the first experiment. After the first week, an even number of crickets from each of the group-living containers were weighed and photographed for body size measurements and then randomly distributed into 60 individual 18.4 x 12.7 x 5.08 cm (length x width x height) 709.8 mL clear plastic take-out containers with aerated lids (Platinum Crown Corporation Limited, Colchester, United Kingdom). Crickets were then provided with two feed dishes containing the 1_P_:0_C_ and 0_P_:1_C_ diets. If crickets ate their two diet choices at random, then they should have selected a 1_P_:1_C_. Position of the feed dishes were randomly assigned across containers to prevent preference for left or right handedness. Crickets were provided with fresh food weekly and fresh water three times a week. For the duration of the experiment (8 weeks), diet consumption was measured weekly until 1-week post adult eclosion. Survival and time to adult eclosion were monitored five times a week. From the original 60 crickets, 18 females and 16 males survived to 1-week post adult eclosion and were used in data analysis.

### 2.4 Statistical analyses – choice of P:C

Principal component analysis was used as described in section 2.1.2 to create a single summary variable for body size (PC1). The first principal component explained 96.4% of the variation (eigenvalue = 2.89) and had equal loadings for all variables. We used a generalized linear mixed-effects regression (family – Gamma, link – log) using the ‘lme4’ package [76] to test if amount of diet eaten was influenced by food type (pure protein or pure carbohydrate), whether food preferences changed over time, and if the consumption of each food type differed between the sexes. We included week, diet, sex, and all possible interactions as fixed effects, week 1 body size PC1 as a covariate, and included cricket ID as a random effect. As described in section 2.2, we performed model selection and identified the ‘best approximating model’ for our diet intake data. Our candidate model set consisted of nine models, in which all four main effects were fixed. After rejecting models with ΔAIC < 4, and further rejecting models using the nesting approach, the ‘best approximating model’ for amount of diet eaten had unequivocal support (Akaike Weight = 1). The observed mean P:C intakes were calculated by summing the total protein and total carbohydrates consumed across 8 weeks and averaging across individuals. These means were then compared with the expected intake of 1:1 by using one sample t-tests with two tailed probabilities. Normality was confirmed using Shapiro-Wilk tests; to meet normality assumptions, male data was square-root transformed, female data was inverse transformed, and all (pooled) data was log10 transformed. Reported P:C ratios are back transformed.

## 3.0 Results

### 3.1 No choice of P:C

Protein and carbohydrate intake influenced cricket body size (PC1) (R^2^ = 0.43; F_5,154_ = 24.48; *P* < 0.001), mass (R^2^ = 0.45; F_5,154_ = 26.57; *P <* 0.001) and development time (R^2^ = 0.45; F_5,155_ = 27.22; *P* < 0.001). Crickets had the largest body size at a high intake of P and C centered around a ratio of 3.14_P_:1_C_ (global maximum: P = 0.546 mg, C = 0.174 mg; Figures 1a, S1), and this effect was explained by significant linear relationships between body size and protein and body size and carbohydrate (both *P* < 0.001; Table S2). This body size effect was also explained by a significant non-linear relationship with protein (*P* < 0.001; Table S2) such that body size peaked and then declined with increasing protein intake.

**Figure 1.**
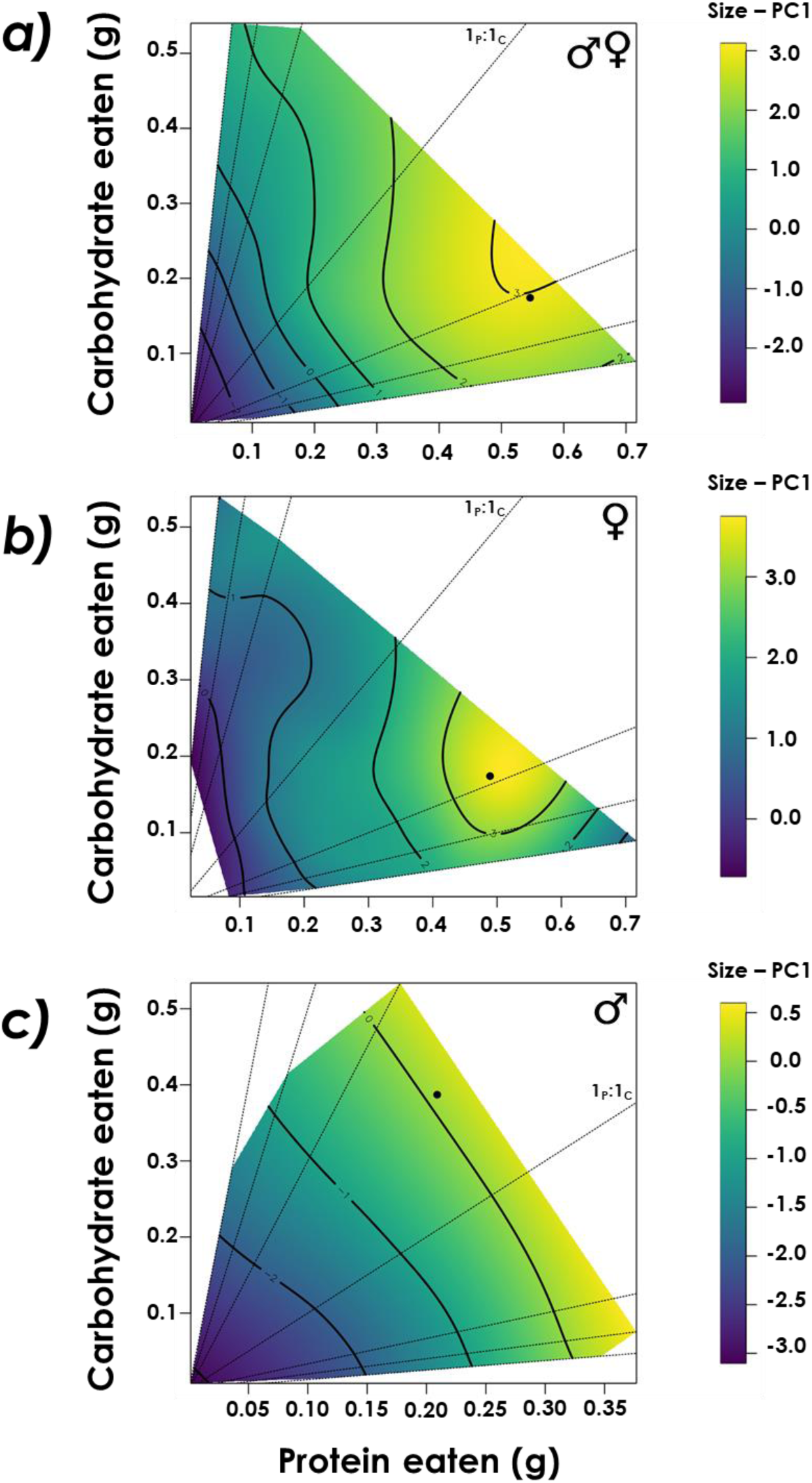
Response surfaces that illustrate the response of dietary protein and carbohydrate intake on pooled (a), female (b), and male (c) *Gryllodes sigillatus* body size (PC1) and the corresponding global maximum (closed black circle). Dotted lines represent nutritional rails defined by the protein:carbohydrate ratios in the experimental diets. As crickets develop and feed on their individual diets, they move along their nutritional rails and grow, represented by the colour gradient. The 1_P_:1_C_ nutritional rail is labelled on each panel.

When examining the sexes separately, protein intake relative to carbohydrates had the greatest influence on female body size (Figure 1b; Table S3). Female body size was explained by significant relationships with protein intake, both linear and non-linear (both *P* < 0.001; Table S2); female body size peaked around a ratio of 2.81_P_:1_C_ (global maximum: P = 0.489 mg, C = 0.174 mg; Figures 1b, S1) and then declined with increasing protein intake. Neither protein or carbohydrate intake significantly influenced male body size (both *P* > 0.05; Tables S1, S2), and the maximum male body size was achieved around a ratio of 1_P_:1.85_C_ (global maximum: P = 0.209 mg, C = 0.387 mg; Figures 1c, S1).

Cricket mass was explained by significant linear and nonlinear relationships with protein intake (both *P* < 0.001; Table S2) and a significant positive linear relationship with carbohydrate intake (*P* = 0.0028; Table S2), such that the greatest mass peaked around a ratio of 3.22_P_:1_C_ (global maximum: P = 0.532 mg, C = 0.165 mg; Figures 2a, S2) and then declined with increasing protein intake. Like body size, protein intake relative to carbohydrates had the greatest influence on female mass (both linear and non-linear effects, both *P* < 0.001; Table S3). Female mass was explained by a significant non-linear relationship with protein (*P* < 0.001; Table S2); females grew the heaviest around a ratio of 3.29_P_:1_C_ (global maximum: P = 0.523, C = 0.159; Figures 2b, S2), but then declined in mass with increasing protein intake. However, neither protein nor carbohydrate intake significantly influenced male mass (both *P* > 0.05; Table S3). Males grew the heaviest around a ratio of 2.20_P_:1_C_ (global maximum: P = 0.310 mg, C = 0.141 mg; Figures 2c, S2).

**Figure 2.**
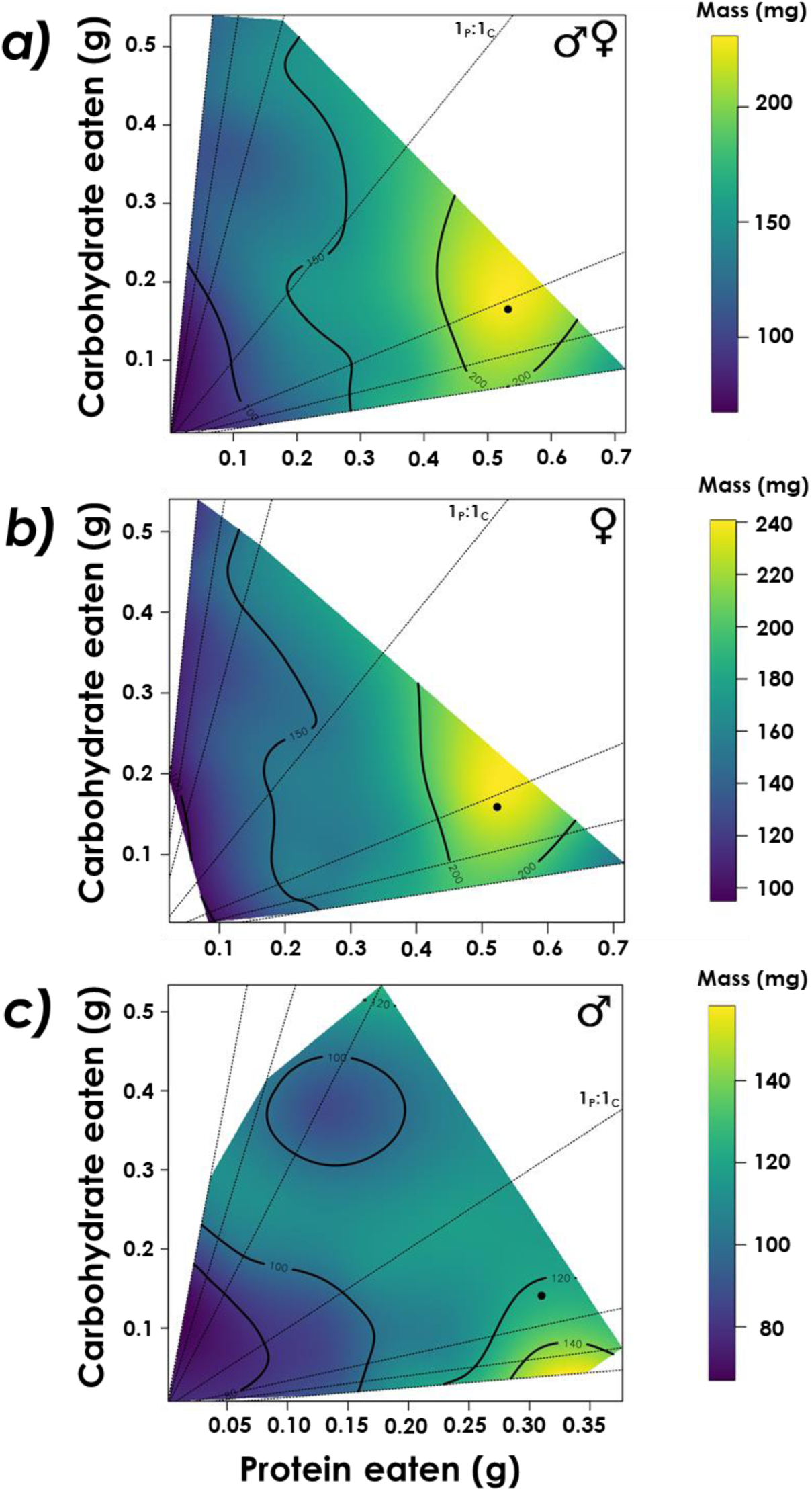
Response surfaces that illustrate the response of dietary protein and carbohydrate intake on pooled (a), female (b), and male (c) *Gryllodes sigillatus* mass and the corresponding global maximum (closed black circle) on each landscape. Dotted lines represent nutritional rails defined by the protein:carbohydrate ratios in the experimental diets. As crickets develop and feed on their individual diets, they move along their nutritional rails and gain mass, represented by the colour gradient. The 1_P_:1_C_ nutritional rail is labelled on each panel.

Development time was described by a significant positive linear relationship with carbohydrate intake (*P* = 0.0014; Table S2). Crickets developed the slowest around a ratio of 1_P_:6.72_C_ (global maximum: P = 0.081 mg, C = 0.544 mg; Figures 3a, S3). Neither protein nor carbohydrate intake significantly influenced female development time (both *P* > 0.05; Tables S1, S2). Females developed the slowest around a ratio of 1_P_:1.98_C_ (global maximum: P = 0.202 mg, C = 0.400 mg; Figures 3b, S3). Carbohydrate intake relative to protein intake significantly influenced male development time, an effect described by a significant positive linear relationship with carbohydrates (*P* = 0.0033; Tables S1, S2). Males developed the slowest around a ratio of 1_P_:6.97_C_ (global maximum: P = 0.093 mg, C = 0.648; Figures 3c, S3).

**Figure 3.**
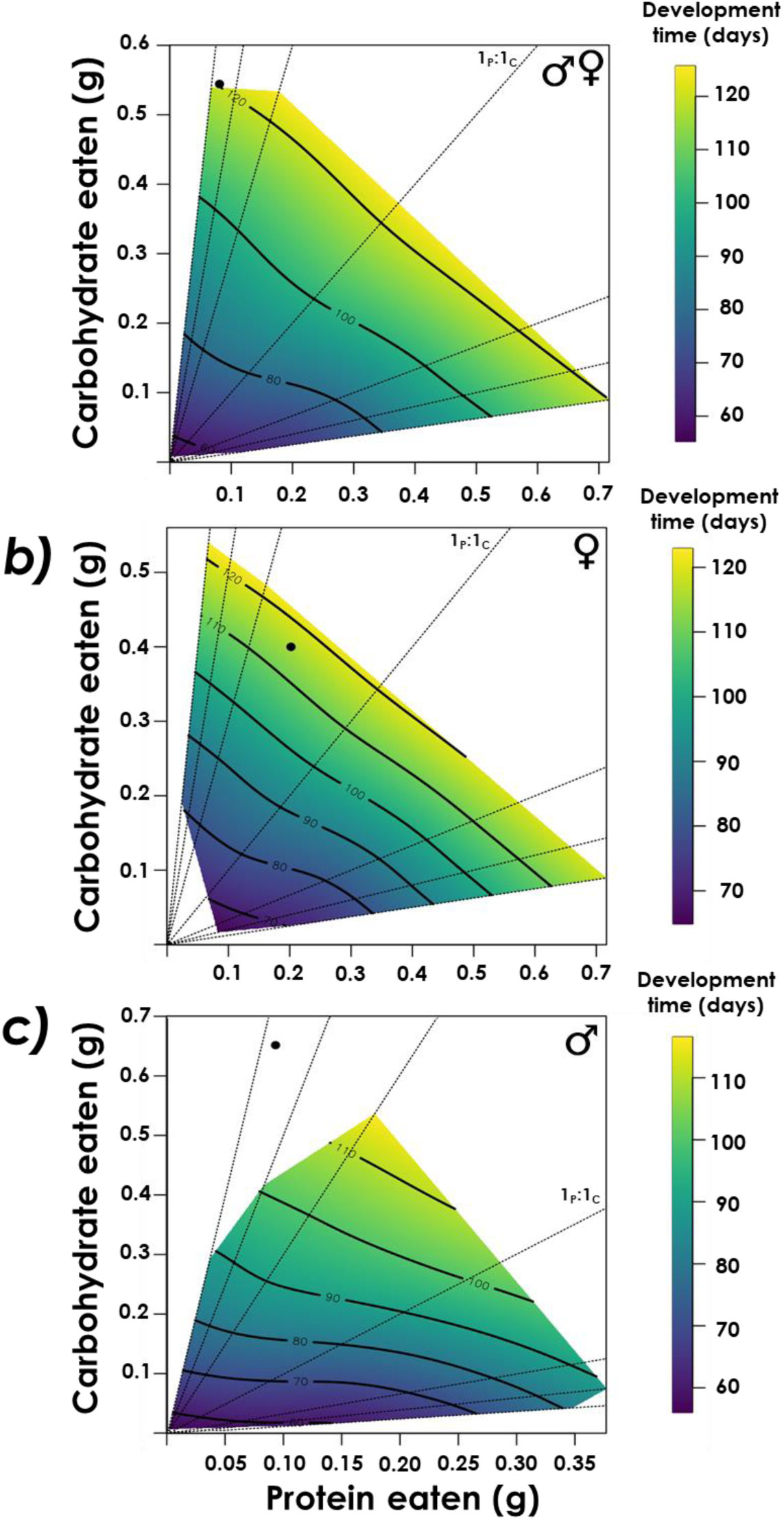
Response surfaces that illustrate the response of dietary protein and carbohydrate intake on pooled (a), female (b), and male (c) *Gryllodes sigillatus* development time and the corresponding global maximum (closed black circle). Global maxima represent undesirable macronutrient ratios, as longer development time potentially reduces yield. Dotted lines represent nutritional rails defined by the protein:carbohydrate ratios in the experimental diets. As crickets develop and feed on their individual diets, they move along their nutritional rails and age, represented by the colour gradient. The 1_P_:1_C_ nutritional rail is labelled on each panel.

The best approximating cox proportional hazard model that explained development time included P:C and cellulose concentration as the main predictors (AIC = 1478.50; Akaike weight = 1). Within the model, P:C had a significant effect across all cellulose concentrations (χ^2^ = 103.97; df = 2; *P* < 0.0001). Crickets fed diets equal in P:C or higher in carbohydrates were less likely to reach adulthood compared to crickets fed the 3_P_:1_C_ diet; crickets fed the 1_P_:8_C_, 1_P_:5_C_, 1_P_:3_C_, and 1_P_:1_C_ diets were 93%, 86%, 78%, and 64% less likely to develop into adults, respectively (*P* < 0.001; Figure 4; Table S6). Crickets fed 5_P_:1_C_ and 8_P_:1_C_ diets did not differ in the likelihood of developing to adults compared to crickets fed the 3_P_:1_C_ diet (*P* = 0.11, *P* = 0.28; Figure 4; Table S6). Cellulose significantly influenced the likelihood of developing to adulthood across all P:C diets (χ^2^ = 21.58; df = 6; *P* < 0.0001); crickets fed high cellulose diets were 63% less likely to develop to adulthood compared to crickets fed low cellulose diets (*P* < 0.001), but there was no difference between crickets fed medium and low cellulose diets (*P* = 0.74; Figure 4; Table S6).

**Figure 4.**
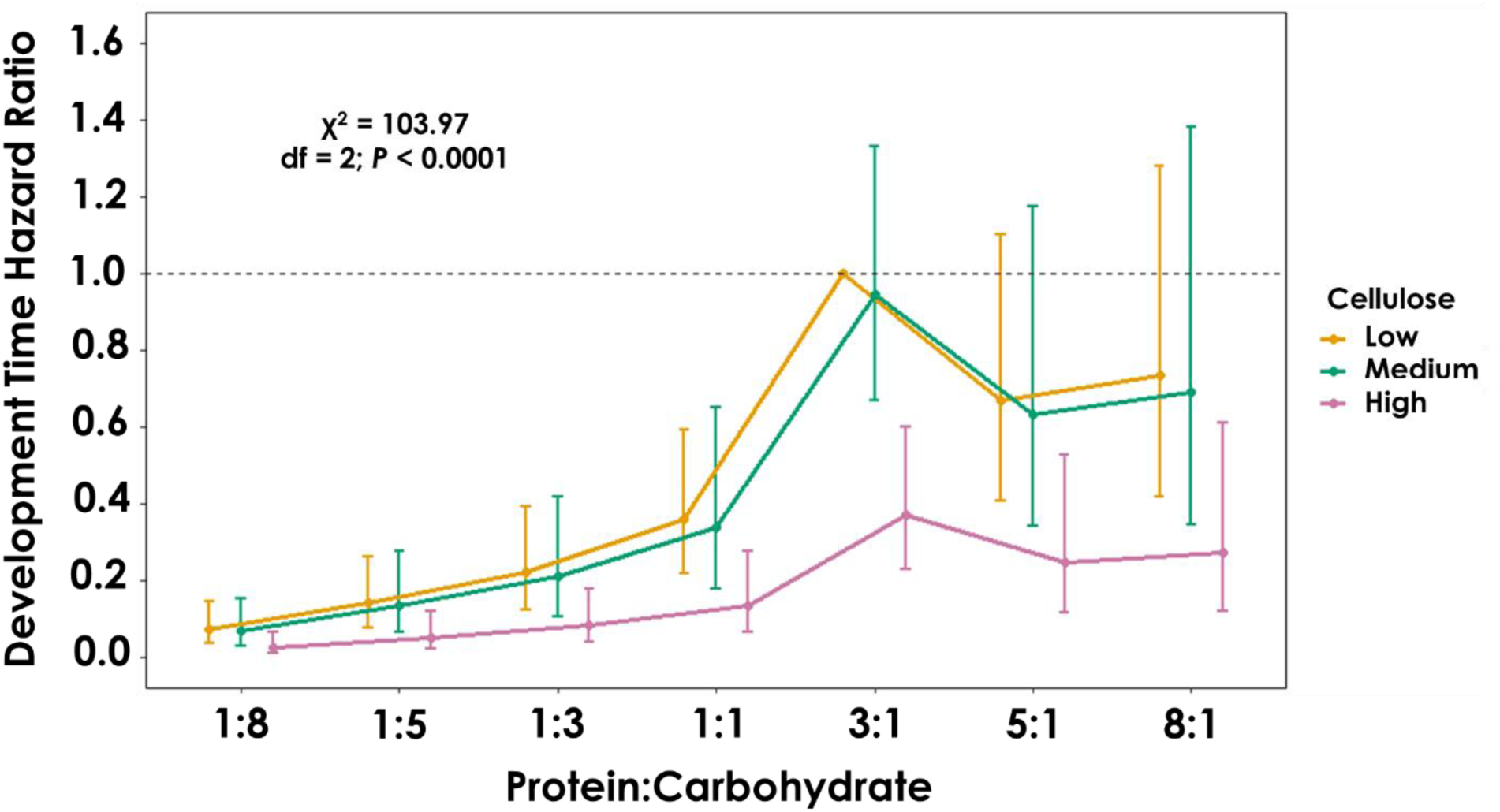
Cox proportional hazard ratios (± SE) for development time of *Gryllodes sigillatus* fed diets different in protein:carbohydrate (P:C) ratios and cellulose levels (Low = 14%, Medium = 45%, High = 76%). The dashed line represents the hazard ratio of the reference category (3_P_:1_C_, low cellulose). Coefficients and hazard ratio estimates may be found in Table S8.

The best approximating cox proportional hazard model that explained survival included P:C and time stratified cellulose as the main predictors (AIC = 7931.70; Akaike weight = 1). Within the model, P:C had a significant effect across all time stratified cellulose concentrations (χ^2^ = 16.12; df = 6; *P* = 0.013; Figure 5). Crickets fed the 8_P_:1_C_ diet had a 45% higher risk of death compared to crickets fed the 3_P_:1_C_ diet (Figure 5; Table S8). Time stratified cellulose also significantly influenced survival (χ^2^ = 106.92; df = 6; *P* < 0.0001). During the first seven days of life, crickets fed high (76%) and medium (45%) cellulose diets had 174% and 89% higher risk of death, respectively, compared to crickets fed low (14%) cellulose diets (Figure 5a; Table S8). During the next week of life (7-14 days), crickets fed high (76%) cellulose diets had a 505% higher risk of death compared to crickets fed low (14%) cellulose diets (Figure 5b; Table S8). There was no significant effect of cellulose concentration on survival during the third time interval (14-153 days; Figure 5c; Table S8).

**Figure 5.**
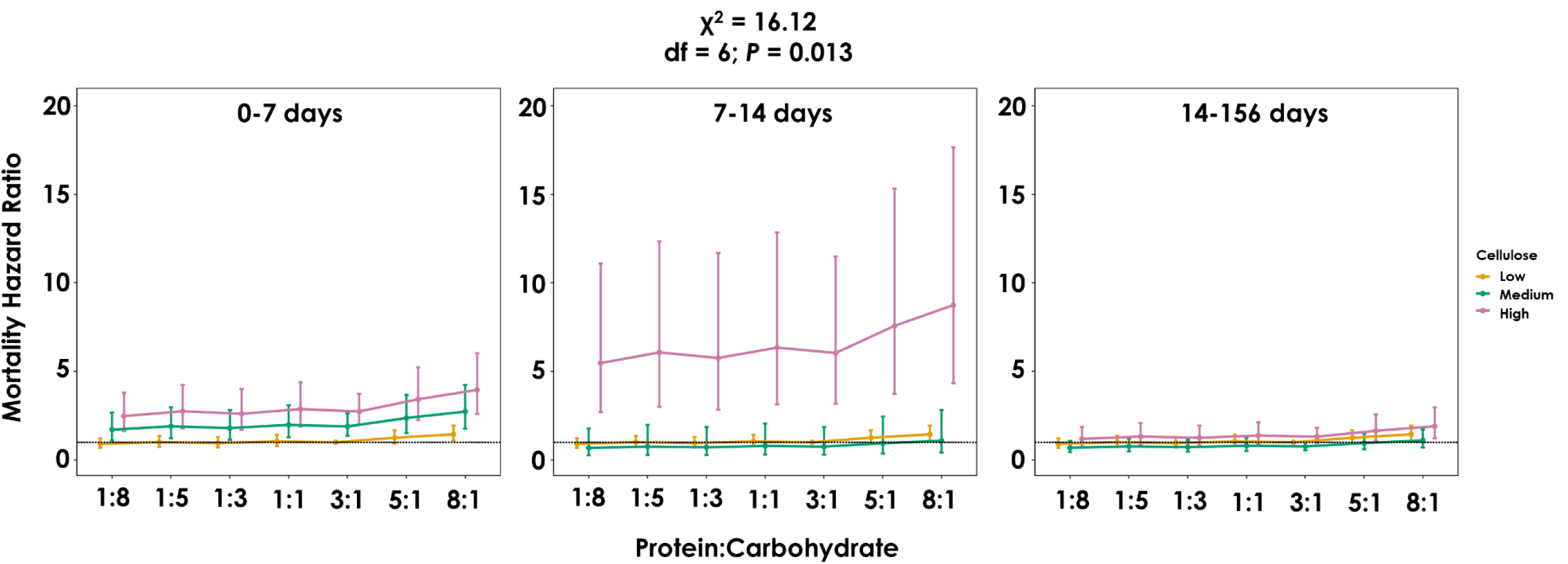
Cox proportional hazard ratios (± SE) for survival of *Gryllodes sigillatus* fed diets different in protein:carbohydrate (P:C) ratios and cellulose levels (Low = 14%, Medium = 45%, High = 76%). The dashed line represents the hazard ratio of the reference category (3_P_:1_C_, low cellulose). Coefficients and hazard ratio estimates may be found in Table S8.

Overall, the 3_P_:1_C_ diet resulted in the highest average mass, proportion surviving, and the shortest development time across the different diets (Table 1). As a result, the yield was the highest for the 3_P_:1_C_ diet whereas it was lowest for the 1_P_:8_C_ diet (Table 1).

**Table 1.** Average mass, proportion surviving, and days to adulthood (+/- standard error) of crickets fed one of seven diets different in protein:carbohydrate (P:C). Values are averaged across cellulose dilutions of 14%, 45%, and 76%. Yield metric is calculated as (Mass*Proportion surviving)/Days to adulthood).

| P:C | Mass (mg) ± SE | N adults | Proportion surviving ± SE | N deaths | Days to adulthood ± SE | Yield ± SE | 95% CI |
| --- | --- | --- | --- | --- | --- | --- | --- |
| 1:8 | 100.00 ± 9.47 | 11 | 0.15 ± 0.034 | 96 | 92.27 ± 2.07 | 0.16 ± 0.039 | 0.083 – 0.24 |
| 1:5 | 112.72 ± 6.32 | 16 | 0.17 ± 0.036 | 89 | 92.88 ± 2.02 | 0.20 ± 0.044 | 0.11 – 0.29 |
| 1:3 | 112.39 ± 8.26 | 17 | 0.18 ± 0.037 | 88 | 90.67 ± 1.94 | 0.22 ± 0.048 | 0.13 – 0.31 |
| 1:1 | 127.90 ± 5.92 | 30 | 0.23 ± 0.037 | 99 | 88.97 ± 1.70 | 0.33 ± 0.056 | 0.22 – 0.44 |
| 3:1 | 132.94 ± 6.89 | 37 | 0.31 ± 0.043 | 81 | 72.43 ± 1.50 | 0.58 ± 0.086 | 0.41 – 0.75 |
| 5:1 | 131.50 ± 7.31 | 29 | 0.22 ± 0.036 | 102 | 73.38 ± 1.51 | 0.40 ± 0.070 | 0.26 – 0.54 |
| 8:1 | 143.00 ± 8.72 | 20 | 0.16 ± 0.032 | 109 | 73.80 ± 1.49 | 0.30 ± 0.063 | 0.18 – 0.42 |

### 3.2 Choice of P:C

The best approximating model for diet choice did not include an interaction term between developmental time (week) and diet consumed (Table S10); therefore, dietary choice did not change throughout development. Week 1 body size (PC1) did not influence how much food crickets consumed (*P* = 0.11; Table S10). Both sexes ate more food over time, regardless of diet composition (*P* < 0.0001; Table S10). The observed pooled intake differed from expected; crickets selected a 1.05 _P_:1_C_ (t = −40.75; df = 51; *P* < 0.001). Males ate ∼17.7% more protein than females (*P* = 0.004; Figure 6; Table S10). Males’ observed P:C intake differed from expected; they selected a 1.38_P_:1_C_ (t = 2.95; df = 15; *P* = 0.01). Females’ observed 1_P_:1.15_C_ intake did not differ from the expected 1_P_:1_C_ (t = 2.00; df = 17; *P* = 0.062).

**Figure 6.**
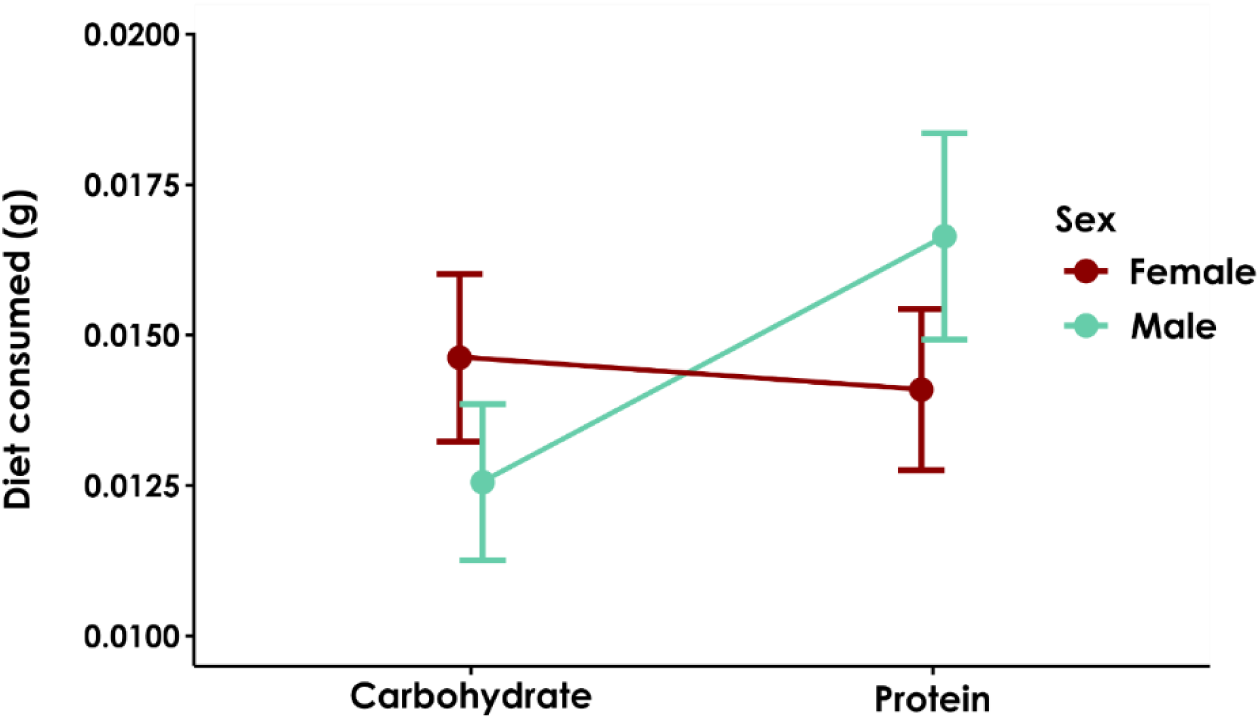
Amount of carbohydrate and protein consumed by individual female and male *Gryllodes sigillatus* when given a choice of 1_P_:0_C_ or 0_P_:1_C_ diet (45% cellulose dilution). There was a significant interaction between diet consumed and sex (*P* = 0.0017; Table S10).

## 4.0 Discussion

Our results add to a small, but growing, body of literature that indicates diets with proportionally more protein than carbohydrates fed from hatch to adulthood can maximize body size and mass of crickets while decreasing time to adulthood [40,57]. Here, adult *G. sigillatus* body size was maximized at 3.14_P_:1_C_ (Figure 1), mass was maximized at 3.22_P_:1_C_ (Figure 2), and crickets were more likely to develop to adulthood when fed a 3_P_:1_C_ diet (Figure 4). Females largely drove these optima toward high protein diets; both female mass and body size were significantly explained by protein intake and optimized around ∼3_P_:1_C_. Despite this, *G. sigillatus* males and females together selected a relatively balanced 1.05_P_:1_C_ diet when given a choice. Taking all the measured life history traits into account, our yield metric was generally higher for protein-skewed diets, with the 3_P_:1_C_ diet resulting in the highest yield -between 2.6 and 3.6 times higher than carbohydrate-skewed diets (Table 1). These results suggest that by optimizing the relative proportion of protein to carbohydrates up to a 3_P_:1_C_, diet can be leveraged as a tool in commercial settings to upscale cricket production by manipulating individuals to grow larger and heavier quickly. Notably, however, the lower 95% confidence interval of the 3_P_:1_C_ yield metric (0.41; Table 1) overlapped with the higher 95% confidence interval of the 1_P_:1_C_ yield metric (0.44; Table 1), which suggests that similar yields could be achieved using diet ratios that fall between 1_P_:1_C_ and 3_P_:1_C_, especially considering that crickets preferred to eat around the 1_P_:1_C_ ratio. An interesting follow-up experiment would be to test 3_P_:1_C_ and 1_P_:1_C_ diets at the colony-level to determine if overall yield differs.

When given a choice, male and female insects of the same species tend to consume different P:C. Female fitness traits such as egg production and fecundity are typically maximized on higher P:C, while male fitness traits such as mate signalling behaviour are typically maximized on lower P:C that favour carbohydrate availability [29,37,41,52–54]. The developmental intake target we report here for *G. sigillatus* (1.05_P_:1_C_) is different from the adult intake targets reported by Rapkin et al. (1_P_:1.74_C_ – 1_P_:2_C_; Rapkin et al., 2016; Rapkin et al., 2018). A major difference between our experiment and those mentioned is that we measured diet choice from the first week of life until adulthood, compared to adult feeding for 10-16 days [41,52]. Regardless, dietary preference did not change over time in our choice experiment, and so the intake target between juvenile and adult stages was not different. Females in our experiment selected a 1_P_:1.15_C_, an intake target relatively close to the 1_P_:1.84_C_ previously reported (Rapkin et al. 2018). However, males preferred slightly more protein than carbohydrates (1.38_P_:1_C_), a stark difference from the previously reported 1_P_:1.74_C_ and 1_P_:2_C_ for male *G. sigillatus* [41,52] and from the 1_P_:4.1_C_ preference recorded for *G. veletis* [29]. Sexual conflict in *G. sigillatus* is regulated by the relative intake of protein and carbohydrates; spermatophylax (an endogenous nuptial gift made by male crickets) traits are maximized at a balanced 1_P_:1.3_C_, but males demonstrate an intake target of 1_P_:1.74_C_ (Rapkin et al. 2016). It is hypothesized that crickets producing an attractive spermatophylax have a high demand for protein, driving the male optimum to a higher amount of protein compared to the intake target (Rapkin et al. 2016). We suggest that this male preference for protein over carbohydrates could be a result of selection pressures that may occur in a mass rearing environment leading to population-specific, or ‘strain’ differences, that are ultimately driving the male intake target towards more protein for spermatophylax production. Strain-specific differences among mass reared insect species have been recently reported, including differences in growth, development time, feed conversion efficiency, nutritional composition, and bacterial communities [11,77–79]. We used eggs from a commercial supplier who has been rearing this species for ∼9 years, while Rapkin et al. have been maintaining a wild-caught colony since 2001. In a mass reared insect facility, billions of crickets are competing with one another in closely confined spaces, and harvest schedules likely exert strong selection pressures on the population. Individuals that consume more carbohydrates, and thus take a longer time to develop to adulthood, may be quickly selected out by the harvest schedule. Further, in such an environment, it may be impossible for females to orient themselves to a calling male. Thus, the selection pressures on calling behaviour, a trait driven by increased carbohydrate consumption, may be greatly decreased. Instead, males that mature and achieve a large size quickly (by consuming more protein) may have an advantage over males that invest more in calling (by consuming more carbohydrate), potentially explaining why males in our experiment preferred to eat higher protein diets. Measuring the sexually selected traits of a farm population such as spermatophylax weight, male calling behaviour, and female fecundity is one method to test this possible avenue of nutrient regulated sexual conflict. Comparisons with wild-caught populations would help demonstrate the evolution of strain-specific differences. To our knowledge, however, no comparable data exist on wild-caught *G. sigillatus*.

Individual-level experiments such as ours that measure the dietary responses to a wide range of nutrient compositions are a crucial first step for developing the nutritional ‘story’ behind an organism. For our results to have direct impact for insect farmers and mass producers, it is critically important to apply the nutritional ecology lessons we learn at the individual level to the colony level. This begins with a single generation experiment to determine if results at the individual level are consistent across group-level trials, and then followed by large-scale multigenerational experiments. Generational effects of diet can influence feeding preferences and offspring life history responses important to yield [80]. Survival, development, and pupal mass of *Plutella xylostella* reared for 350 generations were highest for individuals that were fed the same P:C diet of the ancestral culture [80]. A recent study group-reared *G. sigillatus* for 37-46 generations and revealed compensatory feeding occurred when fed low nutrient diets [81]. Similarly, black soldier fly larva (*Hermetia illucens*) reared for 13 generations adapted to a low-quality diet and grew heavier larva and pupa [82]. To perform multigenerational diet experiments requires time, space, and resources, factors that are species-dependent; the time, space, and resources to rear thousands of *D. melanogaster* for 100 generations is much less compared to rearing thousands of *G. sigillatus* for 100 generations. For nutritional ecology research to make rapid progress, collaboration among researchers performing long-term multigenerational experiments is encouraged so that multiple questions and hypotheses can be asked and applied to insect colonies.

Our results demonstrate that while it is possible for crickets to grow on scant resources, cellulose dilution of diet ingredients is extremely stressful. Highly diluted diets (76% cellulose) had a very low proportion of survivors, with the proportion of crickets surviving fed these diets lower than 0.10 except for the 1_P_:1_C_ (0.11 proportion surviving) and 3_P_:1_C_ (0.15 proportion surviving) diets (Table S4). This necessitates revisiting the holidic diet recipe that has been used to develop foundational insect nutritional geometry research [34]. It would be advantageous if a new recipe was created that could be successfully fed to hatchlings and juvenile insects without the risk of early mortality, or alternatively studies that use holidic diets are also paired with oligidic diet studies. This would allow for more careful research into the lifetime influence of diet on adult life history traits and could help to paint a more complete picture of how diet influences organismal life history throughout their entire lives. An additional challenge is designing homogenous oligidic diets in specific P:C ratios that restrict the selection of ingredients by the crickets. Currently, most commercial cricket feeds are heterogeneous in design; different ingredients are ground separately prior to mixing, and this results in a high distribution of particles sizes among ingredients. Our results suggest that *G. sigillatus* does not select a diet that optimizes life history traits important to increasing commercial yield (ex. mass, body size, development time). Therefore, yield of *G. sigillatus* could be optimized by finely grinding and then pelleting or kibbling the diet to restrict dietary choice. However, females selected a diet slightly higher in carbohydrates (1_P_:1.15_C_) compared to males (1.38_P_:1_C_), and there may be an important female life history trait that we did not measure in this experiment that relies on carbohydrate consumption, such as an energy-intensive behaviour like egg laying. Prior to restricting dietary choice of both males and females to a higher protein diet, multigenerational experiments that feed holidic diets to individually reared crickets and their offspring are required to further tease out the fitness effects of manipulating protein and carbohydrate availability. Measurements of reproduction success, offspring viability, and offspring survival would contribute to a robust measurement of individual fitness for each sex.

Overall, we have presented a comprehensive approach to examining the effects of a wide range of dietary P:C across three different nutrient dilutions from hatch to adulthood of an important commercially reared cricket species. Our results clearly suggest that a 3_P_:1_C_ diet optimizes *G. sigillatus* life history traits important to production yield, but individuals prefer to selectively feed at a balanced 1.05_P_:1_C._

## Supporting information

Electronic Supplemental Tables and Figures

All Data

## Acknowledgements

The authors thank Michelle Léveillée for laboratory assistance and Entomo Farms for providing materials, specimens, and diet. This work was supported by a MITACS Accelerate Internship, funding from the Province of Ontario, Ontario Graduate Scholarships awarded to M.J.M, NSERC Discovery Grants awarded to both H.A.M. and S.M.B., and the Canadian Foundation for Innovation. M.J.M., E.R.M., H.A.M., and S.M.B conceived the experiments and contributed to writing the manuscript. M.J.M., C.T.B., H.B., S.M., E.R.M., and C.C.S. conducted the research. M.J.M., S.J.H., and S.M.B. analysed the data.

