## Supplementary material for "Applying nutritional ecology to optimize diets of crickets raised for food and feed": Electronic Supplemental Tables and Figures

**Table S1.** Censoring data for cox proportional hazards development time and survival time models. The positive status (adult and death) are represented by 1, and the negative status (not adult and alive) are represented by 0.

| | | Development Time Model | | Survival Time Model | |
|---|---|---|---|---|---|
| **Number of Individuals** | **Category** | **Development Time** | **Status = Adult** | **Survival Time** | **Status = Death** |
| 8 | Died naturally as adult | day they eclosed as adult | 1 | day they died | 1 |
| 153 | Euthanized as adult | day they eclosed as adult | 1 | day they eclosed as adult + 7 days | 0 |
| 656 | Died as juveniles | day they died | 0 | day they died | 1 |
| 10 | Experiment ended and never became an adult | day they were euthanized | 0 | day they were euthanized | 0 |
| 101 | Lost - Went missing as juveniles OR death date not recorded | last day seen in the experiment | 0 | last day seen in the experiment | 0 |

**Table S2.** Model results from general linear models of how nutrient intake influenced life history traits of crickets. B-Y p = 0.02. Summary statistics from these models are found in Table A.2.1.

| Model | $R^2$ | Term | Estimate (±SE) | t-value | P-value |
|---|---|---|---|---|---|
| Body size (pooled) | 0.43 | **Intercept** | **-4.19 ± 0.54** | **-7.82** | **<0.001** |
| | | **P** | **21.86 ± 3.32** | **6.58** | **<0.001** |
| | | **C** | **14.48 ± 3.63** | **3.99** | **<0.001** |
| | | **$P^2$** | **-19.54 ± 4.87** | **-4.02** | **<0.001** |
| | | $C^2$ | -13.17 ± 5.97 | -2.21 | 0.029 |
| | | P*C | -18.44 ± 13.38 | -1.38 | 0.17 |
| Body size (females) | 0.46 | **Intercept** | **-1.69 ± 0.56** | **-3.04** | **0.0031** |
| | | **P** | **16.51 ± 3.05** | **5.41** | **<0.001** |
| | | C | 4.05 ± 3.52 | 1.15 | 0.25 |
| | | **$P^2$** | **-17.70 ± 3.78** | **-4.68** | **<0.001** |
| | | $C^2$ | -0.90 ± 5.27 | -0.17 | 0.87 |
| | | P*C | -5.12 ± 11.19 | -0.46 | 0.65 |
| Body size (males) | 0.42 | **Intercept** | **-3.29 ± 0.59** | **-5.60** | **<0.001** |
| | | P | 6.16 ± 5.50 | 1.12 | 0.27 |
| | | C | 6.59 ± 3.66 | 1.80 | 0.077 |
| | | $P^2$ | 15.04 ± 13.18 | 1.14 | 0.26 |
| | | $C^2$ | 2.76 ± 8.26 | 0.33 | 0.74 |
| | | P*C | -26.83 ± 21.00 | -1.28 | 0.21 |
| Mass (pooled) | 0.45 | **Intercept** | **42.28 ± 11.71** | **3.61** | **<0.001** |
| | | **P** | **466.18 ± 72.62** | **6.42** | **<0.001** |
| | | **C** | **240.81 ± 79.25** | **3.04** | **0.0028** |
| | | **$P^2$** | **-400.07 ± 106.38** | **-3.76** | **<0.001** |
| | | $C^2$ | -257.97 ± 130.57 | -1.98 | 0.050 |
| | | P*C | -247.05 ± 292.50 | -0.85 | 0.40 |
| Mass (females) | 0.43 | **Intercept** | **68.38 ± 18.18** | **3.76** | **<0.001** |
| | | **P** | **441.55 ± 99.78** | **4.43** | **<0.001** |
| | | C | 108.41 ± 115.09 | 0.94 | 0.35 |
| | | **$P^2$** | **-443.92 ± 123.53** | **-3.59** | **<0.001** |

| | | | | | |
|---|---|---|---|---|---|
| | | $C^2$ | -95.48 ± 172.09 | -0.56 | 0.58 |
| | | P*C | -27.68 ± 365.83 | -0.076 | 0.94 |
| Mass (males) | 0.37 | **Intercept** | **59.97 ± 14.45** | **4.15** | **<0.001** |
| | | P | 227.93 ± 135.19 | 1.69 | 0.097 |
| | | C | 120.14 ± 89.88 | 1.34 | 0.19 |
| | | $P^2$ | 174.47 ± 324.04 | 0.54 | 0.59 |
| | | $C^2$ | 133.31 ± 203.10 | 0.66 | 0.51 |
| | | P*C | -952.6 ± 516.19 | -1.85 | 0.070 |
| Development time (pooled) | 0.45 | **Intercept** | **57.82 ± 6.50** | **8.89** | **<0.001** |
| | | P | -2.00 ± 40.36 | -0.05 | 0.96 |
| | | **C** | **142.71 ± 43.88** | **3.25** | **0.0014** |
| | | $P^2$ | 100.24 ± 59.13 | 1.70 | 0.092 |
| | | $C^2$ | -98.66 ± 72.38 | -1.36 | 0.18 |
| | | P*C | 162.76 ± 162.61 | 1.00 | 0.32 |
| Development time (females) | 0.41 | **Intercept** | **72.42 ± 11.06** | **6.55** | **<0.001** |
| | | P | -49.95 ± 60.78 | -0.82 | 0.41 |
| | | C | 44.24 ± 69.69 | 0.64 | 0.53 |
| | | $P^2$ | 129.46 ± 75.26 | 1.72 | 0.089 |
| | | $C^2$ | 50.34 ± 104.24 | 0.48 | 0.63 |
| | | P*C | 335.89 ± 222.75 | 1.51 | 0.14 |
| Development time (males) | 0.42 | **Intercept** | **59.35 ± 9.50** | **6.25** | **<0.001** |
| | | P | -98.81 ± 88.85 | -1.11 | 0.27 |
| | | **C** | **180.94 ± 59.08** | **3.06** | **0.0033** |
| | | $P^2$ | 333.60 ± 212.98 | 1.57 | 0.12 |
| | | $C^2$ | -309.55 ± 133.49 | -2.32 | 0.024 |
| | | P*C | 516.70 ± 339.27 | 1.52 | 0.13 |

**Table S3.** Model results from Type II analyses of variance associated with the linear models of how nutrient intake influenced life history traits of crickets. B-Y p = 0.02. The overall model summary statistics are presented in the first row for each model, followed by the term statistics in the next five rows.

| Model | $R^2$ | Term | df | F-value | P-value |
|---|---|---|---|---|---|
| Body size (pooled) | 0.43 | | **5,154** | **24.48** | **< 0.001** |
| | | **P** | **1,154** | **53.72** | **<0.001** |
| | | **C** | **1,154** | **16.10** | **<0.001** |
| | | **$P^2$** | **1,154** | **16.12** | **<0.001** |
| | | $C^2$ | 1,154 | 4.87 | 0.029 |
| | | P*C | 1,154 | 1.90 | 0.17 |
| Body size (females) | 0.46 | | **5,87** | **16.40** | **< 0.001** |
| | | **P** | **1,87** | **52.39** | **<0.001** |
| | | C | 1,87 | 1.46 | 0.23 |
| | | **$P^2$** | **1,87** | **21.94** | **<0.001** |
| | | $C^2$ | 1,87 | 0.029 | 0.87 |
| | | P*C | 1,87 | 0.21 | 0.65 |
| Body size (males) | 0.42 | | **5,61** | **10.46** | **< 0.001** |
| | | P | 1,61 | 0.34 | 0.56 |
| | | C | 1,61 | 2.14 | 0.15 |
| | | $P^2$ | 1,61 | 1.30 | 0.26 |
| | | $C^2$ | 1,61 | 0.11 | 0.74 |
| | | P*C | 1,61 | 1.63 | 0.21 |
| Mass (pooled) | 0.45 | | **5,154** | **26.57** | **< 0.001** |
| | | **P** | **1,154** | **56.99** | **<0.001** |
| | | **C** | **1,154** | **10.35** | **0.0016** |
| | | **$P^2$** | **1,154** | **14.14** | **<0.001** |
| | | $C^2$ | 1,154 | 3.90 | 0.050 |
| | | P*C | 1,154 | 0.71 | 0.40 |
| Mass (females) | 0.43 | | **5,87** | **14.80** | **< 0.001** |
| | | **P** | **1,87** | **38.75** | **<0.001** |
| | | C | 1,87 | 1.73 | 0.19 |

| | | | | | |
|---|---|---|---|---|---|
| | | **$P^2$** | **1,87** | **12.91** | **<0.001** |
| | | $C^2$ | 1,87 | 0.31 | 0.58 |
| | | P*C | 1,87 | 0.0057 | 0.94 |
| Mass (males) | 0.37 | | **5,61** | **8.75** | **< 0.001** |
| | | P | 1,61 | 0.85 | 0.36 |
| | | C | 1,61 | 0.60 | 0.44 |
| | | $P^2$ | 1,61 | 0.29 | 0.59 |
| | | $C^2$ | 1,61 | 0.43 | 0.51 |
| | | P*C | 1,61 | 3.41 | 0.070 |
| Development time (pooled) | 0.45 | | **5,155** | **27.22** | **< 0.001** |
| | | P | 1,155 | 0.54 | 0.46 |
| | | **C** | **1,155** | **24.74** | **<0.001** |
| | | $P^2$ | 1,155 | 2.87 | 0.092 |
| | | $C^2$ | 1,155 | 1.86 | 0.18 |
| | | P*C | 1,155 | 1.00 | 0.32 |
| Development time (females) | 0.41 | | **5,88** | **13.86** | **< 0.001** |
| | | P | 1,88 | 0.13 | 0.72 |
| | | **C** | **1,88** | **6.78** | **0.011** |
| | | $P^2$ | 1,88 | 2.96 | 0.089 |
| | | $C^2$ | 1,88 | 0.23 | 0.63 |
| | | P*C | 1,88 | 2.27 | 0.14 |
| Development time (males) | 0.42 | | **5,61** | **10.56** | **< 0.001** |
| | | P | 1,61 | 0.20 | 0.66 |
| | | **C** | **1,61** | **14.26** | **<0.001** |
| | | $P^2$ | 1,61 | 2.45 | 0.12 |
| | | $C^2$ | 1,61 | 5.38 | 0.024 |
| | | P*C | 1,61 | 2.32 | 0.13 |

**Table S4**. Average mass, proportion surviving, and average days to adulthood (± standard error) of crickets fed one of seven diets different in protein:carbohydrate (P:C) and cellulose.

| P:C | Cellulose | Adult mass (mg) | *N* adults | Proportion survived | *N* deaths | Days to adulthood |
|---|---|---|---|---|---|---|
| 1:8 | 14% | 102.65 ± 12.88 | 4 | 0.21 ± 0.078 | 33 | 93.50 ± 6.21 |
| 1:8 | 45% | 106.48 ± 13.09 | 6 | 0.24 ± 0.071 | 44 | 88.83 ± 2.77 |
| 1:8 | 76% | 50.54 ± 0.00 | 1 | 0.042 ± 0.029 | 59 | 108.00 ± 0.00 |
| 1:5 | 14% | 117.07 ± 11.19 | 5 | 0.24 ± 0.085 | 26 | 95.00 ± 5.00 |
| 1:5 | 45% | 113.42 ± 8.26 | 10 | 0.32 ± 0.080 | 34 | 96.00 ± 2.94 |
| 1:5 | 76% | 83.85 ± 0.00 | 1 | 0.021 ± 0.021 | 56 | 51.00 ± 0.00 |
| 1:3 | 14% | 129.43 ± 15.91 | 6 | 0.23 ± 0.083 | 27 | 87.86 ± 4.31 |
| 1:3 | 45% | 102.29 ± 10.07 | 9 | 0.27 ± 0.078 | 38 | 88.22 ± 2.89 |
| 1:3 | 76% | 106.70 ± 21.77 | 2 | 0.083 ± 0.040 | 50 | 111.50 ± 4.10 |
| 1:1 | 14% | 123.67 ± 8.98 | 10 | 0.33 ± 0.086 | 33 | 85.40 ± 2.81 |
| 1:1 | 45% | 133.44 ± 10.68 | 13 | 0.39 ± 0.085 | 36 | 85.39 ± 3.92 |
| 1:1 | 76% | 123.65 ± 10.57 | 7 | 0.11 ± 0.038 | 73 | 100.72 ± 1.96 |
| 3:1 | 14% | 123.77 ± 8.97 | 14 | 0.42 ± 0.086 | 34 | 63.57 ± 2.20 |
| 3:1 | 45% | 148.93 ± 12.92 | 15 | 0.46 ± 0.087 | 37 | 74.00 ± 2.83 |
| 3:1 | 76% | 118.99 ± 11.07 | 8 | 0.15 ± 0.050 | 63 | 85.00 ± 2.23 |
| 5:1 | 14% | 128.13 ± 8.59 | 15 | 0.50 ± 0.091 | 33 | 70.60 ± 3.14 |
| 5:1 | 45% | 146.69 ± 15.52 | 10 | 0.26 ± 0.070 | 43 | 77.00 ± 3.18 |
| 5:1 | 76% | 106.18 ± 9.89 | 4 | 0.065 ± 0.031 | 67 | 74.75 ± 1.32 |
| 8:1 | 14% | 142.79 ± 15.18 | 8 | 0.28 ± 0.083 | 30 | 72.88 ± 4.10 |
| 8:1 | 45% | 150.47 ± 11.90 | 10 | 0.21 ± 0.060 | 52 | 75.10 ± 2.19 |
| 8:1 | 76% | 106.48 ± 3.62 | 2 | 0.038 ± 0.026 | 60 | 71.00 ± 1.83 |

**Table S5.** Penultimate confidence set of Cox proportional hazard models examining which predictors (P:C and cellulose) and their interaction best explained development time, showing only the subset candidate models with Δ AIC < 4.

| **Model** | **AIC** | **Δ AIC [a]** | **Weight [b]** | **Evidence Ratio [c]** | **Included In Confidence Set [d]** |
|---|---|---|---|---|---|
| **P:C + cellulose** | **1478.50** | **0** | **0.803** | | **Yes** |
| P:C + cellulose + P:C*cellulose | 1481.30 | 2.81 | 0.197 | 4.08 | No[d] |

[a] The difference between the current model's AIC score and that of the top ranked model (bolded). Models with AIC > 4 have been omitted from this table

[b] Akaike weights, or relative likelihood of the current model. Calculated as exp(-0.5 x Δ AIC) divided by the sum of these values across all models in the confidence set.

[c] Evidence ratio, or how many times more parsimonious the top ranked model (bolded) is over the current model. Calculated as the Akaike weight of the top ranked model divided by that of the current model.

[d] Models were excluded from the confidence model set if they were more complex variants of any model with a lower AIC value (nesting approach; Richards 2008, Richards et al 2011).

**Table S6.** Coefficient and hazard ratio (HR) estimates and the 95% confidence intervals for the top ranked (lowest AIC) model showing how predictors (P:C and cellulose) influenced developmental time. Reference categories: P:C = 3:1, cellulose 14%.

| **Model Term** | **B** | **SE(β)** | **Hazard Ratio (HR) (95% CI for HR)** | **z** | ***P*** |
|---|---|---|---|---|---|
| Cellulose 45% | -0.057 | 0.18 | 0.94 (0.67 – 1.33) | -0.33 | 0.74 |
| **Cellulose 76%** | **-0.99** | **0.25** | **0.37 (0.23 – 0.60)** | **-4.04** | **<0.001** |
| **P:C 1:8** | **-2.64** | **0.36** | **0.071 (0.04 – 0.14)** | **-7.37** | **<0.001** |
| **P:C 1:5** | **-1.96** | **0.31** | **0.14 (0.076 – 0.26)** | **-6.27** | **<0.001** |
| **P:C 1:3** | **-1.51** | **0.29** | **0.22 (0.13 – 0.39)** | **-5.14** | **<0.001** |
| **P:C 1:1** | **-1.02** | **0.26** | **0.36 (0.22 – 0.59)** | **-4.02** | **<0.001** |
| P:C 5:1 | -0.40 | 0.25 | 0.67 (0.41 – 1.10) | -1.58 | 0.11 |
| P:C 8:1 | -0.31 | 0.29 | 0.73 (0.42 – 1.28) | -1.09 | 0.28 |

**Table S7.** Penultimate confidence set of Cox proportional hazard models examining which predictors (P:C and cellulose) best explained survival, showing only the subset candidate models with $\Delta$ AIC < 4.

| **Model** | **AIC** | **$\Delta$ AIC [a]** | **Weight [b]** | **Evidence Ratio [c]** | **Included In** | **Confidence Set [d]** |
|---|---|---|---|---|---|---|
| **P:C + Cellulose** | 7931.70 | 0 | 0.92 | | | **Yes** |

[a] The difference between the current model's AIC score and that of the top ranked model (bolded). Models with AIC > 4 have been omitted from this table

[b] Akaike weights, or relative likelihood of the current model. Calculated as exp(-0.5 x $\Delta$ AIC) divided by the sum of these values across all models in the confidence set.

[c] Evidence ratio, or how many times more parsimonious the top ranked model (bolded) is over the current model. Calculated as the Akaike weight of the top ranked model divided by that of the current model.

[d] Models were excluded from the confidence model set if they were more complex variants of any model with a lower AIC value (nesting approach; Richards 2008, Richards et al 2011).

**Table S8.** Coefficient and hazard ratio (HR) estimates for the top ranked (lowest AIC) model showing how predictors (P:C and time stratified cellulose) influenced survival. Reference categories: P:C = 3:1, cellulose 14%. Time interval 1: 0-7 days, time interval 2: 7-14 days, time interval 3: 14-153 days.

| **Model Term** | **B** | **SE(β)** | **Hazard Ratio (HR) (95% CI for HR)** | **z** | ***P*** |
|---|---|---|---|---|---|
| P:C 1:8 | -0.10 | 0.15 | 0.91 (0.67 – 1.22) | -0.65 | 0.51 |
| P:C 1:5 | 0.0050 | 0.16 | 1.01 (0.74 – 1.36) | 0.029 | 0.98 |
| P:C 1:3 | -0.049 | 0.16 | 0.95 (0.70 – 1.29) | -0.31 | 0.75 |
| P:C 1:1 | 0.048 | 0.15 | 1.05 (0.78 – 1.41) | 0.32 | 0.75 |
| P:C 5:1 | 0.23 | 0.15 | 1.25 (0.94 – 1.68) | 1.51 | 0.13 |
| **P:C 8:1** | **0.37** | **0.15** | **1.45 (1.08 – 1.93)** | **2.51** | **0.012** |
| **Cellulose 76%*Time Interval 1** | **1.01** | **0.16** | **2.74 (2.00 – 3.73)** | **6.39** | **<0.0001** |
| **Cellulose 45%*Time Interval 1** | **0.64** | **0.17** | **1.89 (1.35 – 2.64)** | **3.72** | **<0.001** |
| **Cellulose 76%*Time Interval 2** | **1.80** | **0.33** | **6.05 (3.18 – 11.48)** | **5.50** | **<0.0001** |
| Cellulose 45%*Time Interval 2 | -0.29 | 0.47 | 0.75 (0.30 – 1.87) | -0.62 | 0.54 |
| Cellulose 76%*Time Interval 3 | 0.27 | 0.17 | 1.31 (0.95 – 1.82) | 1.65 | 0.10 |
| Cellulose 45%*Time Interval 3 | -0.27 | 0.17 | 0.76 (0.54 – 1.07) | -1.58 | 0.11 |

**Table S9.** Hazard ratios (HR) and associated percent risk of death with increasing 1 standard deviation (SD) in P:C for the 63 diet categories. HR that differ significantly from 1 are bolded.

| Category | P:C | Cellulose | Time Interval | HR | % Risk of Death[a] | Lower 95% CI | Upper 95% CI |
|---|---|---|---|---|---|---|---|
| **1** | **3:1** | **76%** | **1** | **2.74** | **173.60** | **2.01** | **3.73** |
| **2** | **1:8** | **76%** | **1** | **2.48** | **147.60** | **1.62** | **3.80** |
| **3** | **1:5** | **76%** | **1** | **2.75** | **174.80** | **1.79** | **4.23** |
| **4** | **1:3** | **76%** | **1** | **2.61** | **160.60** | **1.69** | **4.01** |
| **5** | **1:1** | **76%** | **1** | **2.87** | **187.00** | **1.88** | **4.39** |
| **6** | **5:1** | **76%** | **1** | **3.43** | **242.80** | **2.25** | **5.23** |
| **7** | **8:1** | **76%** | **1** | **3.96** | **295.70** | **2.60** | **6.03** |
| **8** | **3:1** | **45%** | **1** | **1.89** | **88.90** | **1.35** | **2.64** |
| **9** | **1:8** | **45%** | **1** | **1.71** | **70.90** | **1.10** | **2.67** |
| **10** | **1:5** | **45%** | **1** | **1.90** | **89.70** | **1.21** | **2.97** |
| **11** | **1:3** | **45%** | **1** | **1.80** | **79.90** | **1.15** | **2.82** |
| **12** | **1:1** | **45%** | **1** | **1.98** | **98.10** | **1.27** | **3.09** |
| **13** | **5:1** | **45%** | **1** | **2.37** | **136.60** | **1.52** | **3.67** |
| **14** | **8:1** | **45%** | **1** | **2.73** | **173.10** | **1.77** | **4.22** |
| 15 | 3:1 | 14% | 1 | 1.00 | 0.00 | 1.00 | 1.00 |
| 16 | 1:8 | 14% | 1 | 0.91 | -9.50 | 0.67 | 1.22 |
| 17 | 1:5 | 14% | 1 | 1.00 | 0.40 | 0.74 | 1.36 |
| 18 | 1:3 | 14% | 1 | 0.95 | -4.70 | 0.70 | 1.29 |
| 19 | 1:1 | 14% | 1 | 1.05 | 4.90 | 0.78 | 1.41 |
| 20 | 5:1 | 14% | 1 | 1.25 | 25.30 | 0.94 | 1.68 |
| **21** | **8:1** | **14%** | **1** | **1.45** | **44.60** | **1.08** | **1.93** |
| **22** | **3:1** | **76%** | **2** | **6.05** | **504.50** | **3.18** | **11.48** |
| **23** | **1:8** | **76%** | **2** | **5.47** | **447.20** | **2.70** | **11.09** |
| **24** | **1:5** | **76%** | **2** | **6.07** | **507.20** | **2.99** | **12.35** |
| **25** | **1:3** | **76%** | **2** | **5.76** | **475.80** | **2.84** | **11.69** |
| **26** | **1:1** | **76%** | **2** | **6.34** | **534.10** | **3.13** | **12.85** |
| **27** | **5:1** | **76%** | **2** | **7.57** | **657.30** | **3.74** | **15.33** |
| **28** | **8:1** | **76%** | **2** | **8.74** | **774.30** | **4.33** | **17.67** |

| | | | | | | | |
|---|---|---|---|---|---|---|---|
| 29 | 3:1 | 45% | 2 | 0.75 | -24.90 | 0.30 | 1.87 |
| 30 | 1:8 | 45% | 2 | 0.68 | -32.10 | 0.26 | 1.77 |
| 31 | 1:5 | 45% | 2 | 0.75 | -24.60 | 0.29 | 1.97 |
| 32 | 1:3 | 45% | 2 | 0.72 | -28.50 | 0.27 | 1.87 |
| 33 | 1:1 | 45% | 2 | 0.79 | -21.30 | 0.30 | 2.05 |
| 34 | 5:1 | 45% | 2 | 0.94 | -6.00 | 0.36 | 2.45 |
| 35 | 8:1 | 45% | 2 | 1.09 | 8.60 | 0.42 | 2.82 |
| 36 | 3:1 | 14% | 2 | 1.00 | 0.00 | 1.00 | 1 |
| 37 | 1:8 | 14% | 2 | 0.91 | -9.50 | 0.67 | 1.22 |
| 38 | 1:5 | 14% | 2 | 1.00 | 0.40 | 0.74 | 1.36 |
| 39 | 1:3 | 14% | 2 | 0.95 | -4.70 | 0.70 | 1.29 |
| 40 | 1:1 | 14% | 2 | 1.05 | 4.90 | 0.78 | 1.41 |
| 41 | 5:1 | 14% | 2 | 1.25 | 25.30 | 0.94 | 1.68 |
| **42** | **8:1** | **14%** | **2** | **1.45** | **44.60** | **1.08** | **1.93** |
| **43** | **3:1** | **76%** | **3** | **1.31** | **31.30** | **0.95** | **1.81** |
| 44 | 1:8 | 76% | 3 | 1.19 | 18.80 | 0.76 | 1.86 |
| 45 | 1:5 | 76% | 3 | 1.32 | 31.90 | 0.84 | 2.08 |
| 46 | 1:3 | 76% | 3 | 1.25 | 25.00 | 0.81 | 1.94 |
| 47 | 1:1 | 76% | 3 | 1.38 | 37.70 | 0.89 | 2.14 |
| **48** | **5:1** | **76%** | **3** | **1.65** | **64.50** | **1.05** | **2.57** |
| **49** | **8:1** | **76%** | **3** | **1.90** | **89.90** | **1.22** | **2.96** |
| 50 | 3:1 | 45% | 3 | 0.76 | -23.90 | 0.54 | 1.07 |
| 51 | 1:8 | 45% | 3 | 0.69 | -31.10 | 0.44 | 1.08 |
| 52 | 1:5 | 45% | 3 | 0.76 | -23.60 | 0.48 | 1.21 |
| 53 | 1:3 | 45% | 3 | 0.73 | -27.50 | 0.46 | 1.14 |
| 54 | 1:1 | 45% | 3 | 0.80 | -20.20 | 0.51 | 1.25 |
| 55 | 5:1 | 45% | 3 | 0.95 | -4.70 | 0.61 | 1.49 |
| 56 | 8:1 | 45% | 3 | 1.10 | 10.10 | 0.70 | 1.72 |
| 57 | 3:1 | 14% | 3 | 1.00 | 0.00 | 1.00 | 1 |
| 58 | 1:8 | 14% | 3 | 0.91 | -9.50 | 0.67 | 1.22 |
| 59 | 1:5 | 14% | 3 | 1.00 | 0.40 | 0.74 | 1.36 |

| | | | | | | | |
|---|---|---|---|---|---|---|---|
| 60 | 1:3 | 14% | 3 | 0.95 | -4.70 | 0.70 | 1.29 |
| 61 | 1:1 | 14% | 3 | 1.05 | 4.90 | 0.78 | 1.41 |
| 62 | 5:1 | 14% | 3 | 1.25 | 25.30 | 0.94 | 1.68 |
| 63 | 8:1 | 14% | 3 | 1.45 | 44.60 | 1.08 | 1.93 |

[a] Risk of death associated with every 1 SD increase in P:C, calculated as (1 – HR)*100; negative values = reduced risk of death, positive values = increased risk of death.

**Table S10.** Model summary from the Gamma generalized linear mixed-effects regression with a log link function and ANOVA (Type III Wald chisquare tests) to test cricket dietary preference (protein or carbohydrate).

| | Estimate (± SE) | $X^2$ | df | t value | Pr(>\|z\|) |
|---|---|---|---|---|---|
| **(Intercept)** | **-5.49 ± 0.11** | **2437.54** | **1** | **-49.37** | **<0.0001** |
| Week 1 Body Size (PC1) | 0.067 ± 0.042 | 2.60 | 1 | 1.61 | 0.11 |
| **Week** | **0.33 ± 0.014** | **573.04** | **1** | **23.94** | **<0.0001** |
| Diet Consumed | -0.037 ± 0.067 | 0.31 | 1 | -0.55 | 0.58 |
| Sex | -0.15 ± 0.14 | 1.16 | 1 | -1.08 | 0.28 |
| **Diet Consumed*Sex** | **0.32 ± 0.11** | **9.85** | **1** | **3.14** | **0.0017** |

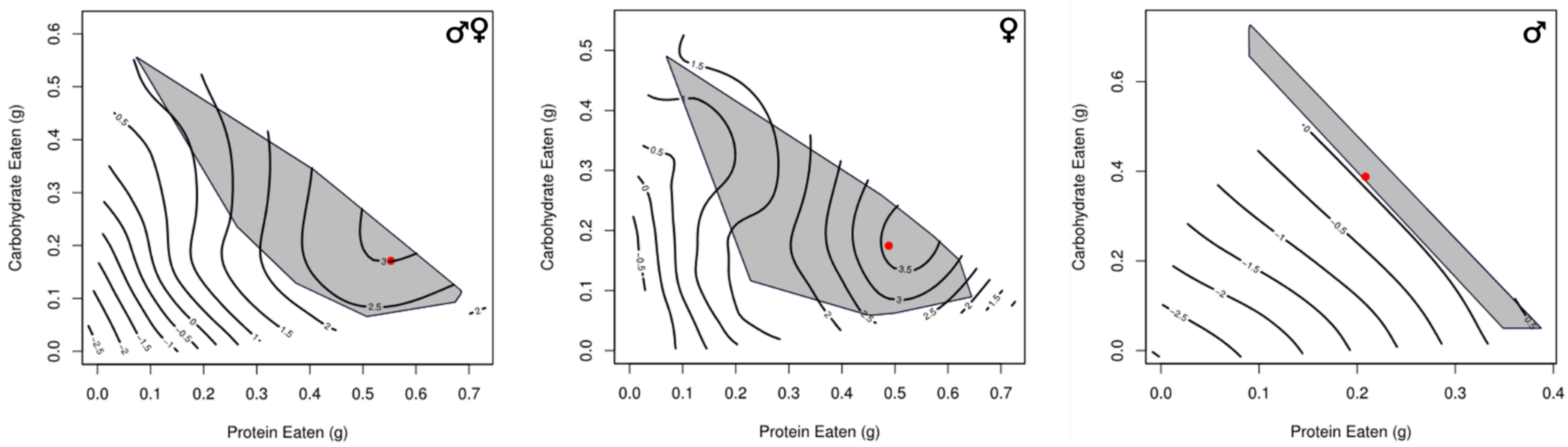


**Figure S1.** Ninety-five percent confidence regions (solid gray fill) for the global maximum (closed red circle) on each landscape for body size (principal component one) in *Gryllodes sigillatus*.

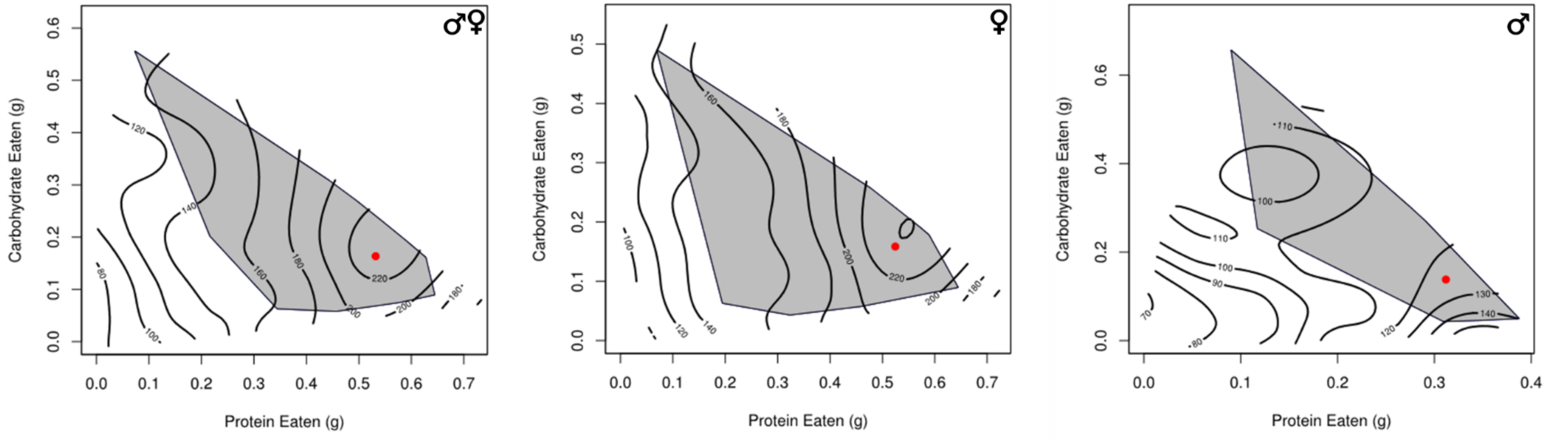


**Figure S2**. Ninety-five percent confidence regions (solid gray fill) for the global maximum (closed red circle) on each landscape for mass in *Gryllodes sigillatus*.

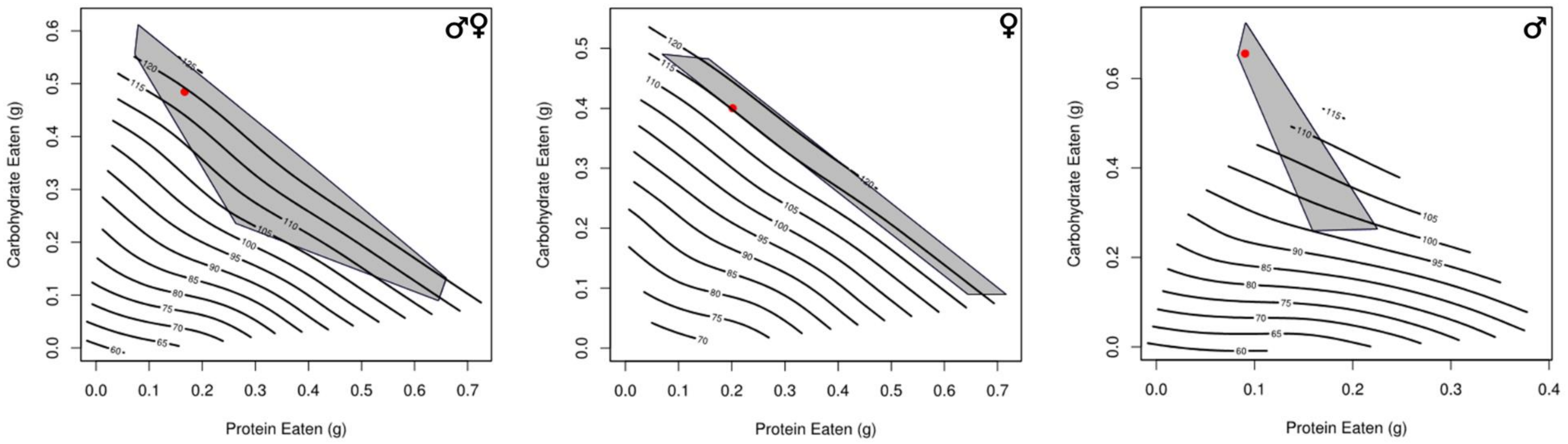


**Figure S3.** Ninety-five percent confidence regions (solid gray fill) for the global maximum (closed red circle) on each landscape for development time in *Gryllodes sigillatus*.
