## Supplementary material for "Applying nutritional ecology to optimize diets of crickets raised for food and feed": R Code

########GLMs associated with landscapes##########

library(readxl)

library(xlsx)

library(car)

library(MASS)

require(car)

require(MASS)

library(writexl)

library(ggplot2)

library(ggeffects)

library(splines)

library(sjPlot)

#DATA <- read_excel("C:/Users/Sarah Harrison/Desktop/Matt/Response Surfaces/MattsData.All - No Choice Exp-Jan 2024.xlsx")

DATA <- read_excel(".../Muzzatti.et.al.2024.xlsx", sheet = "GLMs")

\# Run this before running any pooled model

DATA$Diet=as.factor(DATA$Diet)

DATA$Sex=as.factor(DATA$Sex)

DATA$PC=as.factor(DATA$PC)

P<-DATA$Total.Pro.Eaten.g

C<-DATA$Total.Carb.Eaten.g

DATA<-as.data.frame(DATA)

str(DATA)

#Size Pooled

SizePC1.model.Pooled <- lm(SizePC1~P+C+I(P^2)+I(C^2)+P*C, data=DATA)

summary(SizePC1.model.Pooled)

Anova(SizePC1.model.Pooled)

#Dev Time Pooled

Dev.Time.model.Pooled <- lm(Adult.age.days~P+C+I(P^2)+I(C^2)+P*C, data=DATA)

summary(Dev.Time.model.Pooled)

Anova(Dev.Time.model.Pooled)

#Mass Pooled

Adult.Mass.model.Pooled <- lm(Adult.Mass.mg~P+C+I(P^2)+I(C^2)+P*C, data=DATA)

summary(Adult.Mass.model.Pooled)

Anova(Adult.Mass.model.Pooled)

\# Run pooled models again but remove the 2 outliers for total eaten from the data frame.

\# Compare to decide if you want to remove these outliers.

DATA.No.Outliers <- DATA [DATA$Total.Eaten.g <1.53,]

P<-DATA.No.Outliers$Total.Pro.Eaten.g

C<-DATA.No.Outliers$Total.Carb.Eaten.g

DATA.No.Outliers<-as.data.frame(DATA.No.Outliers)

str(DATA.No.Outliers)

#Size Pooled: With 2 outliers removed

SizePC1.No.Outliers<- lm(SizePC1~P+C+I(P^2)+I(C^2)+P*C, data=DATA.No.Outliers)

summary(SizePC1.No.Outliers)

Anova(SizePC1.No.Outliers)

#Dev Time Pooled : With 2 outliers removed

Dev.Time.No.Outliers <- lm(Adult.age.days~P+C+I(P^2)+I(C^2)+P*C, data=DATA.No.Outliers)

summary(Dev.Time.No.Outliers)

Anova(Dev.Time.No.Outliers)

#Mass Pooled: n=154 individuals

Adult.Mass.No.Outliers <- lm(Adult.Mass.mg~P+C+I(P^2)+I(C^2)+P*C, data=DATA.No.Outliers)

summary(Adult.Mass.No.Outliers)

Anova(Adult.Mass.No.Outliers)

#Run this right before running the female models

fDATA <- DATA[ which(DATA$Sex=='F'), ]

P<-fDATA$Total.Pro.Eaten.g

C<-fDATA$Total.Carb.Eaten.g

fDATA<-as.data.frame(fDATA)

str(fDATA)

#Size Females

SizePC1.model.Females <- lm(SizePC1~P+C+I(P^2)+I(C^2)+P*C, data=fDATA)

summary(SizePC1.model.Females)

Anova(SizePC1.model.Females)

#Dev Time Females

Dev.Time.model.Females <- lm(Adult.age.days~P+C+I(P^2)+I(C^2)+P*C, data=fDATA)

summary(Dev.Time.model.Females)

Anova(Dev.Time.model.Females)

#Mass - Females

Adult.Mass.model.Females <- lm(Adult.Mass.mg~P+C+I(P^2)+I(C^2)+P*C, data=fDATA)

summary(Adult.Mass.model.Females)

Anova(Adult.Mass.model.Females)

#Run this right before running the male models

mDATA <- DATA[ which(DATA$Sex=='M'), ]

P<-mDATA$Total.Pro.Eaten.g

C<-mDATA$Total.Carb.Eaten.g

mDATA<-as.data.frame(mDATA)

str(mDATA)

#Size Males

SizePC1.model.Males <- lm(SizePC1~P+C+I(P^2)+I(C^2)+P*C, data=mDATA)

summary(SizePC1.model.Males)

Anova(SizePC1.model.Males)

#Dev Time Males

Dev.Time.model.Males <- lm(Adult.age.days~P+C+I(P^2)+I(C^2)+P*C, data=mDATA)

summary(Dev.Time.model.Males)

Anova(Dev.Time.model.Males)

#Mass - Males

Adult.Mass.model.Males <- lm(Adult.Mass.mg~P+C+I(P^2)+I(C^2)+P*C, data=mDATA)

summary(Adult.Mass.model.Males)

Anova(Adult.Mass.model.Males)

###########################Landscapes and 95%CRs########################

install.packages(c("fields"))

library("fields")

#tips to avoid problems:

#If you are making new data spreadsheets, copy-paste your data (Do not copy any empty cells) into a new .cvs file.

#Dont copy extra data and then delete in the .cvs file - causes an error

\# Unless changing the code on line 20, the .cvs spreadsheets should have headers written as P, C, Response. In that order

#Close R and restart fresh in between multiple plots. When you get any error, always try restarting R.

surf.ind = read.csv(file.choose()) # this brings up a dialogue box (sometimes hidden on your taskbar) - use it to open the .cvs file

#these are saved as tabs in the Muzzatti.et.a.2024.Data sheet

#Open .cvs file from dialogue before running the following

surf.ind

attach(surf.ind)

names(surf.ind)

#edit(surf.ind)

library(fields)

library(ggplot2)

#Create thin plate spline eg Tps(cbind(x-axis, y-axis), Fitness)

surf.te=Tps(cbind(P, C),Response)#, lambda = 0.003) #Omitting lambda allows automatic estimation by minimizing GCV score

par(mar = c(7, 7, 5, 4))

summary(surf.te)

#surface(surf.te)#this is just the basic plot, with simple axis labels. Will give an error unless you expand the plots panel first

#Use this code for all plots/all .cvs spreadsheets. Just change the legend label (text = " ") for whatever plot you are currently doing

#Note: This code takes about 15 seconds to run

surface(surf.te,grid.list=list(P=seq(min(surf.ind$P),max(surf.ind$P),len=1000), # lower len gives pixelated contour lines and edges of the surface

C=seq(min(surf.ind$C),max(surf.ind$C),len=1000)), # but higher len values make the plot take much longer to load

xlab="Protein Eaten (g)",ylab="Carbohydrate Eaten (g)",

legend.args=list( text="Females - Weight", col="black",cex=1, side=1, line=-20)) #Change legend label to match which .csv you use

abline(0,1, lty=2, col="black", lwd=1) #1P:1C

abline(0,8, lty=2, lwd=1)	#8P:1C line

abline(0,5, lty=2, lwd=1)	#5P:1C line

abline(0,3, lty=2, lwd=1)	#3P:1C line

abline(0,0.333, lty=2, lwd=1)	#1P:3C line

abline(0,0.2, lty=2, lwd=1)	#1P:5C line

abline(0,0.125, lty=2, lwd=1)	#1P:8C line

points(0.523,0.159, pch=16) #manually add global optimum from CR (mean Point from OUT under global environment)

#global optima points

#All - weight points(0.532,0.165, pch=16)

#female - weight points(0.523,0.159, pch=16)

#male - weight points(0.31,0.141, pch=16)

#All - PC1 points(0.546,0.174, pch=16)

#female - PC1 points(0.489,0.174, pch=16)

#male - PC1 points(0.209,0.387, pch=16)

#Can we alter the x and y axis white space?

\# Dev Time - All - Max landscape needs the Carb axis extended in order to include global optima

\# Old y axis limit is max(surf.ind$C) = 0.5397881

\# meanPoint from 95 CR needs y axis to be > 0.5443

\# meanpoint from 95 CR for dev males needs y axis to be > 0.6477

surface(surf.te,grid.list=list(P=seq(0,max(surf.ind$P),len=1000), # lower len gives pixelated contour lines and edges of the surface

C=seq(0,0.7,len=1000)), # but higher len values make the plot take much longer to load

xlab="Protein Eaten (g)",ylab="Carbohydrate Eaten (g)",

legend.args=list( text="Males - Dev Time", col="black",cex=1, side=1, line=-24)) #Change legend label to match which .csv you use

abline(0,1, lty=2, col="black", lwd=1) #1P:1C

abline(0,8, lty=2, lwd=1)	#8P:1C line

abline(0,5, lty=2, lwd=1)	#5P:1C line

abline(0,3, lty=2, lwd=1)	#3P:1C line

abline(0,0.333, lty=2, lwd=1)	#1P:3C line

abline(0,0.2, lty=2, lwd=1)	#1P:5C line

abline(0,0.125, lty=2, lwd=1)	#1P:8C line

points(0.0927,0.6477, pch=16) #manually add global optimum from CR (mean Point from OUT under global environment)

#global optima for dev time

\# all max - points(0.0809,0.5443, pch=16)

\# all min - points(4.11e-18,2.97e-18, pch=16)

\# fem max - points(0.202,0.4, pch=16)

\# fem min - points(3.82e-18,3.20e-18, pch=16)

\# male max - points(0.0927,0.6477, pch=16)

\# male min - points(0.0126,0.0134, pch=16)

\# In legend.args, set line= -20 if you want to see the legend title in the preview

\# or set line= -24, export as 710 x 669 if you want the legend title to be in the right spot for high dpi image

\# You still wont be able to see the legend title in the export window, but it will be in the exported image

\## CONFIDENCE REGION GRAPHS

install.packages(c("OptimaRegion"))

library(OptimaRegion)

set.seed(1) # Run this before running 95 CR plots. Set of random numbers used for simulations will not change will not change

set.seed(Sys.time())#Use to reset R's random number generator behaviour to default

\# Load the appropriate .csv file for the Confidence Region you are running

\# You can skip this step if you have just run the code for the response surface for the same variable

\# The surf.ind dataframe then gets renamed for each confidence region plot below

surf.ind = read.csv(file.choose()) # this brings up a dialogue box (sometimes hidden on your taskbar) - use it to open the .cvs file

#these are saved as tabs in the Muzzatti.et.al.2024.Data excel file

###Body Size PC1 - All

CR.DATA.Size.ALL<-as.data.frame(surf.ind)

View(CR.DATA.Size.ALL)# Use to find the vertices that are within the landscape bounds

#Vertex1 from landscape extrapolation

#0.073 P/x

#0.618 C/y max

#Vertex2 from data

#0.71627942 P/x max

#0.08958410 C/y

set.seed(1)

out <- OptRegionTps(X=CR.DATA.Size.ALL[,1:2], y=CR.DATA.Size.ALL[,3], nosim = 500,

lambda = 0.01572, #take this value from summary(surf.te) from nutritional landscape code above

LB=c(0,0), # lower bounds (origin)

UB=c(0.71627942, 0.618), #upper bounds on the nutritional landscape (max on P/x axis , max on C/y axis)

xlab = "Protein Eaten (g)",

ylab = "Carbohydrate Eaten (g)",

triangularRegion=TRUE,

vertex1=c(0.073, 0.618),# vertex 1 (x,y) is the corner of the triangle near the far end of the carb (y) axis

vertex2=c(0.71627942, 0.08958410), # vertex 2 (x,y) is the corner of the triangle near the far end of the protein (x) axis

outputPDFFile = "95CR - Body Size PC1 - All.pdf")

\### Dev Time - All

CR.DATA.DevTime.ALL<-as.data.frame(surf.ind)

View(CR.DATA.DevTime.ALL)# Use to find the vertices that are within the landscape bounds

#Vertex1 from landscape extrapolation

#0.073 P/x

#0.618 C/y max

#Vertex2 from data

\# 0.71627942 P/x max

\# 0.08958410 C/y

\# CR on the Minimum

set.seed(1)

out <- OptRegionTps(X=CR.DATA.DevTime.ALL[,1:2], y=CR.DATA.DevTime.ALL[,3], nosim = 500,

lambda = 0.05640, #take this value from summary(surf.te) from nutritional landscape code above

LB=c(0,0), # lower bounds (origin)

UB=c(0.71627942, 0.618), #upper bounds on the nutritional landscape (max on P/x axis , max on C/y axis)

xlab = "Protein Eaten (g)",

ylab = "Carbohydrate Eaten (g)",

triangularRegion=TRUE,

vertex1=c(0.073, 0.618),# vertex 1 (x,y) is the corner of the triangle near the far end of the carb (y) axis

vertex2=c(0.71627942, 0.08958410), # vertex 2 (x,y) is the corner of the triangle near the far end of the protein (x) axis

maximization=FALSE, #looks for CR on minimum (trough) within the landscape

outputPDFFile = "95CR - Dev Time Min - All.pdf")

\# CR on the Maximum

set.seed(1)

out <- OptRegionTps(X=CR.DATA.DevTime.ALL[,1:2], y=CR.DATA.DevTime.ALL[,3], nosim = 500,

lambda = 0.05640, #take this value from summary(surf.te) from nutritional landscape code above

LB=c(0,0), # lower bounds (origin)

UB=c(0.71627942, 0.618), #upper bounds on the nutritional landscape (max on P/x axis , max on C/y axis)

xlab = "Protein Eaten (g)",

ylab = "Carbohydrate Eaten (g)",

triangularRegion=TRUE,

vertex1=c(0.073, 0.618),# vertex 1 (x,y) is the corner of the triangle near the far end of the carb (y) axis

vertex2=c(0.71627942, 0.08958410), # vertex 2 (x,y) is the corner of the triangle near the far end of the protein (x) axis

maximization=TRUE, #True is Default. Looks for CR on maximum (peak) within the landscape

outputPDFFile = "95CR - Dev Time Max - All.pdf")

###Weight - All

CR.DATA.Weight.ALL<-as.data.frame(surf.ind)

View(CR.DATA.Weight.ALL)# Use to find the vertices that are within the landscape bounds

#Vertex1 from landscape extrapolation

#0.073 P/x

#0.618 C/y max

#Vertex2 from data

#0.71627942 P/x max

#0.08958410 C/y

set.seed(1)

out <- OptRegionTps(X=CR.DATA.Weight.ALL[,1:2], y=CR.DATA.Weight.ALL[,3], nosim = 500,

lambda = 0.002767, #take this value from summary(surf.te) from nutritional landscape code above

LB=c(0,0), # lower bounds (origin)

UB=c(0.71627942, 0.618), #upper bounds on the nutritional landscape (max on P/x axis , max on C/y axis)

xlab = "Protein Eaten (g)",

ylab = "Carbohydrate Eaten (g)",

triangularRegion=TRUE,

vertex1=c(0.073, 0.618),# vertex 1 (x,y)is the corner of the triangle near the far end of the carb (y) axis

vertex2=c(0.71627942, 0.08958410), # vertex 2 (x,y) is the corner of the triangle near the far end of the protein (x) axis

outputPDFFile = "95CR - Weight - ALL.pdf")

###Body Size PC1 - Males

CR.DATA.Size.Males<-as.data.frame(surf.ind)

View(CR.DATA.Size.Males)# Use to find the vertices that are within the landscape bounds

#Vertex1 from landscape extrapolation

\# 0.09 P/x

\# 0.73 C/y max

#Vertex2 from landscape extrapolation

\# 0.3875 P/x max

\# 0.05 C/y

set.seed(1)

out <- OptRegionTps(X=CR.DATA.Size.Males[,1:2], y=CR.DATA.Size.Males[,3], nosim = 500,

lambda = 0.06730, #take this value from summary(surf.te) from nutritional landscape code above

LB=c(0,0), # lower bounds (origin)

UB=c(0.3875, 0.73), #upper bounds on the nutritional landscape (max on P/x axis , max on C/y axis)

xlab = "Protein Eaten (g)",

ylab = "Carbohydrate Eaten (g)",

triangularRegion=TRUE,

vertex1=c(0.09, 0.73),# vertex 1 (x,y) is the corner of the triangle near the far end of the carb (y) axis

vertex2=c(0.3875, 0.05), # vertex 2 (x,y) is the corner of the triangle near the far end of the protein (x) axis

outputPDFFile = "95CR - Body Size PC1 - Males.pdf")

###Dev Time - Males

CR.DATA.DevTime.Males<-as.data.frame(surf.ind)

View(CR.DATA.DevTime.Males)# Use to find the vertices that are within the landscape bounds

#Vertex1 from landscape extrapolation

\# 0.09 P/x

\# 0.73 C/y max

#Vertex2 from landscape extrapolation

\# 0.3875 P/x max

\# 0.05 C/y

#CR on the maximum (peak)

set.seed(1)

out <- OptRegionTps(X=CR.DATA.DevTime.Males[,1:2], y=CR.DATA.DevTime.Males[,3], nosim = 500,

lambda = 0.04455, #take this value from summary(surf.te) from nutritional landscape code above

LB=c(0,0), # lower bounds (origin)

UB=c(0.3875, 0.73), #upper bounds on the nutritional landscape (max on P/x axis , max on C/y axis)

xlab = "Protein Eaten (g)",

ylab = "Carbohydrate Eaten (g)",

triangularRegion=TRUE,

vertex1=c(0.09, 0.73),# vertex 1 (x,y) is the corner of the triangle near the far end of the carb (y) axis

vertex2=c(0.3875, 0.05), # vertex 2 (x,y) is the corner of the triangle near the far end of the protein (x) axis

maximization=TRUE, #True is Default. Looks for CR on maximum (peak) within the landscape

outputPDFFile = "95CR - Dev Time Max - Males.pdf")

###Weight - Males

CR.DATA.Weight.Males<-as.data.frame(surf.ind)

View(CR.DATA.Weight.Males)# Use to find the vertices that are within the landscape bounds

#Vertex1 from landscape extrapolation

\# 0.09 P/x

\# 0.73 C/y max

#Vertex2 from landscape extrapolation

\# 0.3875 P/x max

\# 0.05 C/y

set.seed(1)

out <- OptRegionTps(X=CR.DATA.Weight.Males[,1:2], y=CR.DATA.Weight.Males[,3], nosim = 500,

lambda = 0.003104, #take this value from summary(surf.te) from nutritional landscape code above

LB=c(0,0), # lower bounds (origin)

UB=c(0.3875, 0.73), #upper bounds on the nutritional landscape (max on P/x axis , max on C/y axis)

xlab = "Protein Eaten (g)",

ylab = "Carbohydrate Eaten (g)",

triangularRegion=TRUE,

vertex1=c(0.09, 0.73),# vertex 1 (x,y) is the corner of the triangle near the far end of the carb (y) axis

vertex2=c(0.3875, 0.05), # vertex 2 (x,y) is the corner of the triangle near the far end of the protein (x) axis

outputPDFFile = "95CR - Weight - Males.pdf")

###Body Size PC1 - Females

CR.DATA.Size.Females<-as.data.frame(surf.ind)

View(CR.DATA.Size.Females)# Use to find the vertices that are within the landscape bounds

#Vertex1 from landscape extrapolation

#0.069 P/x

#0.545 C/y max

#Vertex2 from data

\# 0.71627942 P/x max

\# 0.08958410 C/y

set.seed(1)

out <- OptRegionTps(X=CR.DATA.Size.Females[,1:2], y=CR.DATA.Size.Females[,3], nosim = 500,

lambda = 0.002367, #take this value from summary(surf.te) from nutritional landscape code above

LB=c(0,0), # lower bounds (origin)

UB=c(0.71627942,0.545), #upper bounds on the nutritional landscape (max on P/x axis , max on C/y axis)

xlab = "Protein Eaten (g)",

ylab = "Carbohydrate Eaten (g)",

triangularRegion=TRUE,

vertex1=c(0.069, 0.545),# vertex 1 (x,y)is the corner of the triangle near the far end of the carb (y) axis

vertex2=c(0.71627942, 0.08958410), # vertex 2 (x,y) is the corner of the triangle near the far end of the protein (x) axis

outputPDFFile = "95CR - Body Size PC1 - Females.pdf")

###Dev Time - Females

CR.DATA.DevTime.Females<-as.data.frame(surf.ind)

View(CR.DATA.DevTime.Females)# Use to find the vertices that are within the landscape bounds

#Vertex1 from landscape extrapolation

#0.069 P/x

#0.545 C/y max

#Vertex2 from data

\# 0.71627942 P/x max

\# 0.08958410 C/y

#CR on the minimum (trough)

set.seed(1)

out <- OptRegionTps(X=CR.DATA.DevTime.Females[,1:2], y=CR.DATA.DevTime.Females[,3], nosim = 500,

lambda = 0.06085, #take this value from summary(surf.te) from nutritional landscape code above

LB=c(0,0), # lower bounds (origin)

UB=c(0.71627942,0.545), #upper bounds on the nutritional landscape (max on P/x axis , max on C/y axis)

xlab = "Protein Eaten (g)",

ylab = "Carbohydrate Eaten (g)",

triangularRegion=TRUE,

vertex1=c(0.069, 0.545),# vertex 1 (x,y)is the corner of the triangle near the far end of the carb (y) axis

vertex2=c(0.71627942, 0.08958410), # vertex 2 (x,y) is the corner of the triangle near the far end of the protein (x) axis

maximization=FALSE,#looks for the CR on the minimum (trough) within the landscape

outputPDFFile = "95CR - Dev Time Min - Females.pdf")

#CR on the maximum (peak)

set.seed(1)

out <- OptRegionTps(X=CR.DATA.DevTime.Females[,1:2], y=CR.DATA.DevTime.Females[,3], nosim = 500,

lambda = 0.06085, #take this value from summary(surf.te) from nutritional landscape code above

LB=c(0,0), # lower bounds (origin)

UB=c(0.71627942,0.545), #upper bounds on the nutritional landscape (max on P/x axis , max on C/y axis)

xlab = "Protein Eaten (g)",

ylab = "Carbohydrate Eaten (g)",

triangularRegion=TRUE,

vertex1=c(0.069, 0.545),# vertex 1 (x,y)is the corner of the triangle near the far end of the carb (y) axis

vertex2=c(0.71627942, 0.08958410), # vertex 2 (x,y) is the corner of the triangle near the far end of the protein (x) axis

maximization=TRUE, #True is Default. Looks for CR on maximum (peak) within the landscape

outputPDFFile = "95CR - Dev Time Max - Females.pdf")

###Weight - Females

CR.DATA.Weight.Females<-as.data.frame(surf.ind)

View(CR.DATA.Weight.Females)# Use to find the vertices that are within the landscape bounds

#Vertex1 from landscape extrapolation

#0.069 P/x

#0.545 C/y max

#Vertex2 from data

\# 0.71627942 P/x max

\# 0.08958410 C/y

set.seed(1)

out <- OptRegionTps(X=CR.DATA.Weight.Females[,1:2], y=CR.DATA.Weight.Females[,3], nosim = 500,

lambda = 0.001976, #take this value from summary(surf.te) from nutritional landscape code above

LB=c(0,0), # lower bounds (origin)

UB=c(0.71627942,0.545), #upper bounds on the nutritional landscape (max on P/x axis , max on C/y axis)

xlab = "Protein Eaten (g)",

ylab = "Carbohydrate Eaten (g)",

triangularRegion=TRUE,

vertex1=c(0.069, 0.545),# vertex 1 (x,y)is the corner of the triangle near the far end of the carb (y) axis

vertex2=c(0.71627942, 0.08958410), # vertex 2 (x,y) is the corner of the triangle near the far end of the protein (x) axis

outputPDFFile = "95CR - Weight - Females.pdf")

\## Dev Time Plots without CRs:

#Dev Time - All - Minimum: 20 simulations/all with different random numbers

surf.ind = read.csv(file.choose())#Select Dev Time All data

CR.DATA.DevTime.ALL<-as.data.frame(surf.ind)

#Vertex1 from landscape extrapolation

#0.073 P/x

#0.618 C/y max

#Vertex2 from data

\# 0.71627942 P/x max

\# 0.08958410 C/y

set.seed(1, kind="Wichmann-Hill")

out <- OptRegionTps(X=CR.DATA.DevTime.ALL[,1:2], y=CR.DATA.DevTime.ALL[,3], nosim = 2000,

lambda = 0.05640, #take this value from summary(surf.te) from nutritional landscape code above

LB=c(0,0), # lower bounds (origin)

UB=c(0.71627942, 0.618), #upper bounds on the nutritional landscape (max on P/x axis , max on C/y axis)

xlab = "Protein Eaten (g)",

ylab = "Carbohydrate Eaten (g)",

triangularRegion=TRUE,

vertex1=c(0.073, 0.618),# vertex 1 (x,y) is the corner of the triangle near the far end of the carb (y) axis

vertex2=c(0.71627942, 0.08958410), # vertex 2 (x,y) is the corner of the triangle near the far end of the protein (x) axis

maximization=FALSE, #looks for CR on minimum (trough) within the landscape

outputPDFFile = "95CR - Dev Time Min - All -WH1.pdf")

set.seed(2, kind="Wichmann-Hill")

out <- OptRegionTps(X=CR.DATA.DevTime.ALL[,1:2], y=CR.DATA.DevTime.ALL[,3], nosim = 2000,

lambda = 0.05640, #take this value from summary(surf.te) from nutritional landscape code above

LB=c(0,0), # lower bounds (origin)

UB=c(0.71627942, 0.618), #upper bounds on the nutritional landscape (max on P/x axis , max on C/y axis)

xlab = "Protein Eaten (g)",

ylab = "Carbohydrate Eaten (g)",

triangularRegion=TRUE,

vertex1=c(0.073, 0.618),# vertex 1 (x,y) is the corner of the triangle near the far end of the carb (y) axis

vertex2=c(0.71627942, 0.08958410), # vertex 2 (x,y) is the corner of the triangle near the far end of the protein (x) axis

maximization=FALSE, #looks for CR on minimum (trough) within the landscape

outputPDFFile = "95CR - Dev Time Min - All -WH2.pdf")

set.seed(3, kind="Wichmann-Hill")

out <- OptRegionTps(X=CR.DATA.DevTime.ALL[,1:2], y=CR.DATA.DevTime.ALL[,3], nosim = 2000,

lambda = 0.05640, #take this value from summary(surf.te) from nutritional landscape code above

LB=c(0,0), # lower bounds (origin)

UB=c(0.71627942, 0.618), #upper bounds on the nutritional landscape (max on P/x axis , max on C/y axis)

xlab = "Protein Eaten (g)",

ylab = "Carbohydrate Eaten (g)",

triangularRegion=TRUE,

vertex1=c(0.073, 0.618),# vertex 1 (x,y) is the corner of the triangle near the far end of the carb (y) axis

vertex2=c(0.71627942, 0.08958410), # vertex 2 (x,y) is the corner of the triangle near the far end of the protein (x) axis

maximization=FALSE, #looks for CR on minimum (trough) within the landscape

outputPDFFile = "95CR - Dev Time Min - All -WH3.pdf")

set.seed(4, kind="Wichmann-Hill")

out <- OptRegionTps(X=CR.DATA.DevTime.ALL[,1:2], y=CR.DATA.DevTime.ALL[,3], nosim = 2000,

lambda = 0.05640, #take this value from summary(surf.te) from nutritional landscape code above

LB=c(0,0), # lower bounds (origin)

UB=c(0.71627942, 0.618), #upper bounds on the nutritional landscape (max on P/x axis , max on C/y axis)

xlab = "Protein Eaten (g)",

ylab = "Carbohydrate Eaten (g)",

triangularRegion=TRUE,

vertex1=c(0.073, 0.618),# vertex 1 (x,y) is the corner of the triangle near the far end of the carb (y) axis

vertex2=c(0.71627942, 0.08958410), # vertex 2 (x,y) is the corner of the triangle near the far end of the protein (x) axis

maximization=FALSE, #looks for CR on minimum (trough) within the landscape

outputPDFFile = "95CR - Dev Time Min - All -WH4.pdf")

set.seed(5, kind="Wichmann-Hill")

out <- OptRegionTps(X=CR.DATA.DevTime.ALL[,1:2], y=CR.DATA.DevTime.ALL[,3], nosim = 2000,

lambda = 0.05640, #take this value from summary(surf.te) from nutritional landscape code above

LB=c(0,0), # lower bounds (origin)

UB=c(0.71627942, 0.618), #upper bounds on the nutritional landscape (max on P/x axis , max on C/y axis)

xlab = "Protein Eaten (g)",

ylab = "Carbohydrate Eaten (g)",

triangularRegion=TRUE,

vertex1=c(0.073, 0.618),# vertex 1 (x,y) is the corner of the triangle near the far end of the carb (y) axis

vertex2=c(0.71627942, 0.08958410), # vertex 2 (x,y) is the corner of the triangle near the far end of the protein (x) axis

maximization=FALSE, #looks for CR on minimum (trough) within the landscape

outputPDFFile = "95CR - Dev Time Min - All -WH5.pdf")

set.seed(6, kind="Wichmann-Hill")

out <- OptRegionTps(X=CR.DATA.DevTime.ALL[,1:2], y=CR.DATA.DevTime.ALL[,3], nosim = 2000,

lambda = 0.05640, #take this value from summary(surf.te) from nutritional landscape code above

LB=c(0,0), # lower bounds (origin)

UB=c(0.71627942, 0.618), #upper bounds on the nutritional landscape (max on P/x axis , max on C/y axis)

xlab = "Protein Eaten (g)",

ylab = "Carbohydrate Eaten (g)",

triangularRegion=TRUE,

vertex1=c(0.073, 0.618),# vertex 1 (x,y) is the corner of the triangle near the far end of the carb (y) axis

vertex2=c(0.71627942, 0.08958410), # vertex 2 (x,y) is the corner of the triangle near the far end of the protein (x) axis

maximization=FALSE, #looks for CR on minimum (trough) within the landscape

outputPDFFile = "95CR - Dev Time Min - All -WH6.pdf")

set.seed(7, kind="Wichmann-Hill")

out <- OptRegionTps(X=CR.DATA.DevTime.ALL[,1:2], y=CR.DATA.DevTime.ALL[,3], nosim = 2000,

lambda = 0.05640, #take this value from summary(surf.te) from nutritional landscape code above

LB=c(0,0), # lower bounds (origin)

UB=c(0.71627942, 0.618), #upper bounds on the nutritional landscape (max on P/x axis , max on C/y axis)

xlab = "Protein Eaten (g)",

ylab = "Carbohydrate Eaten (g)",

triangularRegion=TRUE,

vertex1=c(0.073, 0.618),# vertex 1 (x,y) is the corner of the triangle near the far end of the carb (y) axis

vertex2=c(0.71627942, 0.08958410), # vertex 2 (x,y) is the corner of the triangle near the far end of the protein (x) axis

maximization=FALSE, #looks for CR on minimum (trough) within the landscape

outputPDFFile = "95CR - Dev Time Min - All -WH7.pdf")

set.seed(8, kind="Wichmann-Hill")

out <- OptRegionTps(X=CR.DATA.DevTime.ALL[,1:2], y=CR.DATA.DevTime.ALL[,3], nosim = 2000,

lambda = 0.05640, #take this value from summary(surf.te) from nutritional landscape code above

LB=c(0,0), # lower bounds (origin)

UB=c(0.71627942, 0.618), #upper bounds on the nutritional landscape (max on P/x axis , max on C/y axis)

xlab = "Protein Eaten (g)",

ylab = "Carbohydrate Eaten (g)",

triangularRegion=TRUE,

vertex1=c(0.073, 0.618),# vertex 1 (x,y) is the corner of the triangle near the far end of the carb (y) axis

vertex2=c(0.71627942, 0.08958410), # vertex 2 (x,y) is the corner of the triangle near the far end of the protein (x) axis

maximization=FALSE, #looks for CR on minimum (trough) within the landscape

outputPDFFile = "95CR - Dev Time Min - All -WH8.pdf")

set.seed(9, kind="Wichmann-Hill")

out <- OptRegionTps(X=CR.DATA.DevTime.ALL[,1:2], y=CR.DATA.DevTime.ALL[,3], nosim = 2000,

lambda = 0.05640, #take this value from summary(surf.te) from nutritional landscape code above

LB=c(0,0), # lower bounds (origin)

UB=c(0.71627942, 0.618), #upper bounds on the nutritional landscape (max on P/x axis , max on C/y axis)

xlab = "Protein Eaten (g)",

ylab = "Carbohydrate Eaten (g)",

triangularRegion=TRUE,

vertex1=c(0.073, 0.618),# vertex 1 (x,y) is the corner of the triangle near the far end of the carb (y) axis

vertex2=c(0.71627942, 0.08958410), # vertex 2 (x,y) is the corner of the triangle near the far end of the protein (x) axis

maximization=FALSE, #looks for CR on minimum (trough) within the landscape

outputPDFFile = "95CR - Dev Time Min - All -WH9.pdf")

set.seed(10, kind="Wichmann-Hill")

out <- OptRegionTps(X=CR.DATA.DevTime.ALL[,1:2], y=CR.DATA.DevTime.ALL[,3], nosim = 2000,

lambda = 0.05640, #take this value from summary(surf.te) from nutritional landscape code above

LB=c(0,0), # lower bounds (origin)

UB=c(0.71627942, 0.618), #upper bounds on the nutritional landscape (max on P/x axis , max on C/y axis)

xlab = "Protein Eaten (g)",

ylab = "Carbohydrate Eaten (g)",

triangularRegion=TRUE,

vertex1=c(0.073, 0.618),# vertex 1 (x,y) is the corner of the triangle near the far end of the carb (y) axis

vertex2=c(0.71627942, 0.08958410), # vertex 2 (x,y) is the corner of the triangle near the far end of the protein (x) axis

maximization=FALSE, #looks for CR on minimum (trough) within the landscape

outputPDFFile = "95CR - Dev Time Min - All -WH10.pdf")

set.seed(11, kind="Wichmann-Hill")

out <- OptRegionTps(X=CR.DATA.DevTime.ALL[,1:2], y=CR.DATA.DevTime.ALL[,3], nosim = 2000,

lambda = 0.05640, #take this value from summary(surf.te) from nutritional landscape code above

LB=c(0,0), # lower bounds (origin)

UB=c(0.71627942, 0.618), #upper bounds on the nutritional landscape (max on P/x axis , max on C/y axis)

xlab = "Protein Eaten (g)",

ylab = "Carbohydrate Eaten (g)",

triangularRegion=TRUE,

vertex1=c(0.073, 0.618),# vertex 1 (x,y) is the corner of the triangle near the far end of the carb (y) axis

vertex2=c(0.71627942, 0.08958410), # vertex 2 (x,y) is the corner of the triangle near the far end of the protein (x) axis

maximization=FALSE, #looks for CR on minimum (trough) within the landscape

outputPDFFile = "95CR - Dev Time Min - All -WH11.pdf")

set.seed(12, kind="Wichmann-Hill")

out <- OptRegionTps(X=CR.DATA.DevTime.ALL[,1:2], y=CR.DATA.DevTime.ALL[,3], nosim = 2000,

lambda = 0.05640, #take this value from summary(surf.te) from nutritional landscape code above

LB=c(0,0), # lower bounds (origin)

UB=c(0.71627942, 0.618), #upper bounds on the nutritional landscape (max on P/x axis , max on C/y axis)

xlab = "Protein Eaten (g)",

ylab = "Carbohydrate Eaten (g)",

triangularRegion=TRUE,

vertex1=c(0.073, 0.618),# vertex 1 (x,y) is the corner of the triangle near the far end of the carb (y) axis

vertex2=c(0.71627942, 0.08958410), # vertex 2 (x,y) is the corner of the triangle near the far end of the protein (x) axis

maximization=FALSE, #looks for CR on minimum (trough) within the landscape

outputPDFFile = "95CR - Dev Time Min - All -WH12.pdf")

set.seed(13, kind="Wichmann-Hill")

out <- OptRegionTps(X=CR.DATA.DevTime.ALL[,1:2], y=CR.DATA.DevTime.ALL[,3], nosim = 2000,

lambda = 0.05640, #take this value from summary(surf.te) from nutritional landscape code above

LB=c(0,0), # lower bounds (origin)

UB=c(0.71627942, 0.618), #upper bounds on the nutritional landscape (max on P/x axis , max on C/y axis)

xlab = "Protein Eaten (g)",

ylab = "Carbohydrate Eaten (g)",

triangularRegion=TRUE,

vertex1=c(0.073, 0.618),# vertex 1 (x,y) is the corner of the triangle near the far end of the carb (y) axis

vertex2=c(0.71627942, 0.08958410), # vertex 2 (x,y) is the corner of the triangle near the far end of the protein (x) axis

maximization=FALSE, #looks for CR on minimum (trough) within the landscape

outputPDFFile = "95CR - Dev Time Min - All -WH13.pdf")

set.seed(14, kind="Wichmann-Hill")

out <- OptRegionTps(X=CR.DATA.DevTime.ALL[,1:2], y=CR.DATA.DevTime.ALL[,3], nosim = 2000,

lambda = 0.05640, #take this value from summary(surf.te) from nutritional landscape code above

LB=c(0,0), # lower bounds (origin)

UB=c(0.71627942, 0.618), #upper bounds on the nutritional landscape (max on P/x axis , max on C/y axis)

xlab = "Protein Eaten (g)",

ylab = "Carbohydrate Eaten (g)",

triangularRegion=TRUE,

vertex1=c(0.073, 0.618),# vertex 1 (x,y) is the corner of the triangle near the far end of the carb (y) axis

vertex2=c(0.71627942, 0.08958410), # vertex 2 (x,y) is the corner of the triangle near the far end of the protein (x) axis

maximization=FALSE, #looks for CR on minimum (trough) within the landscape

outputPDFFile = "95CR - Dev Time Min - All -WH14.pdf")

set.seed(15, kind="Wichmann-Hill")

out <- OptRegionTps(X=CR.DATA.DevTime.ALL[,1:2], y=CR.DATA.DevTime.ALL[,3], nosim = 2000,

lambda = 0.05640, #take this value from summary(surf.te) from nutritional landscape code above

LB=c(0,0), # lower bounds (origin)

UB=c(0.71627942, 0.618), #upper bounds on the nutritional landscape (max on P/x axis , max on C/y axis)

xlab = "Protein Eaten (g)",

ylab = "Carbohydrate Eaten (g)",

triangularRegion=TRUE,

vertex1=c(0.073, 0.618),# vertex 1 (x,y) is the corner of the triangle near the far end of the carb (y) axis

vertex2=c(0.71627942, 0.08958410), # vertex 2 (x,y) is the corner of the triangle near the far end of the protein (x) axis

maximization=FALSE, #looks for CR on minimum (trough) within the landscape

outputPDFFile = "95CR - Dev Time Min - All -WH15.pdf")

set.seed(16, kind="Wichmann-Hill")

out <- OptRegionTps(X=CR.DATA.DevTime.ALL[,1:2], y=CR.DATA.DevTime.ALL[,3], nosim = 2000,

lambda = 0.05640, #take this value from summary(surf.te) from nutritional landscape code above

LB=c(0,0), # lower bounds (origin)

UB=c(0.71627942, 0.618), #upper bounds on the nutritional landscape (max on P/x axis , max on C/y axis)

xlab = "Protein Eaten (g)",

ylab = "Carbohydrate Eaten (g)",

triangularRegion=TRUE,

vertex1=c(0.073, 0.618),# vertex 1 (x,y) is the corner of the triangle near the far end of the carb (y) axis

vertex2=c(0.71627942, 0.08958410), # vertex 2 (x,y) is the corner of the triangle near the far end of the protein (x) axis

maximization=FALSE, #looks for CR on minimum (trough) within the landscape

outputPDFFile = "95CR - Dev Time Min - All -WH16.pdf")

set.seed(17, kind="Wichmann-Hill")

out <- OptRegionTps(X=CR.DATA.DevTime.ALL[,1:2], y=CR.DATA.DevTime.ALL[,3], nosim = 2000,

lambda = 0.05640, #take this value from summary(surf.te) from nutritional landscape code above

LB=c(0,0), # lower bounds (origin)

UB=c(0.71627942, 0.618), #upper bounds on the nutritional landscape (max on P/x axis , max on C/y axis)

xlab = "Protein Eaten (g)",

ylab = "Carbohydrate Eaten (g)",

triangularRegion=TRUE,

vertex1=c(0.073, 0.618),# vertex 1 (x,y) is the corner of the triangle near the far end of the carb (y) axis

vertex2=c(0.71627942, 0.08958410), # vertex 2 (x,y) is the corner of the triangle near the far end of the protein (x) axis

maximization=FALSE, #looks for CR on minimum (trough) within the landscape

outputPDFFile = "95CR - Dev Time Min - All -WH17.pdf")

set.seed(18, kind="Wichmann-Hill")

out <- OptRegionTps(X=CR.DATA.DevTime.ALL[,1:2], y=CR.DATA.DevTime.ALL[,3], nosim = 2000,

lambda = 0.05640, #take this value from summary(surf.te) from nutritional landscape code above

LB=c(0,0), # lower bounds (origin)

UB=c(0.71627942, 0.618), #upper bounds on the nutritional landscape (max on P/x axis , max on C/y axis)

xlab = "Protein Eaten (g)",

ylab = "Carbohydrate Eaten (g)",

triangularRegion=TRUE,

vertex1=c(0.073, 0.618),# vertex 1 (x,y) is the corner of the triangle near the far end of the carb (y) axis

vertex2=c(0.71627942, 0.08958410), # vertex 2 (x,y) is the corner of the triangle near the far end of the protein (x) axis

maximization=FALSE, #looks for CR on minimum (trough) within the landscape

outputPDFFile = "95CR - Dev Time Min - All -WH18.pdf")

set.seed(19, kind="Wichmann-Hill")

out <- OptRegionTps(X=CR.DATA.DevTime.ALL[,1:2], y=CR.DATA.DevTime.ALL[,3], nosim = 2000,

lambda = 0.05640, #take this value from summary(surf.te) from nutritional landscape code above

LB=c(0,0), # lower bounds (origin)

UB=c(0.71627942, 0.618), #upper bounds on the nutritional landscape (max on P/x axis , max on C/y axis)

xlab = "Protein Eaten (g)",

ylab = "Carbohydrate Eaten (g)",

triangularRegion=TRUE,

vertex1=c(0.073, 0.618),# vertex 1 (x,y) is the corner of the triangle near the far end of the carb (y) axis

vertex2=c(0.71627942, 0.08958410), # vertex 2 (x,y) is the corner of the triangle near the far end of the protein (x) axis

maximization=FALSE, #looks for CR on minimum (trough) within the landscape

outputPDFFile = "95CR - Dev Time Min - All -WH19.pdf")

set.seed(20, kind="Wichmann-Hill")

out <- OptRegionTps(X=CR.DATA.DevTime.ALL[,1:2], y=CR.DATA.DevTime.ALL[,3], nosim = 2000,

lambda = 0.05640, #take this value from summary(surf.te) from nutritional landscape code above

LB=c(0,0), # lower bounds (origin)

UB=c(0.71627942, 0.618), #upper bounds on the nutritional landscape (max on P/x axis , max on C/y axis)

xlab = "Protein Eaten (g)",

ylab = "Carbohydrate Eaten (g)",

triangularRegion=TRUE,

vertex1=c(0.073, 0.618),# vertex 1 (x,y) is the corner of the triangle near the far end of the carb (y) axis

vertex2=c(0.71627942, 0.08958410), # vertex 2 (x,y) is the corner of the triangle near the far end of the protein (x) axis

maximization=FALSE, #looks for CR on minimum (trough) within the landscape

outputPDFFile = "95CR - Dev Time Min - All -WH20.pdf")

#Dev Time Females Min

surf.ind = read.csv(file.choose())#Select Dev Time Female data

CR.DATA.DevTime.Females<-as.data.frame(surf.ind)

View(CR.DATA.DevTime.Females)# Use to find the vertices that are within the landscape bounds

#Vertex1 from landscape extrapolation

#0.069 P/x

#0.545 C/y max

#Vertex2 from data

\# 0.71627942 P/x max

\# 0.08958410 C/y

#CR on the minimum (trough)

set.seed(1, kind="Wichmann-Hill")

out <- OptRegionTps(X=CR.DATA.DevTime.Females[,1:2], y=CR.DATA.DevTime.Females[,3], nosim = 2000,

lambda = 0.06085, #take this value from summary(surf.te) from nutritional landscape code above

LB=c(0,0), # lower bounds (origin)

UB=c(0.71627942,0.545), #upper bounds on the nutritional landscape (max on P/x axis , max on C/y axis)

xlab = "Protein Eaten (g)",

ylab = "Carbohydrate Eaten (g)",

triangularRegion=TRUE,

vertex1=c(0.069, 0.545),# vertex 1 (x,y)is the corner of the triangle near the far end of the carb (y) axis

vertex2=c(0.71627942, 0.08958410), # vertex 2 (x,y) is the corner of the triangle near the far end of the protein (x) axis

maximization=FALSE,#looks for the CR on the minimum (trough) within the landscape

outputPDFFile = "95CR - Dev Time Min - Females- WH1.pdf")

set.seed(2, kind="Wichmann-Hill")

out <- OptRegionTps(X=CR.DATA.DevTime.Females[,1:2], y=CR.DATA.DevTime.Females[,3], nosim = 2000,

lambda = 0.06085, #take this value from summary(surf.te) from nutritional landscape code above

LB=c(0,0), # lower bounds (origin)

UB=c(0.71627942,0.545), #upper bounds on the nutritional landscape (max on P/x axis , max on C/y axis)

xlab = "Protein Eaten (g)",

ylab = "Carbohydrate Eaten (g)",

triangularRegion=TRUE,

vertex1=c(0.069, 0.545),# vertex 1 (x,y)is the corner of the triangle near the far end of the carb (y) axis

vertex2=c(0.71627942, 0.08958410), # vertex 2 (x,y) is the corner of the triangle near the far end of the protein (x) axis

maximization=FALSE,#looks for the CR on the minimum (trough) within the landscape

outputPDFFile = "95CR - Dev Time Min - Females- WH2.pdf")

set.seed(3, kind="Wichmann-Hill")

out <- OptRegionTps(X=CR.DATA.DevTime.Females[,1:2], y=CR.DATA.DevTime.Females[,3], nosim = 2000,

lambda = 0.06085, #take this value from summary(surf.te) from nutritional landscape code above

LB=c(0,0), # lower bounds (origin)

UB=c(0.71627942,0.545), #upper bounds on the nutritional landscape (max on P/x axis , max on C/y axis)

xlab = "Protein Eaten (g)",

ylab = "Carbohydrate Eaten (g)",

triangularRegion=TRUE,

vertex1=c(0.069, 0.545),# vertex 1 (x,y)is the corner of the triangle near the far end of the carb (y) axis

vertex2=c(0.71627942, 0.08958410), # vertex 2 (x,y) is the corner of the triangle near the far end of the protein (x) axis

maximization=FALSE,#looks for the CR on the minimum (trough) within the landscape

outputPDFFile = "95CR - Dev Time Min - Females- WH3.pdf")

set.seed(4, kind="Wichmann-Hill")

out <- OptRegionTps(X=CR.DATA.DevTime.Females[,1:2], y=CR.DATA.DevTime.Females[,3], nosim = 2000,

lambda = 0.06085, #take this value from summary(surf.te) from nutritional landscape code above

LB=c(0,0), # lower bounds (origin)

UB=c(0.71627942,0.545), #upper bounds on the nutritional landscape (max on P/x axis , max on C/y axis)

xlab = "Protein Eaten (g)",

ylab = "Carbohydrate Eaten (g)",

triangularRegion=TRUE,

vertex1=c(0.069, 0.545),# vertex 1 (x,y)is the corner of the triangle near the far end of the carb (y) axis

vertex2=c(0.71627942, 0.08958410), # vertex 2 (x,y) is the corner of the triangle near the far end of the protein (x) axis

maximization=FALSE,#looks for the CR on the minimum (trough) within the landscape

outputPDFFile = "95CR - Dev Time Min - Females- WH4.pdf")

set.seed(5, kind="Wichmann-Hill")

out <- OptRegionTps(X=CR.DATA.DevTime.Females[,1:2], y=CR.DATA.DevTime.Females[,3], nosim = 2000,

lambda = 0.06085, #take this value from summary(surf.te) from nutritional landscape code above

LB=c(0,0), # lower bounds (origin)

UB=c(0.71627942,0.545), #upper bounds on the nutritional landscape (max on P/x axis , max on C/y axis)

xlab = "Protein Eaten (g)",

ylab = "Carbohydrate Eaten (g)",

triangularRegion=TRUE,

vertex1=c(0.069, 0.545),# vertex 1 (x,y)is the corner of the triangle near the far end of the carb (y) axis

vertex2=c(0.71627942, 0.08958410), # vertex 2 (x,y) is the corner of the triangle near the far end of the protein (x) axis

maximization=FALSE,#looks for the CR on the minimum (trough) within the landscape

outputPDFFile = "95CR - Dev Time Min - Females- WH5.pdf")

set.seed(6, kind="Wichmann-Hill")

out <- OptRegionTps(X=CR.DATA.DevTime.Females[,1:2], y=CR.DATA.DevTime.Females[,3], nosim = 2000,

lambda = 0.06085, #take this value from summary(surf.te) from nutritional landscape code above

LB=c(0,0), # lower bounds (origin)

UB=c(0.71627942,0.545), #upper bounds on the nutritional landscape (max on P/x axis , max on C/y axis)

xlab = "Protein Eaten (g)",

ylab = "Carbohydrate Eaten (g)",

triangularRegion=TRUE,

vertex1=c(0.069, 0.545),# vertex 1 (x,y)is the corner of the triangle near the far end of the carb (y) axis

vertex2=c(0.71627942, 0.08958410), # vertex 2 (x,y) is the corner of the triangle near the far end of the protein (x) axis

maximization=FALSE,#looks for the CR on the minimum (trough) within the landscape

outputPDFFile = "95CR - Dev Time Min - Females- WH6.pdf")

set.seed(7, kind="Wichmann-Hill")

out <- OptRegionTps(X=CR.DATA.DevTime.Females[,1:2], y=CR.DATA.DevTime.Females[,3], nosim = 2000,

lambda = 0.06085, #take this value from summary(surf.te) from nutritional landscape code above

LB=c(0,0), # lower bounds (origin)

UB=c(0.71627942,0.545), #upper bounds on the nutritional landscape (max on P/x axis , max on C/y axis)

xlab = "Protein Eaten (g)",

ylab = "Carbohydrate Eaten (g)",

triangularRegion=TRUE,

vertex1=c(0.069, 0.545),# vertex 1 (x,y)is the corner of the triangle near the far end of the carb (y) axis

vertex2=c(0.71627942, 0.08958410), # vertex 2 (x,y) is the corner of the triangle near the far end of the protein (x) axis

maximization=FALSE,#looks for the CR on the minimum (trough) within the landscape

outputPDFFile = "95CR - Dev Time Min - Females- WH7.pdf")

set.seed(8, kind="Wichmann-Hill")

out <- OptRegionTps(X=CR.DATA.DevTime.Females[,1:2], y=CR.DATA.DevTime.Females[,3], nosim = 2000,

lambda = 0.06085, #take this value from summary(surf.te) from nutritional landscape code above

LB=c(0,0), # lower bounds (origin)

UB=c(0.71627942,0.545), #upper bounds on the nutritional landscape (max on P/x axis , max on C/y axis)

xlab = "Protein Eaten (g)",

ylab = "Carbohydrate Eaten (g)",

triangularRegion=TRUE,

vertex1=c(0.069, 0.545),# vertex 1 (x,y)is the corner of the triangle near the far end of the carb (y) axis

vertex2=c(0.71627942, 0.08958410), # vertex 2 (x,y) is the corner of the triangle near the far end of the protein (x) axis

maximization=FALSE,#looks for the CR on the minimum (trough) within the landscape

outputPDFFile = "95CR - Dev Time Min - Females- WH8.pdf")

set.seed(9, kind="Wichmann-Hill")

out <- OptRegionTps(X=CR.DATA.DevTime.Females[,1:2], y=CR.DATA.DevTime.Females[,3], nosim = 2000,

lambda = 0.06085, #take this value from summary(surf.te) from nutritional landscape code above

LB=c(0,0), # lower bounds (origin)

UB=c(0.71627942,0.545), #upper bounds on the nutritional landscape (max on P/x axis , max on C/y axis)

xlab = "Protein Eaten (g)",

ylab = "Carbohydrate Eaten (g)",

triangularRegion=TRUE,

vertex1=c(0.069, 0.545),# vertex 1 (x,y)is the corner of the triangle near the far end of the carb (y) axis

vertex2=c(0.71627942, 0.08958410), # vertex 2 (x,y) is the corner of the triangle near the far end of the protein (x) axis

maximization=FALSE,#looks for the CR on the minimum (trough) within the landscape

outputPDFFile = "95CR - Dev Time Min - Females- WH9.pdf")

set.seed(10, kind="Wichmann-Hill")

out <- OptRegionTps(X=CR.DATA.DevTime.Females[,1:2], y=CR.DATA.DevTime.Females[,3], nosim = 2000,

lambda = 0.06085, #take this value from summary(surf.te) from nutritional landscape code above

LB=c(0,0), # lower bounds (origin)

UB=c(0.71627942,0.545), #upper bounds on the nutritional landscape (max on P/x axis , max on C/y axis)

xlab = "Protein Eaten (g)",

ylab = "Carbohydrate Eaten (g)",

triangularRegion=TRUE,

vertex1=c(0.069, 0.545),# vertex 1 (x,y)is the corner of the triangle near the far end of the carb (y) axis

vertex2=c(0.71627942, 0.08958410), # vertex 2 (x,y) is the corner of the triangle near the far end of the protein (x) axis

maximization=FALSE,#looks for the CR on the minimum (trough) within the landscape

outputPDFFile = "95CR - Dev Time Min - Females- WH10.pdf")

set.seed(11, kind="Wichmann-Hill")

out <- OptRegionTps(X=CR.DATA.DevTime.Females[,1:2], y=CR.DATA.DevTime.Females[,3], nosim = 2000,

lambda = 0.06085, #take this value from summary(surf.te) from nutritional landscape code above

LB=c(0,0), # lower bounds (origin)

UB=c(0.71627942,0.545), #upper bounds on the nutritional landscape (max on P/x axis , max on C/y axis)

xlab = "Protein Eaten (g)",

ylab = "Carbohydrate Eaten (g)",

triangularRegion=TRUE,

vertex1=c(0.069, 0.545),# vertex 1 (x,y)is the corner of the triangle near the far end of the carb (y) axis

vertex2=c(0.71627942, 0.08958410), # vertex 2 (x,y) is the corner of the triangle near the far end of the protein (x) axis

maximization=FALSE,#looks for the CR on the minimum (trough) within the landscape

outputPDFFile = "95CR - Dev Time Min - Females- WH11.pdf")

set.seed(12, kind="Wichmann-Hill")

out <- OptRegionTps(X=CR.DATA.DevTime.Females[,1:2], y=CR.DATA.DevTime.Females[,3], nosim = 2000,

lambda = 0.06085, #take this value from summary(surf.te) from nutritional landscape code above

LB=c(0,0), # lower bounds (origin)

UB=c(0.71627942,0.545), #upper bounds on the nutritional landscape (max on P/x axis , max on C/y axis)

xlab = "Protein Eaten (g)",

ylab = "Carbohydrate Eaten (g)",

triangularRegion=TRUE,

vertex1=c(0.069, 0.545),# vertex 1 (x,y)is the corner of the triangle near the far end of the carb (y) axis

vertex2=c(0.71627942, 0.08958410), # vertex 2 (x,y) is the corner of the triangle near the far end of the protein (x) axis

maximization=FALSE,#looks for the CR on the minimum (trough) within the landscape

outputPDFFile = "95CR - Dev Time Min - Females- WH12.pdf")

set.seed(13, kind="Wichmann-Hill")

out <- OptRegionTps(X=CR.DATA.DevTime.Females[,1:2], y=CR.DATA.DevTime.Females[,3], nosim = 2000,

lambda = 0.06085, #take this value from summary(surf.te) from nutritional landscape code above

LB=c(0,0), # lower bounds (origin)

UB=c(0.71627942,0.545), #upper bounds on the nutritional landscape (max on P/x axis , max on C/y axis)

xlab = "Protein Eaten (g)",

ylab = "Carbohydrate Eaten (g)",

triangularRegion=TRUE,

vertex1=c(0.069, 0.545),# vertex 1 (x,y)is the corner of the triangle near the far end of the carb (y) axis

vertex2=c(0.71627942, 0.08958410), # vertex 2 (x,y) is the corner of the triangle near the far end of the protein (x) axis

maximization=FALSE,#looks for the CR on the minimum (trough) within the landscape

outputPDFFile = "95CR - Dev Time Min - Females- WH13.pdf")

set.seed(14, kind="Wichmann-Hill")

out <- OptRegionTps(X=CR.DATA.DevTime.Females[,1:2], y=CR.DATA.DevTime.Females[,3], nosim = 2000,

lambda = 0.06085, #take this value from summary(surf.te) from nutritional landscape code above

LB=c(0,0), # lower bounds (origin)

UB=c(0.71627942,0.545), #upper bounds on the nutritional landscape (max on P/x axis , max on C/y axis)

xlab = "Protein Eaten (g)",

ylab = "Carbohydrate Eaten (g)",

triangularRegion=TRUE,

vertex1=c(0.069, 0.545),# vertex 1 (x,y)is the corner of the triangle near the far end of the carb (y) axis

vertex2=c(0.71627942, 0.08958410), # vertex 2 (x,y) is the corner of the triangle near the far end of the protein (x) axis

maximization=FALSE,#looks for the CR on the minimum (trough) within the landscape

outputPDFFile = "95CR - Dev Time Min - Females- WH14.pdf")

set.seed(15, kind="Wichmann-Hill")

out <- OptRegionTps(X=CR.DATA.DevTime.Females[,1:2], y=CR.DATA.DevTime.Females[,3], nosim = 2000,

lambda = 0.06085, #take this value from summary(surf.te) from nutritional landscape code above

LB=c(0,0), # lower bounds (origin)

UB=c(0.71627942,0.545), #upper bounds on the nutritional landscape (max on P/x axis , max on C/y axis)

xlab = "Protein Eaten (g)",

ylab = "Carbohydrate Eaten (g)",

triangularRegion=TRUE,

vertex1=c(0.069, 0.545),# vertex 1 (x,y)is the corner of the triangle near the far end of the carb (y) axis

vertex2=c(0.71627942, 0.08958410), # vertex 2 (x,y) is the corner of the triangle near the far end of the protein (x) axis

maximization=FALSE,#looks for the CR on the minimum (trough) within the landscape

outputPDFFile = "95CR - Dev Time Min - Females- WH15.pdf")

set.seed(16, kind="Wichmann-Hill")

out <- OptRegionTps(X=CR.DATA.DevTime.Females[,1:2], y=CR.DATA.DevTime.Females[,3], nosim = 2000,

lambda = 0.06085, #take this value from summary(surf.te) from nutritional landscape code above

LB=c(0,0), # lower bounds (origin)

UB=c(0.71627942,0.545), #upper bounds on the nutritional landscape (max on P/x axis , max on C/y axis)

xlab = "Protein Eaten (g)",

ylab = "Carbohydrate Eaten (g)",

triangularRegion=TRUE,

vertex1=c(0.069, 0.545),# vertex 1 (x,y)is the corner of the triangle near the far end of the carb (y) axis

vertex2=c(0.71627942, 0.08958410), # vertex 2 (x,y) is the corner of the triangle near the far end of the protein (x) axis

maximization=FALSE,#looks for the CR on the minimum (trough) within the landscape

outputPDFFile = "95CR - Dev Time Min - Females- WH16.pdf")

set.seed(17, kind="Wichmann-Hill")

out <- OptRegionTps(X=CR.DATA.DevTime.Females[,1:2], y=CR.DATA.DevTime.Females[,3], nosim = 2000,

lambda = 0.06085, #take this value from summary(surf.te) from nutritional landscape code above

LB=c(0,0), # lower bounds (origin)

UB=c(0.71627942,0.545), #upper bounds on the nutritional landscape (max on P/x axis , max on C/y axis)

xlab = "Protein Eaten (g)",

ylab = "Carbohydrate Eaten (g)",

triangularRegion=TRUE,

vertex1=c(0.069, 0.545),# vertex 1 (x,y)is the corner of the triangle near the far end of the carb (y) axis

vertex2=c(0.71627942, 0.08958410), # vertex 2 (x,y) is the corner of the triangle near the far end of the protein (x) axis

maximization=FALSE,#looks for the CR on the minimum (trough) within the landscape

outputPDFFile = "95CR - Dev Time Min - Females- WH17.pdf")

set.seed(18, kind="Wichmann-Hill")

out <- OptRegionTps(X=CR.DATA.DevTime.Females[,1:2], y=CR.DATA.DevTime.Females[,3], nosim = 2000,

lambda = 0.06085, #take this value from summary(surf.te) from nutritional landscape code above

LB=c(0,0), # lower bounds (origin)

UB=c(0.71627942,0.545), #upper bounds on the nutritional landscape (max on P/x axis , max on C/y axis)

xlab = "Protein Eaten (g)",

ylab = "Carbohydrate Eaten (g)",

triangularRegion=TRUE,

vertex1=c(0.069, 0.545),# vertex 1 (x,y)is the corner of the triangle near the far end of the carb (y) axis

vertex2=c(0.71627942, 0.08958410), # vertex 2 (x,y) is the corner of the triangle near the far end of the protein (x) axis

maximization=FALSE,#looks for the CR on the minimum (trough) within the landscape

outputPDFFile = "95CR - Dev Time Min - Females- WH18.pdf")

set.seed(19, kind="Wichmann-Hill")

out <- OptRegionTps(X=CR.DATA.DevTime.Females[,1:2], y=CR.DATA.DevTime.Females[,3], nosim = 2000,

lambda = 0.06085, #take this value from summary(surf.te) from nutritional landscape code above

LB=c(0,0), # lower bounds (origin)

UB=c(0.71627942,0.545), #upper bounds on the nutritional landscape (max on P/x axis , max on C/y axis)

xlab = "Protein Eaten (g)",

ylab = "Carbohydrate Eaten (g)",

triangularRegion=TRUE,

vertex1=c(0.069, 0.545),# vertex 1 (x,y)is the corner of the triangle near the far end of the carb (y) axis

vertex2=c(0.71627942, 0.08958410), # vertex 2 (x,y) is the corner of the triangle near the far end of the protein (x) axis

maximization=FALSE,#looks for the CR on the minimum (trough) within the landscape

outputPDFFile = "95CR - Dev Time Min - Females- WH19.pdf")

set.seed(20, kind="Wichmann-Hill")

out <- OptRegionTps(X=CR.DATA.DevTime.Females[,1:2], y=CR.DATA.DevTime.Females[,3], nosim = 2000,

lambda = 0.06085, #take this value from summary(surf.te) from nutritional landscape code above

LB=c(0,0), # lower bounds (origin)

UB=c(0.71627942,0.545), #upper bounds on the nutritional landscape (max on P/x axis , max on C/y axis)

xlab = "Protein Eaten (g)",

ylab = "Carbohydrate Eaten (g)",

triangularRegion=TRUE,

vertex1=c(0.069, 0.545),# vertex 1 (x,y)is the corner of the triangle near the far end of the carb (y) axis

vertex2=c(0.71627942, 0.08958410), # vertex 2 (x,y) is the corner of the triangle near the far end of the protein (x) axis

maximization=FALSE,#looks for the CR on the minimum (trough) within the landscape

outputPDFFile = "95CR - Dev Time Min - Females- WH20.pdf")

######### DEVELOPMENT TIME MODEL ###############

#Development time and Survival Analyses

\# Install all the packages listed in lines 7-22, either individually through the Tools menu, or with the shortcut code on line 6

#After all the changes to these stats, I have actually lost track which of these packages are being used in the code. Load em all anyways.

\# Make sure to change all directory paths to your own (e.g. line 24, or wherever write_xlsx is used)

\# Highlight all lines between 7 - 42, and then hit run before running any models below

install.packages(c("readxl","survival", "splines")) # complete the list for the packages that you need to install

library(readxl)

library(survival)

library(splines)

library(survminer)

library(car)#for 2 and 3 way Anova

library(writexl)

library(ggplot2)

library(MASS)

library(multcomp)

library(MASS) #for AIC model selection

library(emmeans)

library(xlsx)

library(pspline)

library(effects)

library(MuMIn)

library(ggpubr)

DATA <- read_excel(".../Muzzatti.et.al.2024.Data.xlsx", sheet = "dev and survival")

#change variables to the correct types (numeric or factor)

DATA$Cellulose=as.factor(DATA$Cellulose)

DATA$P.C=as.factor(DATA$P.C)

DATA$Status.Adult=as.numeric(DATA$Status.Adult)

DATA$Status.Death=as.numeric(DATA$Status.Death)

#This sets the order of the P:C and Cellulose levels, making them ordinal categorical.

\# The first level listed for each automatically becomes the reference category for the model

\# If for some reason you wish to change the reference category in the model, do it by changing the order of the levels here

DATA$P.C <- factor(DATA$P.C, levels = c('1:8', '1:5','1:3','1:1','3:1','5:1','8:1'))

DATA$Cellulose<-factor(DATA$Cellulose, levels=c("L","M","H"))

DATA<-as.data.frame(DATA) # spreadsheet must be in a data frame for cox model to work, data frame is just how R stores a spreadsheet

str(DATA)

View(DATA)

\## Fit the Full model in prep for model selection

m1=coxph(Surv(Dev.Time,Status.Adult)~P.C*Cellulose,data=DATA) # Full model, includes interaction

summary(m1)

\# Test assumptions of the full model

\# no need to test for assumption of linearity because there are no continuous terms in the model

zfit.m1<-cox.zph(m1,transform = 'identity', terms =FALSE)

zfit.m1 # no significant terms = meets assumption of proportional hazards over time (i.e. coef estimates do not change with time)

ggcoxdiagnostics(m1, type = "deviance",linear.predictions = FALSE, ggtheme = theme_bw()) # No influential outliers

#Troubleshooting: If you get an error for this plot (or any plot for R studio), stretch the plot window and clear plot history and retry. There's a bug.

\## MODEL SELECTION - DEVELOPMENT TIME

\# Symonds and Moussali 2011 gives a very concise overview of the methods for model selection

\# Produce a candidate set of models, ranked by AIC from lowest to highest.

\# Candidate set = every possible combo of predictors, from the null model with just intercept up to the full model

\# Which information criterion to use for the ranking? AIC or AICc? Your model has n=928 and k=4 (4 terms in the full model,

\# including intercept), n/k = 232 which is > 40 (the standard cutoff), so use AIC because not a small sample size.

\# "Weight" = Akaike weight, or the probability that a given model is the best approximating model for your data.

options(na.action= "na.fail") # necessary for dredge to work

m1.Global.mod.eval<-dredge(m1,rank=AIC,evaluate = TRUE)

m1.Global.mod.eval # this is your CANDIDATE SET. 5 models total, where + indicates whether that term is in the model. Cll = cellulose

#reject models with delta AIC < 4

\# A cutoff of 4 ensures with 95% confidence (approximately) that you have included the best approximating

\# model - but see symonds & Moussali 2011 for discussion on what delta AIC cutoffs are commonly used and why

m1.subset4<-subset(m1.Global.mod.eval, delta < 4,recalc.weights = TRUE,recalc.delta=TRUE)# Weight column recalculated to sum to 1

m1.subset4 #Show the models with AIC<4, penultimate confidence set

#Further reject models that are just more complex versions of other higher ranked models (nesting approach - Richards 2008, Richards et al 2011)

#I.e. Model 8 is just a more complex version of 4, which has a lower AIC, so we reject model 8.

nested(m1.subset4) # gives a T/F list whether model X has other models 'nested' within it

nst <- nested(m1.subset4, indices = "row")

nst # print model "x" and all models nested within it

m1.subset4$nested <- sapply(nst, paste, collapse = ",")

m1.subset4 #Original model selection table with the nested models indicated

m1.nest.reduced.subset4<-subset(m1.subset4, !nested(.)) # Automatically filter out overly complex models according to the "nesting selection rule"

m1.nest.reduced.subset4 #THIS TABLE IS YOUR CONFIDENCE SET. Weight column recalculated to sum to 1

#In this case, your top ranked model has unequivocal support (weight = 1)

\## FINAL MODEL - DEVELOPMENT TIME

#Best approximating model from model selection (ie has all the support (weight = 1) after rejecting models above)

m1.Best.Approx.Model<- (get.models(m1.nest.reduced.subset4, 1)[[1]])

m1.Best.Approx.Model

#Re-check model assumptions

zfit.m1Best<-cox.zph(m1.Best.Approx.Model,transform = 'identity', terms =FALSE)

zfit.m1Best # no significant terms = meets assumption of proportional hazards over time (i.e. coef estimates do not change with time)

ggcoxdiagnostics(m1.Best.Approx.Model, type = "deviance",linear.predictions = FALSE, ggtheme = theme_bw())#No influential outliers

#Overall effects of each model parameter - might not be necessary to present this in your results

Anova(m1.Best.Approx.Model, type = 2) # use type 2 when no interactions significant

\# Parameter/Hazard ratio estimates

summary(m1.Best.Approx.Model)

\## HAZARD RATIOS FOR ALL 21 CATEGORIES - DEVELOPMENT TIME

\# This data is needed later for the PLOTS and CONTRASTS

\# Use the emmeans package to get HRs for each of the 21 categories of P:C x Cellulose

\# Even though there is no interaction in the final model, we ask for predicted fits/means

\# for each of the 21 categories by specifying the interaction P:C x Cellulose.

All.21.HRs.m1<-emmeans(m1.Best.Approx.Model, ~P.C:Cellulose, type = "response")# You know that "response" is HR scale because 1:8 = 1, not 0

All.21.HRs.m1 # For the few categories that are shown in the model summary tables, these HRs match

write.xlsx(as.data.frame(All.21.HRs.m1),"...")

#Plot for this data is several sections below

#Proof/Check that these HRs match what is being shown in the model summary tables

summary(m1.Best.Approx.Model)

\# E.g. compare CelluloseM HR and CIs in the model summary table with 1:8-Med in the All.21.HRs table

\# Both are: HR= 0.9442, LwrCI = 0.6697, UprCI= 1.331

\# E.g. compare P.C1:5 HR and CIs in the model summary table with the 1:5-Low in the All.21.HRs table

\# Both are: HR= 1.97288, LwrCI = 0.9145, UprCI= 4.256

\## HAZARD RATIOS - AVERAGED ACROSS CATEGORIES - DEVELOPMENT TIME

Average.PC.HRs.m1<-emmeans(m1.Best.Approx.Model, ~P.C, type = "response")

Average.PC.HRs.m1 # HRs averaged over levels of cellulose. Averaging is performed on the Coef scale, then converted to HRs

write_xlsx(as.data.frame(Average.PC.HRs.m1),"C:\\Users\\Sarah Harrison\\Desktop\\Matt\\Survival Analysis Code for Matt\\DevTime - Average.PC.HRs.xlsx")

write.xlsx(as.data.frame(Average.PC.HRs.m1),"C:\\Users\\Work\\OneDrive - Carleton University\\PhD\\Chapter 3 - 21 diets\\Experiment 1 - no choice\\From Sarah_12-05\\Development Time Model\\DevTime - Average.PC.HRs.xlsx") #for Matt's directory

Average.Cellulose.HRs.m1<-emmeans(m1.Best.Approx.Model, ~Cellulose, type = "response")

Average.Cellulose.HRs.m1 # HRs averaged over levels of P:C. Averaging is performed on the Coef scale, then converted to HRs

write_xlsx(as.data.frame(Average.Cellulose.HRs.m1),"C:\\Users\\Sarah Harrison\\Desktop\\Matt\\Survival Analysis Code for Matt\\DevTime - Average.Cellulose.HRs.xlsx")

write.xlsx(as.data.frame(Average.Cellulose.HRs.m1),"C:\\Users\\Work\\OneDrive - Carleton University\\PhD\\Chapter 3 - 21 diets\\Experiment 1 - no choice\\From Sarah_12-05\\Development Time Model\\DevTime - Average.Cellulose.HRs.xlsx") #for Matt's directory

#As a double check that these averages are correct: 1) calculate coefficients for all 21 categories,

\# 2) manually calculate averages across the relevant categories, 3) then convert to HR with exp(coef)

\# You cannot calculate averages on the hazard ratio scale. Must be on the coefficient scale, and then convert to hazard ratio

All.21.Coefs.m1<-emmeans(m1.Best.Approx.Model, ~P.C:Cellulose)# You know that "response" is Coef scale because 1:8 = 0, not 1

All.21.Coefs.m1 # emmean = coefficient

write_xlsx(as.data.frame(All.21.Coefs.m1),"...")

\## CONTRASTS - DEVELOPMENT TIME

\# First assign vector coordinates for the different rows of the All.21.HRs table from previous section in order

\# to identify the rows to contrast.

PC3.1L<-c(1,0,0,0,0,0,0,0,0,0,0,0,0,0,0,0,0,0,0,0,0)

PC1.8L<-c(0,1,0,0,0,0,0,0,0,0,0,0,0,0,0,0,0,0,0,0,0)

PC1.5L<-c(0,0,1,0,0,0,0,0,0,0,0,0,0,0,0,0,0,0,0,0,0)

PC1.3L<-c(0,0,0,1,0,0,0,0,0,0,0,0,0,0,0,0,0,0,0,0,0)

PC1.1L<-c(0,0,0,0,1,0,0,0,0,0,0,0,0,0,0,0,0,0,0,0,0)

PC5.1L<-c(0,0,0,0,0,1,0,0,0,0,0,0,0,0,0,0,0,0,0,0,0)

PC8.1L<-c(0,0,0,0,0,0,1,0,0,0,0,0,0,0,0,0,0,0,0,0,0)

PC3.1M<-c(0,0,0,0,0,0,0,1,0,0,0,0,0,0,0,0,0,0,0,0,0)

PC1.8M<-c(0,0,0,0,0,0,0,0,1,0,0,0,0,0,0,0,0,0,0,0,0)

PC1.5M<-c(0,0,0,0,0,0,0,0,0,1,0,0,0,0,0,0,0,0,0,0,0)

PC1.3M<-c(0,0,0,0,0,0,0,0,0,0,1,0,0,0,0,0,0,0,0,0,0)

PC1.1M<-c(0,0,0,0,0,0,0,0,0,0,0,1,0,0,0,0,0,0,0,0,0)

PC5.1M<-c(0,0,0,0,0,0,0,0,0,0,0,0,1,0,0,0,0,0,0,0,0)

PC8.1M<-c(0,0,0,0,0,0,0,0,0,0,0,0,0,1,0,0,0,0,0,0,0)

PC3.1H<-c(0,0,0,0,0,0,0,0,0,0,0,0,0,0,1,0,0,0,0,0,0)

PC1.8H<-c(0,0,0,0,0,0,0,0,0,0,0,0,0,0,0,1,0,0,0,0,0)

PC1.5H<-c(0,0,0,0,0,0,0,0,0,0,0,0,0,0,0,0,1,0,0,0,0)

PC1.3H<-c(0,0,0,0,0,0,0,0,0,0,0,0,0,0,0,0,0,1,0,0,0)

PC1.1H<-c(0,0,0,0,0,0,0,0,0,0,0,0,0,0,0,0,0,0,1,0,0)

PC5.1H<-c(0,0,0,0,0,0,0,0,0,0,0,0,0,0,0,0,0,0,0,1,0)

PC8.1H<-c(0,0,0,0,0,0,0,0,0,0,0,0,0,0,0,0,0,0,0,0,1)

\# Perform contrasts to the reference category (1:8-Low)

All.HRs.Contrasts.to.RefCat.m1<- contrast(All.21.HRs.m1,adjust="none", CI=TRUE, method=list( (PC1.8L-PC1.8L),# Obviously = 1, because comparing to itself

(PC1.5L-PC1.8L),#Matches line P.C1:5 in model summary

(PC1.3L-PC1.8L),#Matches line P.C1:3 in model summary

(PC1.1L-PC1.8L),#Matches line P.C1:1 in model summary

(PC3.1L-PC1.8L),#Matches line P.C3:1 in model summary

(PC5.1L-PC1.8L),#Matches line P.C5:1 in model summary

(PC8.1L-PC1.8L),#Matches line P.C8:1 in model summary

(PC1.8M-PC1.8L),#Matches line CelluloseM in model summary

(PC1.5M-PC1.8L),

(PC1.3M-PC1.8L),

(PC1.1M-PC1.8L),

(PC3.1M-PC1.8L),

(PC5.1M-PC1.8L),

(PC8.1M-PC1.8L),

(PC1.8H-PC1.8L),#Matches line CelluloseH in model summary

(PC1.5H-PC1.8L),

(PC1.3H-PC1.8L),

(PC1.1H-PC1.8L),

(PC3.1H-PC1.8L),

(PC5.1H-PC1.8L),

(PC8.1H-PC1.8L)))

#Ref category of 3.1PC

All.HRs.Contrasts.to.RefCat.m1<- contrast(All.21.HRs.m1,adjust="none", CI=TRUE, method=list(

+	(PC1.8L-PC3.1L),

+	(PC1.5L-PC3.1L),

+	(PC1.3L-PC3.1L),

+	(PC1.1L-PC3.1L),

+	(PC3.1L-PC3.1L),

+	(PC5.1L-PC3.1L),

+	(PC8.1L-PC3.1L),

+	(PC1.8M-PC3.1L),

+	(PC1.5M-PC3.1L),

+	(PC1.3M-PC3.1L),

+	(PC1.1M-PC3.1L),

+	(PC3.1M-PC3.1L),

+	(PC5.1M-PC3.1L),

+	(PC8.1M-PC3.1L),

+	(PC1.8H-PC3.1L),

+	(PC1.5H-PC3.1L),

+	(PC1.3H-PC3.1L),

+	(PC1.1H-PC3.1L),

+	(PC3.1H-PC3.1L),

+	(PC5.1H-PC3.1L),

+	(PC8.1H-PC3.1L)))

All.HRs.Contrasts.to.RefCat.m1

\# Contrasts are labeled using vector coordinates. But the contrasts are in the same order as those listed

\# in method = list, so use that to properly label the contrasts in this table

\# As a double check that these contrasts are indeed correct, compare the relevant contrasts with the exp(coef), z, and p stats

\# in the model summary tables - see notes beside individual contrasts above (they do match)

#Write to excel

write_xlsx(as.data.frame(All.HRs.Contrasts.to.RefCat.m1),"...")

\# ALL pairwise contrasts between HRs

All.HR.Contrasts.m1<-contrast(All.21.HRs.m1, "pairwise", adjust = "none")

options(max.print = 5000)#increase max print so that table isnt cut off

All.HR.Contrasts.m1 #df = inf is normal for a cox model. I cant remember why though

#Write to excel

write_xlsx(as.data.frame(All.HR.Contrasts.m1),"...")

\## PLOT - DEVELOPMENT TIME

#Import the All.21.HRs excel doc. No changes are needed before importing

plotdata<- read_excel("...")

plotdata<- read_excel("...") #new directory

plotdata<-as.data.frame(plotdata)

str(plotdata)

#Set the order of different levels of the diet factors so they appear in the correct order in the plot

plotdata$Cellulose <- factor(plotdata$Cellulose, levels = c('L', 'M', 'H'))

plotdata$P.C <- factor(plotdata$P.C, levels = c('1:8','1:5','1:3','1:1','3:1','5:1','8:1'))

\# A an adjustment to use later to make the plot elements not overlap

pd <- position_dodge(0.6) #increase or decrease value to change the distance between the plot elements

cols <- c("L" = "#E69F00", "M" = "#009E73", "H" = "#CC79A7")

DT.Plot<-ggplot(plotdata, aes(x=P.C,y=response, colour = Cellulose, group=Cellulose))+

geom_errorbar(aes(ymin=asymp.LCL, ymax=asymp.UCL), width=0.2,lwd=0.9,position=pd, size = 1)+

geom_line(position=pd, size = 1.2)+

geom_point(position=pd, size = 2)+

geom_hline(yintercept = 1, size = 0.5, lty =2, colour = "black")+ # horizontal line showing where risk of death switches from increasing to decreasing

#scale_x_discrete(labels=c('label1', 'label2', 'label3', ...)+# this code could be used to change x axis labels

scale_color_manual(values = cols, labels=c('Low', 'Medium', 'High'))+ #Change cellulose legend labels here

scale_y_continuous(limits = c(0, 1.6),breaks=c(0,0.2,0.4,0.6,0.8,1.0,1.2,1.4,1.6))+ # Change y axis labels here

labs(x = "P:C",y = " Development Time Hazard Ratio", colour = "Cellulose") # Change x and y axis titles here

DT.Plot+theme(panel.border = element_rect(colour = "black", fill=NA, size=0.5),

panel.grid.major = element_blank(),

panel.grid.minor = element_blank(),

panel.background = element_blank(),

axis.line = element_line(colour = "black"),

axis.text=element_text(size=16),

axis.title=element_text(size=16),

legend.key=element_blank(),

legend.text = element_text(size=16),

legend.title = element_text(size=16))

\# troubleshooting: if you get a margins or graphics state error, clear the plot history and increase the size of the plot area

#saves the last plot that was created, but there is a time limit - rerun the plot code if this doesnt work

ggsave("DevTimePlot.jpeg", width = 15, height = 10, units = "cm", dpi = 500, path = "...")

\# Extension can be altered (e.g. .pdf, .jpeg, .tiff etc)

\# If the proportions/sizing of plot labels seems off, try changing the width and height here and then re-save

\# Change the dpi to get more crisp lines.

######### SURVIVAL TIME MODEL ###############

\## Fit the Full model in prep for model selection

m2=coxph(Surv(Survival.Time,Status.Death)~P.C*Cellulose,data=DATA) # Full model, includes interaction

summary(m2)

\# Test assumptions of the full model

\# no need to test for assumption of linearity because there are no continuous terms in the model

ggcoxdiagnostics(m2, type = "deviance",linear.predictions = FALSE, ggtheme = theme_bw()) # No influential outliers? I think. Its a bit wavey, but not bad

#Troubleshooting: If you get an error for this plot (or any plot for R studio), stretch the plot window and clear plot history and retry. There's a bug.

zfit.m2.Full<-cox.zph(m2, transform = 'identity', terms = FALSE) #Identity means no time axis transformation, terms = FALSE separates the cellulose coef terms (because categorical)

zfit.m2.Full # significant terms = violation of assumption of proportional hazards.

\# Overall, the violation of PH in the full model are not bad (see next 3 plots).

\# Problem is with CelluloseH coef, and then that is causing P.C1:1:CelluloseH term to be in violation as well

#Plot of CelluloseM coefficient vs Time (linear/identity/no transformation)

plot(zfit.m2.Full[7], lwd=1.5)

abline(0,0, col=2)# adds a red line at zero

abline(h= m2$coef[7], col=3, lwd=2, lty=2)# Green Dashed is the singular coef estimate for CelluloseM from m2 model summary

#Not a terrible fit, green line stays within black line CIs

#Plot of CelluloseH coefficient vs Time (linear/identity/no transformation)

plot(zfit.m2.Full[8], lwd=1.5)

abline(0,0, col=2)# adds a red line at zero

abline(h= m2$coef[8], col=3, lwd=2, lty=2)# Green Dashed is the singular coef estimate for CelluloseH from m2 model summary

#Not a terrible fit, green line stays within black line CIs most of the time, except for near the start

#Plot of P.C1:1:CelluloseH coefficient vs Time (linear/identity/no transformation)

plot(zfit.m2.Full[14], lwd=1.5)

abline(0,0, col=2)# adds a red line at zero

abline(h= m2$coef[14], col=3, lwd=2, lty=2)# Green Dashed is the singular coef estimate for P.C1:1:CelluloseH from m2 model summary

#Not a terrible fit at all, green line stays within black line CIs

\## MODEL SELECTION - SURVIVAL TIME

options(na.action= "na.fail") # necessary for dredge to work

m2.Global.mod.eval<-dredge(m2,rank=AIC,evaluate = TRUE)

m2.Global.mod.eval # this is your CANDIDATE SET. 5 models total, where + indicates whether that term is in the model. Cll = cellulose

#reject models with delta AIC < 4

\# A cutoff of 4 ensures with 95% confidence (approximately) that you have included the best approximating

\# model - but see symonds & Moussali 2011 for discussion on what delta AIC cutoffs are commonly used and why

m2.subset4<-subset(m2.Global.mod.eval, delta < 4,recalc.weights = TRUE,recalc.delta=TRUE)# Weight column recalculated to sum to 1

m2.subset4 #Show the models with AIC<4, this is your CONFIDENCE SET

\# In this case, your top ranked model of your confidence set has unequivocal support (weight = 1)

\# No need to proceed with the nesting approach because there's only 1 model

#Best approximating model from model selection (ie has all the support (weight = 1) after rejecting models above)

m2.Best.Approx.Model<- (get.models(m2.subset4, 1)[[1]])

m2.Best.Approx.Model

sw(m2.Global.mod.eval)

#Re-check model assumptions

ggcoxdiagnostics(m2.Best.Approx.Model, type = "deviance",linear.predictions = FALSE, ggtheme = theme_bw())#No outliers I think, But a little wavy

zfit.m2.Best<-cox.zph(m2.Best.Approx.Model, transform = 'identity', terms = FALSE)

zfit.m2.Best # still there are non proportional hazards for celluloseH (i.e. the coef changes over time)

#Even though the problem is only with CelluloseH, the time transformation gets applied

\# to both cellulose coefficients, so need to look at both CelluloseM and CelluloseH

#Plot of CelluloseM coefficient vs Time (linear/identity/no transformation)

plot(zfit.m2.Best[1], lwd=1.5)

abline(0,0, col=2)# adds a red line at zero

abline(h= m2$coef[1], col=3, lwd=2, lty=2)# Green Dashed is the singular coef estimate for CelluloseM from m2 model summary

#Not a terrible fit, green line stays within black line CIs

#Plot of CelluloseH coefficient vs Time (linear/identity/no transformation)

plot(zfit.m2.Best[2], lwd=1.5)

abline(0,0, col=2)# adds a red line at zero

abline(h= m2$coef[2], col=3, lwd=2, lty=2)# Green Dashed is the singular coef estimate for CelluloseM from m2 model summary

#Terrible fit. Green line goes outside CIs a lot

\## Add time stratification to the best approximating model above

\# See section 4.1 of this vingnette, but you will likely need to read the whole document to understand that section:

\# https://cran.r-project.org/web/packages/survival/vignettes/timedep.pdf

#Time is NOW cut into 3 different sections: 0-7, 7-14, 14-153

DATA2<-survSplit(Surv(Survival.Time,Status.Death)~., data=DATA, cut=c(7,14), episode="Time.Interval", id="NEW.ID")

\# Adds some new columns to the end of DATA

DATA2[c("NEW.ID", "ID", "tstart", "Survival.Time","Status.Death", "Time.Interval", "P.C", "Cellulose")]

print(DATA2)

DATA2<-as.data.frame(DATA2)

#Set variables to the correct type, and order levels

DATA2$Time.Interval=as.factor(DATA2$Time.Interval)

DATA2$P.C <- factor(DATA2$P.C, levels = c('3:1', '1:8','1:5','1:3','1:1','5:1','8:1')) #remember, change first ratio to be whichever ref category you want

DATA2$Cellulose<-factor(DATA2$Cellulose, levels=c("H","M","L")) # had to reverse the order so that L category was reference

str(DATA2)

DATA2$Time <- strata(DATA2$Time.Interval)

\###### FINAL MODEL - SURVIVAL TIME, Best approx model stratified by time

\## Now add the time interval variable into the model as an interaction with Cellulose

\# strata() is for nuisance variables whose effects you dont care about (ie you dont care about the relationship between survival and Time.Interval itself)

m3.Strata.Best.Approx.model<- coxph(Surv(tstart,Survival.Time,Status.Death)~P.C+(Cellulose:strata(Time)), data=DATA2) #dont forget to add tstart inside Surv()

m3.Strata.Best.Approx.model<- coxph(Surv(tstart,Survival.Time,Status.Death)~P.C+(Cellulose:Time), data=DATA2) #dont forget to add tstart inside Surv()

m3.Strata.Best.Approx.model <- coxph(Surv(tstart, Survival.Time, Status.Death) ~ P.C + Cellulose + Time + P.C:Cellulose + P.C:Time + Cellulose:Time, data = DATA2)

m3.Strata.Best.Approx.model

#Re-check proportional hazards assumptions

cox.zph(m3.Strata.Best.Approx.model, transform = "identity", terms = FALSE)# No significant violations of proportional hazards assumption

#Re-Check assumption of no influential outliers

ggcoxdiagnostics(m3.Strata.Best.Approx.model, type = "deviance",linear.predictions = FALSE, ggtheme = theme_bw()) # No influential outliers

#Overall effects of each model parameter - might not be necessary to present this in your results

Anova(m3.Strata.Best.Approx.model)

\# A quick double check that we'd get the same result had we done model selection on the time stratified full model

\# Model fails to converge - not enough degrees of freedom when we include P:C X Cellulose?

\# So we were right to wait to do the time stratification on the simplified model

\# m4 is the same as m3

m4.Strata.Full <- coxph(Surv(tstart,Survival.Time,Status.Death)~P.C*(Cellulose:strata(Time.Interval)), data=DATA2)

options(na.action= "na.fail")

m4.Strata.Full.mod.eval<-dredge(m4.Strata.Full,rank=AIC,evaluate = TRUE)

m4.Strata.Full.mod.eval

m4.subset4<-subset(m4.Strata.Full.mod.eval, delta < 4,recalc.weights = TRUE, recalc.delta=TRUE)

m4.subset4

m4.Strata.Best<- (get.models(m4.subset4, 1)[[1]])

m4.Strata.Best

#All 63 HAZARD RATIOS - SURVIVAL TIME - for all combinations of P:C, Cellulose and Time.Interval

#See same section for development time model for explanations

All.HRs.Strata.m3 <- emmeans(m3.Strata.Best.Approx.model, ~P.C:(Cellulose:Time), type = "response", data= DATA2) # You know that "response" is HR scale because 1:8L = 1, not 0

All.HRs.Strata.m3 <- emmeans(

m3.Strata.Best.Approx.model,

~ P.C:Cellulose:Time,

type = "response",

data = DATA2)

All.HRs.Strata.m3 # For the few categories that are shown in the model summary tables, these HRs match

\## CONTRASTS - SURVIVAL TIME

PC3.1H1<-c(1,0,0,0,0,0,0,0,0,0,0,0,0,0,0,0,0,0,0,0,0,0,0,0,0,0,0,0,0,0,0,0,0,0,0,0,0,0,0,0,0,0,0,0,0,0,0,0,0,0,0,0,0,0,0,0,0,0,0,0,0,0,0)

PC1.8H1<-c(0,1,0,0,0,0,0,0,0,0,0,0,0,0,0,0,0,0,0,0,0,0,0,0,0,0,0,0,0,0,0,0,0,0,0,0,0,0,0,0,0,0,0,0,0,0,0,0,0,0,0,0,0,0,0,0,0,0,0,0,0,0,0)

PC1.5H1<-c(0,0,1,0,0,0,0,0,0,0,0,0,0,0,0,0,0,0,0,0,0,0,0,0,0,0,0,0,0,0,0,0,0,0,0,0,0,0,0,0,0,0,0,0,0,0,0,0,0,0,0,0,0,0,0,0,0,0,0,0,0,0,0)

PC1.3H1<-c(0,0,0,1,0,0,0,0,0,0,0,0,0,0,0,0,0,0,0,0,0,0,0,0,0,0,0,0,0,0,0,0,0,0,0,0,0,0,0,0,0,0,0,0,0,0,0,0,0,0,0,0,0,0,0,0,0,0,0,0,0,0,0)

PC1.1H1<-c(0,0,0,0,1,0,0,0,0,0,0,0,0,0,0,0,0,0,0,0,0,0,0,0,0,0,0,0,0,0,0,0,0,0,0,0,0,0,0,0,0,0,0,0,0,0,0,0,0,0,0,0,0,0,0,0,0,0,0,0,0,0,0)

PC5.1H1<-c(0,0,0,0,0,1,0,0,0,0,0,0,0,0,0,0,0,0,0,0,0,0,0,0,0,0,0,0,0,0,0,0,0,0,0,0,0,0,0,0,0,0,0,0,0,0,0,0,0,0,0,0,0,0,0,0,0,0,0,0,0,0,0)

PC8.1H1<-c(0,0,0,0,0,0,1,0,0,0,0,0,0,0,0,0,0,0,0,0,0,0,0,0,0,0,0,0,0,0,0,0,0,0,0,0,0,0,0,0,0,0,0,0,0,0,0,0,0,0,0,0,0,0,0,0,0,0,0,0,0,0,0)

PC3.1M1<-c(0,0,0,0,0,0,0,1,0,0,0,0,0,0,0,0,0,0,0,0,0,0,0,0,0,0,0,0,0,0,0,0,0,0,0,0,0,0,0,0,0,0,0,0,0,0,0,0,0,0,0,0,0,0,0,0,0,0,0,0,0,0,0)

PC1.8M1<-c(0,0,0,0,0,0,0,0,1,0,0,0,0,0,0,0,0,0,0,0,0,0,0,0,0,0,0,0,0,0,0,0,0,0,0,0,0,0,0,0,0,0,0,0,0,0,0,0,0,0,0,0,0,0,0,0,0,0,0,0,0,0,0)

PC1.5M1<-c(0,0,0,0,0,0,0,0,0,1,0,0,0,0,0,0,0,0,0,0,0,0,0,0,0,0,0,0,0,0,0,0,0,0,0,0,0,0,0,0,0,0,0,0,0,0,0,0,0,0,0,0,0,0,0,0,0,0,0,0,0,0,0)

PC1.3M1<-c(0,0,0,0,0,0,0,0,0,0,1,0,0,0,0,0,0,0,0,0,0,0,0,0,0,0,0,0,0,0,0,0,0,0,0,0,0,0,0,0,0,0,0,0,0,0,0,0,0,0,0,0,0,0,0,0,0,0,0,0,0,0,0)

PC1.1M1<-c(0,0,0,0,0,0,0,0,0,0,0,1,0,0,0,0,0,0,0,0,0,0,0,0,0,0,0,0,0,0,0,0,0,0,0,0,0,0,0,0,0,0,0,0,0,0,0,0,0,0,0,0,0,0,0,0,0,0,0,0,0,0,0)

PC5.1M1<-c(0,0,0,0,0,0,0,0,0,0,0,0,1,0,0,0,0,0,0,0,0,0,0,0,0,0,0,0,0,0,0,0,0,0,0,0,0,0,0,0,0,0,0,0,0,0,0,0,0,0,0,0,0,0,0,0,0,0,0,0,0,0,0)

PC8.1M1<-c(0,0,0,0,0,0,0,0,0,0,0,0,0,1,0,0,0,0,0,0,0,0,0,0,0,0,0,0,0,0,0,0,0,0,0,0,0,0,0,0,0,0,0,0,0,0,0,0,0,0,0,0,0,0,0,0,0,0,0,0,0,0,0)

PC3.1L1<-c(0,0,0,0,0,0,0,0,0,0,0,0,0,0,1,0,0,0,0,0,0,0,0,0,0,0,0,0,0,0,0,0,0,0,0,0,0,0,0,0,0,0,0,0,0,0,0,0,0,0,0,0,0,0,0,0,0,0,0,0,0,0,0)

PC1.8L1<-c(0,0,0,0,0,0,0,0,0,0,0,0,0,0,0,1,0,0,0,0,0,0,0,0,0,0,0,0,0,0,0,0,0,0,0,0,0,0,0,0,0,0,0,0,0,0,0,0,0,0,0,0,0,0,0,0,0,0,0,0,0,0,0)

PC1.5L1<-c(0,0,0,0,0,0,0,0,0,0,0,0,0,0,0,0,1,0,0,0,0,0,0,0,0,0,0,0,0,0,0,0,0,0,0,0,0,0,0,0,0,0,0,0,0,0,0,0,0,0,0,0,0,0,0,0,0,0,0,0,0,0,0)

PC1.3L1<-c(0,0,0,0,0,0,0,0,0,0,0,0,0,0,0,0,0,1,0,0,0,0,0,0,0,0,0,0,0,0,0,0,0,0,0,0,0,0,0,0,0,0,0,0,0,0,0,0,0,0,0,0,0,0,0,0,0,0,0,0,0,0,0)

PC1.1L1<-c(0,0,0,0,0,0,0,0,0,0,0,0,0,0,0,0,0,0,1,0,0,0,0,0,0,0,0,0,0,0,0,0,0,0,0,0,0,0,0,0,0,0,0,0,0,0,0,0,0,0,0,0,0,0,0,0,0,0,0,0,0,0,0)

PC5.1L1<-c(0,0,0,0,0,0,0,0,0,0,0,0,0,0,0,0,0,0,0,1,0,0,0,0,0,0,0,0,0,0,0,0,0,0,0,0,0,0,0,0,0,0,0,0,0,0,0,0,0,0,0,0,0,0,0,0,0,0,0,0,0,0,0)

PC8.1L1<-c(0,0,0,0,0,0,0,0,0,0,0,0,0,0,0,0,0,0,0,0,1,0,0,0,0,0,0,0,0,0,0,0,0,0,0,0,0,0,0,0,0,0,0,0,0,0,0,0,0,0,0,0,0,0,0,0,0,0,0,0,0,0,0)

PC3.1H2<-c(0,0,0,0,0,0,0,0,0,0,0,0,0,0,0,0,0,0,0,0,0,1,0,0,0,0,0,0,0,0,0,0,0,0,0,0,0,0,0,0,0,0,0,0,0,0,0,0,0,0,0,0,0,0,0,0,0,0,0,0,0,0,0)

PC1.8H2<-c(0,0,0,0,0,0,0,0,0,0,0,0,0,0,0,0,0,0,0,0,0,0,1,0,0,0,0,0,0,0,0,0,0,0,0,0,0,0,0,0,0,0,0,0,0,0,0,0,0,0,0,0,0,0,0,0,0,0,0,0,0,0,0)

PC1.5H2<-c(0,0,0,0,0,0,0,0,0,0,0,0,0,0,0,0,0,0,0,0,0,0,0,1,0,0,0,0,0,0,0,0,0,0,0,0,0,0,0,0,0,0,0,0,0,0,0,0,0,0,0,0,0,0,0,0,0,0,0,0,0,0,0)

PC1.3H2<-c(0,0,0,0,0,0,0,0,0,0,0,0,0,0,0,0,0,0,0,0,0,0,0,0,1,0,0,0,0,0,0,0,0,0,0,0,0,0,0,0,0,0,0,0,0,0,0,0,0,0,0,0,0,0,0,0,0,0,0,0,0,0,0)

PC1.1H2<-c(0,0,0,0,0,0,0,0,0,0,0,0,0,0,0,0,0,0,0,0,0,0,0,0,0,1,0,0,0,0,0,0,0,0,0,0,0,0,0,0,0,0,0,0,0,0,0,0,0,0,0,0,0,0,0,0,0,0,0,0,0,0,0)

PC5.1H2<-c(0,0,0,0,0,0,0,0,0,0,0,0,0,0,0,0,0,0,0,0,0,0,0,0,0,0,1,0,0,0,0,0,0,0,0,0,0,0,0,0,0,0,0,0,0,0,0,0,0,0,0,0,0,0,0,0,0,0,0,0,0,0,0)

PC8.1H2<-c(0,0,0,0,0,0,0,0,0,0,0,0,0,0,0,0,0,0,0,0,0,0,0,0,0,0,0,1,0,0,0,0,0,0,0,0,0,0,0,0,0,0,0,0,0,0,0,0,0,0,0,0,0,0,0,0,0,0,0,0,0,0,0)

PC3.1M2<-c(0,0,0,0,0,0,0,0,0,0,0,0,0,0,0,0,0,0,0,0,0,0,0,0,0,0,0,0,1,0,0,0,0,0,0,0,0,0,0,0,0,0,0,0,0,0,0,0,0,0,0,0,0,0,0,0,0,0,0,0,0,0,0)

PC1.8M2<-c(0,0,0,0,0,0,0,0,0,0,0,0,0,0,0,0,0,0,0,0,0,0,0,0,0,0,0,0,0,1,0,0,0,0,0,0,0,0,0,0,0,0,0,0,0,0,0,0,0,0,0,0,0,0,0,0,0,0,0,0,0,0,0)

PC1.5M2<-c(0,0,0,0,0,0,0,0,0,0,0,0,0,0,0,0,0,0,0,0,0,0,0,0,0,0,0,0,0,0,1,0,0,0,0,0,0,0,0,0,0,0,0,0,0,0,0,0,0,0,0,0,0,0,0,0,0,0,0,0,0,0,0)

PC1.3M2<-c(0,0,0,0,0,0,0,0,0,0,0,0,0,0,0,0,0,0,0,0,0,0,0,0,0,0,0,0,0,0,0,1,0,0,0,0,0,0,0,0,0,0,0,0,0,0,0,0,0,0,0,0,0,0,0,0,0,0,0,0,0,0,0)

PC1.1M2<-c(0,0,0,0,0,0,0,0,0,0,0,0,0,0,0,0,0,0,0,0,0,0,0,0,0,0,0,0,0,0,0,0,1,0,0,0,0,0,0,0,0,0,0,0,0,0,0,0,0,0,0,0,0,0,0,0,0,0,0,0,0,0,0)

PC5.1M2<-c(0,0,0,0,0,0,0,0,0,0,0,0,0,0,0,0,0,0,0,0,0,0,0,0,0,0,0,0,0,0,0,0,0,1,0,0,0,0,0,0,0,0,0,0,0,0,0,0,0,0,0,0,0,0,0,0,0,0,0,0,0,0,0)

PC8.1M2<-c(0,0,0,0,0,0,0,0,0,0,0,0,0,0,0,0,0,0,0,0,0,0,0,0,0,0,0,0,0,0,0,0,0,0,1,0,0,0,0,0,0,0,0,0,0,0,0,0,0,0,0,0,0,0,0,0,0,0,0,0,0,0,0)

PC3.1L2<-c(0,0,0,0,0,0,0,0,0,0,0,0,0,0,0,0,0,0,0,0,0,0,0,0,0,0,0,0,0,0,0,0,0,0,0,1,0,0,0,0,0,0,0,0,0,0,0,0,0,0,0,0,0,0,0,0,0,0,0,0,0,0,0)

PC1.8L2<-c(0,0,0,0,0,0,0,0,0,0,0,0,0,0,0,0,0,0,0,0,0,0,0,0,0,0,0,0,0,0,0,0,0,0,0,0,1,0,0,0,0,0,0,0,0,0,0,0,0,0,0,0,0,0,0,0,0,0,0,0,0,0,0)

PC1.5L2<-c(0,0,0,0,0,0,0,0,0,0,0,0,0,0,0,0,0,0,0,0,0,0,0,0,0,0,0,0,0,0,0,0,0,0,0,0,0,1,0,0,0,0,0,0,0,0,0,0,0,0,0,0,0,0,0,0,0,0,0,0,0,0,0)

PC1.3L2<-c(0,0,0,0,0,0,0,0,0,0,0,0,0,0,0,0,0,0,0,0,0,0,0,0,0,0,0,0,0,0,0,0,0,0,0,0,0,0,1,0,0,0,0,0,0,0,0,0,0,0,0,0,0,0,0,0,0,0,0,0,0,0,0)

PC1.1L2<-c(0,0,0,0,0,0,0,0,0,0,0,0,0,0,0,0,0,0,0,0,0,0,0,0,0,0,0,0,0,0,0,0,0,0,0,0,0,0,0,1,0,0,0,0,0,0,0,0,0,0,0,0,0,0,0,0,0,0,0,0,0,0,0)

PC5.1L2<-c(0,0,0,0,0,0,0,0,0,0,0,0,0,0,0,0,0,0,0,0,0,0,0,0,0,0,0,0,0,0,0,0,0,0,0,0,0,0,0,0,1,0,0,0,0,0,0,0,0,0,0,0,0,0,0,0,0,0,0,0,0,0,0)

PC8.1L2<-c(0,0,0,0,0,0,0,0,0,0,0,0,0,0,0,0,0,0,0,0,0,0,0,0,0,0,0,0,0,0,0,0,0,0,0,0,0,0,0,0,0,1,0,0,0,0,0,0,0,0,0,0,0,0,0,0,0,0,0,0,0,0,0)

PC3.1H3<-c(0,0,0,0,0,0,0,0,0,0,0,0,0,0,0,0,0,0,0,0,0,0,0,0,0,0,0,0,0,0,0,0,0,0,0,0,0,0,0,0,0,0,1,0,0,0,0,0,0,0,0,0,0,0,0,0,0,0,0,0,0,0,0)

PC1.8H3<-c(0,0,0,0,0,0,0,0,0,0,0,0,0,0,0,0,0,0,0,0,0,0,0,0,0,0,0,0,0,0,0,0,0,0,0,0,0,0,0,0,0,0,0,1,0,0,0,0,0,0,0,0,0,0,0,0,0,0,0,0,0,0,0)

PC1.5H3<-c(0,0,0,0,0,0,0,0,0,0,0,0,0,0,0,0,0,0,0,0,0,0,0,0,0,0,0,0,0,0,0,0,0,0,0,0,0,0,0,0,0,0,0,0,1,0,0,0,0,0,0,0,0,0,0,0,0,0,0,0,0,0,0)

PC1.3H3<-c(0,0,0,0,0,0,0,0,0,0,0,0,0,0,0,0,0,0,0,0,0,0,0,0,0,0,0,0,0,0,0,0,0,0,0,0,0,0,0,0,0,0,0,0,0,1,0,0,0,0,0,0,0,0,0,0,0,0,0,0,0,0,0)

PC1.1H3<-c(0,0,0,0,0,0,0,0,0,0,0,0,0,0,0,0,0,0,0,0,0,0,0,0,0,0,0,0,0,0,0,0,0,0,0,0,0,0,0,0,0,0,0,0,0,0,1,0,0,0,0,0,0,0,0,0,0,0,0,0,0,0,0)

PC5.1H3<-c(0,0,0,0,0,0,0,0,0,0,0,0,0,0,0,0,0,0,0,0,0,0,0,0,0,0,0,0,0,0,0,0,0,0,0,0,0,0,0,0,0,0,0,0,0,0,0,1,0,0,0,0,0,0,0,0,0,0,0,0,0,0,0)

PC8.1H3<-c(0,0,0,0,0,0,0,0,0,0,0,0,0,0,0,0,0,0,0,0,0,0,0,0,0,0,0,0,0,0,0,0,0,0,0,0,0,0,0,0,0,0,0,0,0,0,0,0,1,0,0,0,0,0,0,0,0,0,0,0,0,0,0)

PC3.1M3<-c(0,0,0,0,0,0,0,0,0,0,0,0,0,0,0,0,0,0,0,0,0,0,0,0,0,0,0,0,0,0,0,0,0,0,0,0,0,0,0,0,0,0,0,0,0,0,0,0,0,1,0,0,0,0,0,0,0,0,0,0,0,0,0)

PC1.8M3<-c(0,0,0,0,0,0,0,0,0,0,0,0,0,0,0,0,0,0,0,0,0,0,0,0,0,0,0,0,0,0,0,0,0,0,0,0,0,0,0,0,0,0,0,0,0,0,0,0,0,0,1,0,0,0,0,0,0,0,0,0,0,0,0)

PC1.5M3<-c(0,0,0,0,0,0,0,0,0,0,0,0,0,0,0,0,0,0,0,0,0,0,0,0,0,0,0,0,0,0,0,0,0,0,0,0,0,0,0,0,0,0,0,0,0,0,0,0,0,0,0,1,0,0,0,0,0,0,0,0,0,0,0)

PC1.3M3<-c(0,0,0,0,0,0,0,0,0,0,0,0,0,0,0,0,0,0,0,0,0,0,0,0,0,0,0,0,0,0,0,0,0,0,0,0,0,0,0,0,0,0,0,0,0,0,0,0,0,0,0,0,1,0,0,0,0,0,0,0,0,0,0)

PC1.1M3<-c(0,0,0,0,0,0,0,0,0,0,0,0,0,0,0,0,0,0,0,0,0,0,0,0,0,0,0,0,0,0,0,0,0,0,0,0,0,0,0,0,0,0,0,0,0,0,0,0,0,0,0,0,0,1,0,0,0,0,0,0,0,0,0)

PC5.1M3<-c(0,0,0,0,0,0,0,0,0,0,0,0,0,0,0,0,0,0,0,0,0,0,0,0,0,0,0,0,0,0,0,0,0,0,0,0,0,0,0,0,0,0,0,0,0,0,0,0,0,0,0,0,0,0,1,0,0,0,0,0,0,0,0)

PC8.1M3<-c(0,0,0,0,0,0,0,0,0,0,0,0,0,0,0,0,0,0,0,0,0,0,0,0,0,0,0,0,0,0,0,0,0,0,0,0,0,0,0,0,0,0,0,0,0,0,0,0,0,0,0,0,0,0,0,1,0,0,0,0,0,0,0)

PC3.1L3<-c(0,0,0,0,0,0,0,0,0,0,0,0,0,0,0,0,0,0,0,0,0,0,0,0,0,0,0,0,0,0,0,0,0,0,0,0,0,0,0,0,0,0,0,0,0,0,0,0,0,0,0,0,0,0,0,0,1,0,0,0,0,0,0)

PC1.8L3<-c(0,0,0,0,0,0,0,0,0,0,0,0,0,0,0,0,0,0,0,0,0,0,0,0,0,0,0,0,0,0,0,0,0,0,0,0,0,0,0,0,0,0,0,0,0,0,0,0,0,0,0,0,0,0,0,0,0,1,0,0,0,0,0)

PC1.5L3<-c(0,0,0,0,0,0,0,0,0,0,0,0,0,0,0,0,0,0,0,0,0,0,0,0,0,0,0,0,0,0,0,0,0,0,0,0,0,0,0,0,0,0,0,0,0,0,0,0,0,0,0,0,0,0,0,0,0,0,1,0,0,0,0)

PC1.3L3<-c(0,0,0,0,0,0,0,0,0,0,0,0,0,0,0,0,0,0,0,0,0,0,0,0,0,0,0,0,0,0,0,0,0,0,0,0,0,0,0,0,0,0,0,0,0,0,0,0,0,0,0,0,0,0,0,0,0,0,0,1,0,0,0)

PC1.1L3<-c(0,0,0,0,0,0,0,0,0,0,0,0,0,0,0,0,0,0,0,0,0,0,0,0,0,0,0,0,0,0,0,0,0,0,0,0,0,0,0,0,0,0,0,0,0,0,0,0,0,0,0,0,0,0,0,0,0,0,0,0,1,0,0)

PC5.1L3<-c(0,0,0,0,0,0,0,0,0,0,0,0,0,0,0,0,0,0,0,0,0,0,0,0,0,0,0,0,0,0,0,0,0,0,0,0,0,0,0,0,0,0,0,0,0,0,0,0,0,0,0,0,0,0,0,0,0,0,0,0,0,1,0)

PC8.1L3<-c(0,0,0,0,0,0,0,0,0,0,0,0,0,0,0,0,0,0,0,0,0,0,0,0,0,0,0,0,0,0,0,0,0,0,0,0,0,0,0,0,0,0,0,0,0,0,0,0,0,0,0,0,0,0,0,0,0,0,0,0,0,0,1)

All.HRs.Contrasts.to.RefCat.m3<- contrast(All.HRs.Strata.m3,adjust="none", CI=TRUE, method=list(

+	(PC1.8L1-PC3.1L1),

+	(PC1.5L1-PC3.1L1),

+	(PC1.3L1-PC3.1L1),

+	(PC1.1L1-PC3.1L1),

+	(PC3.1L1-PC3.1L1),

+	(PC5.1L1-PC3.1L1),

+	(PC8.1L1-PC3.1L1),

+	(PC1.8M1-PC3.1L1),

+	(PC1.5M1-PC3.1L1),

+	(PC1.3M1-PC3.1L1),

+	(PC1.1M1-PC3.1L1),

+	(PC3.1M1-PC3.1L1),

+	(PC5.1M1-PC3.1L1),

+	(PC8.1M1-PC3.1L1),

+	(PC1.8H1-PC3.1L1),

+	(PC1.5H1-PC3.1L1),

+	(PC1.3H1-PC3.1L1),

+	(PC1.1H1-PC3.1L1),

+	(PC3.1H1-PC3.1L1),

+	(PC5.1H1-PC3.1L1),

+	(PC8.1H1-PC3.1L1),

(PC1.8L2-PC3.1L2),

+	(PC1.5L2-PC3.1L2),

+	(PC1.3L2-PC3.1L2),

+	(PC1.1L2-PC3.1L2),

+	(PC3.1L2-PC3.1L2),

+	(PC5.1L2-PC3.1L2),

+	(PC8.1L2-PC3.1L2),

+	(PC1.8M2-PC3.1L2),

+	(PC1.5M2-PC3.1L2),

+	(PC1.3M2-PC3.1L2),

+	(PC1.1M2-PC3.1L2),

+	(PC3.1M2-PC3.1L2),

+	(PC5.1M2-PC3.1L2),

+	(PC8.1M2-PC3.1L2),

+	(PC1.8H2-PC3.1L2),

+	(PC1.5H2-PC3.1L2),

+	(PC1.3H2-PC3.1L2),

+	(PC1.1H2-PC3.1L2),

+	(PC3.1H2-PC3.1L2),

+	(PC5.1H2-PC3.1L2),

+	(PC8.1H2-PC3.1L2),

(PC1.8L3-PC3.1L3),

+	(PC1.5L3-PC3.1L3),

+	(PC1.3L3-PC3.1L3),

+	(PC1.1L3-PC3.1L3),

+	(PC3.1L3-PC3.1L3),

+	(PC5.1L3-PC3.1L3),

+	(PC8.1L3-PC3.1L3),

+	(PC1.8M3-PC3.1L3),

+	(PC1.5M3-PC3.1L3),

+	(PC1.3M3-PC3.1L3),

+	(PC1.1M3-PC3.1L3),

+	(PC3.1M3-PC3.1L3),

+	(PC5.1M3-PC3.1L3),

+	(PC8.1M3-PC3.1L3),

+	(PC1.8H3-PC3.1L3),

+	(PC1.5H3-PC3.1L3),

+	(PC1.3H3-PC3.1L3),

+	(PC1.1H3-PC3.1L3),

+	(PC3.1H3-PC3.1L3),

+	(PC5.1H3-PC3.1L3),

+	(PC8.1H3-PC3.1L3)

))

All.HRs.Contrasts.to.RefCat.m3

\## PLOTS - SURVIVAL TIME - Time stratified

#Use the All.HRs.Strata.m3 table

plotdata<-as.data.frame(All.HRs.Strata.m3)

str(plotdata)

#Set the order of different levels of the diet factors so they appear in the correct order in the plot

plotdata$Cellulose <- factor(plotdata$Cellulose, levels = c('L', 'M', 'H'))

plotdata$P.C <- factor(plotdata$P.C, levels = c('1:8','1:5','1:3','1:1','3:1','5:1','8:1'))

#Separate the plotdata by Time

Time1.plotdata <- plotdata[ which(plotdata$Time=='1'), ]

str(Time1.plotdata)

Time1.plotdata$Cellulose <- factor(Time1.plotdata$Cellulose, levels = c('L', 'M', 'H'))

Time1.plotdata$P.C <- factor(Time1.plotdata$P.C, levels = c('1:8','1:5','1:3','1:1','3:1','5:1','8:1'))

Time2.plotdata <- plotdata[ which(plotdata$Time=='2'), ]

str(Time2.plotdata)

Time3.plotdata <- plotdata[ which(plotdata$Time=='3'), ]

str(Time3.plotdata)

\# A an adjustment to use later to make the plot elements not overlap

pd <- position_dodge(0.6) #increase or decrease value to change the distance between the plot elements

Time1.plotdata$P.C <- factor(Time1.plotdata$P.C, levels = c('1:8', '1:5', '1:3', '1:1', '3:1', '5:1', '8:1'))

#PLOT For Time.Interval = 1

ST.Plot<-ggplot(Time1.plotdata, aes(x=P.C, y=response, colour = Cellulose, group=Cellulose))+

#annotate("text", x="8:1", y=20, label="0-7 days", size=6)+

geom_errorbar(aes(ymin=asymp.LCL, ymax=asymp.UCL), width=0.2,lwd=0.9,position=pd, size=1)+

geom_line(position=pd, size=1.2)+

geom_point(position=pd, size = 2)+

geom_hline(yintercept = 1, linewidth = 0.5, lty =2, colour = "black")+ # horizontal line showing where risk of death switches from increasing to decreasing

scale_colour_manual(values = cols, labels=c('Low', 'Medium', 'High'))+ #Change cellulose legend labels here

scale_y_continuous(limits = c(0.1,20),breaks=c(0,5,10,15,20))+ # Change y axis limits and labels here

labs(x = "P:C",y = "Mortality Hazard Ratio", colour = "Cellulose")+ # Change x and y axis titles here

theme(panel.border = element_rect(colour = "black", fill=NA, linewidth=0.5),

panel.grid.major = element_blank(),

panel.grid.minor = element_blank(),

panel.background = element_blank(),

axis.line = element_line(colour = "black"),

axis.text=element_text(size=12),

axis.title=element_text(size=16),

legend.key=element_blank(),

legend.text = element_text(size=16),

legend.title = element_text(size=16))

ST.Plot

ST.Plot + geom_text(x = "8:1", y = 20, label = "0-7 days", size = 6)

\# troubleshooting: if you get a margins or graphics state error, clear the plot history and increase the size of the plot area

#saves the last plot that was created, but there is a time limit - rerun the plot code if this doesnt work

#IMPORTANT: Change the

ggsave("SurvTimePlot.Time1.jpeg", width = 15, height = 10, units = "cm", dpi = 500, path = "...")

#PLOT For Time.Interval = 2

ST.Plot2<-ggplot(Time2.plotdata, aes(x=P.C,y=response, colour = Cellulose, group=Cellulose))+

#annotate("text", x="8:1", y=20, label="7-14 days", size=6)+

geom_errorbar(aes(ymin=asymp.LCL, ymax=asymp.UCL), width=0.2,lwd=0.9,position=pd, size=1)+

geom_line(position=pd, size=1.2)+

geom_point(position=pd, size = 2)+

geom_hline(yintercept = 1, linewidth = 0.5, lty =2, colour = "black")+ # horizontal line showing where risk of death switches from increasing to decreasing

#scale_x_discrete(labels=c('label1', 'label2', 'label3', ...)+ # This code could be used to change x axis labels

scale_colour_manual(values = cols, labels=c('Low', 'Medium', 'High'))+ #Change cellulose legend labels here

scale_y_continuous(limits = c(0.1,20),breaks=c(0,5,10,15,20))+ # Change y axis limits and labels here

labs(x = "P:C",y = "Mortality Hazard Ratio", colour = "Cellulose")+ # Change x and y axis titles here

theme(panel.border = element_rect(colour = "black", fill=NA, linewidth=0.5),

panel.grid.major = element_blank(),

panel.grid.minor = element_blank(),

panel.background = element_blank(),

axis.line = element_line(colour = "black"),

axis.text=element_text(size=12),

axis.title=element_text(size=16),

legend.key=element_blank(),

legend.text = element_text(size=16),

legend.title = element_text(size=16))

ST.Plot2

ggsave("SurvTimePlot.Time2.jpeg", width = 15, height = 10, units = "cm", dpi = 500, path = "...")

#PLOT For Time.Interval = 3

ST.Plot3<-ggplot(Time3.plotdata, aes(x=P.C,y=response, colour = Cellulose, group=Cellulose))+

#annotate("text", x="8:1", y=20, label="7-14 days", size=6)+

geom_errorbar(aes(ymin=asymp.LCL, ymax=asymp.UCL), width=0.2,lwd=0.9,position=pd, size=1)+

geom_line(position=pd, size=1.2)+

geom_point(position=pd, size = 2)+

geom_hline(yintercept = 1, linewidth = 0.5, lty =2, colour = "black")+ # horizontal line showing where risk of death switches from increasing to decreasing

#scale_x_discrete(labels=c('label1', 'label2', 'label3', ...)+ # This code could be used to change x axis labels

scale_colour_manual(values = cols, labels=c('Low', 'Medium', 'High'))+ #Change cellulose legend labels here

scale_y_continuous(limits = c(0.1,20),breaks=c(0,5,10,15,20))+ # Change y axis limits and labels here

labs(x = "P:C",y = "Mortality Hazard Ratio", colour = "Cellulose")+ # Change x and y axis titles here

theme(panel.border = element_rect(colour = "black", fill=NA, linewidth=0.5),

panel.grid.major = element_blank(),

panel.grid.minor = element_blank(),

panel.background = element_blank(),

axis.line = element_line(colour = "black"),

axis.text=element_text(size=12),

axis.title=element_text(size=16),

legend.key=element_blank(),

legend.text = element_text(size=16),

legend.title = element_text(size=16))

ST.Plot3

\# troubleshooting: if you get a margins or graphics state error, clear the plot history and increase the size of the plot area

ggsave("SurvTimePlot.Time3.jpeg", width = 15, height = 10, units = "cm", dpi = 500, path = "...")

#############Yield Calculations################

library(car)#for 2 and 3 way Anova

library(writexl)

library(readxl)

library(ggplot2)

library(MASS) #for AIC model selection

library(tidyverse)

library(dplyr)

DATA <- read_excel(".../Muzzatti.et.al.2024.Data.xlsx", sheet = "dev and survival")

DATA$Cellulose=as.factor(DATA$Cellulose)

DATA$P.C=as.factor(DATA$P.C)

DATA$Status.Adult=as.numeric(DATA$Status.Adult)

DATA$Status.Death=as.numeric(DATA$Status.Death)

DATA$P.C <- factor(DATA$P.C, levels = c('1:8', '1:5','1:3','1:1','3:1','5:1','8:1'))

DATA$Cellulose<-factor(DATA$Cellulose, levels=c("L","M","H"))

#we want means for each P.C category and each cellulose category of adult mass, body size, proportion surviving, and development time

#need to filter by Status.Adult because there were some crickets that survived to adulthood, but then died during the following week they were kept alive.

summary_table <- DATA %>%

group_by(PC.Ratio, Cellulose) %>%

summarise(

avg_mass = mean(Adult_mass_g[Status.Adult == 1], na.rm = TRUE),

avg_surviving = mean(1-Status.Death, na.rm = TRUE), #Status.Death = 1 means individual died. Need 1-column for survival

avg_development_time = mean(Dev.Time[Status.Adult == 1], na.rm = TRUE))

summary_table <- as.data.frame(summary_table)

summary_table

summary_table <- summary_table %>%

mutate(

yield = (avg_mass * avg_surviving)/avg_development_time

) %>%

mutate(logPC = log(PC.Ratio))

summary_table

print(summary_table)

summary_table3 <- DATA %>%

group_by(PC.Ratio) %>%

summarise(

avg_mass = mean(Adult_mass_g[Status.Adult == 1], na.rm = TRUE),

avg_surviving = mean(1-Status.Death, na.rm = TRUE),

avg_development_time = mean(Dev.Time[Status.Adult == 1], na.rm = TRUE),

SE_mass = sd(Adult_mass_g, na.rm = TRUE) / sqrt(n()),

SE_surviving = sqrt(avg_surviving * (1 - avg_surviving) / n()),

SE_development_time = sd(Dev.Time, na.rm = TRUE) / sqrt(n()))

summary_table3 <- as.data.frame(summary_table3)

summary_table3

summary_table3 <- summary_table3 %>%

mutate(

yield = (avg_mass * avg_surviving)/avg_development_time

) %>%

mutate(logPC = log(PC.Ratio))

summary_table3

print(summary_table3)

####choice####

library(car)

library(MASS)

require(car)

require(MASS)

library(ggplot2)

library(sjPlot)

library(mlogit)

library(nnet)

library(ggplot2)##This package needs to be called first before sjplot

library(lme4)

require(lme4)#THis is where glmer is

library(cAIC4)

library(MuMIn)

library(lmerTest)# does the same thing as lme4 (so loading it masks lme4), but gives denominator df and p values

library(sjPlot)

library(multcomp)

library(effects)

library(xlsx)

library(nlme)

library(readxl)

library(writexl)

library(ggpubr)

library(splines)

data <- read_xlsx(".../Muzzatti.et.al.2024.Data.xlsx", sheet = "Choice")

data$Diet=as.factor(data$Diet)

data$Sex=as.factor(data$Sex)

data$Period=as.factor(data$Period)

str(data)

PC.ChoiceM1 <- lme(Amt.Eaten.g ~Wk1A.BodySizePC1+Week*Diet,random = ~1|ID, data)

summary(PC.ChoiceM1)

anova(PC.ChoiceM1)

plot(PC.ChoiceM1)#Lmm isnt a good fit - need to switch to glmm

PC.ChoiceM2 <- lme(Amt.Eaten.g ~Wk1A.BodySizePC1+I(Week^2)*Diet,random = ~1|ID, data)

summary(PC.ChoiceM2)

plot(PC.ChoiceM2)

anova(PC.ChoiceM1,PC.ChoiceM2) # M1 has the highest logLik, so it is a better fit

qqp(data$Amt.Eaten.g, "norm")# Isnt normal

qqp(data$Amt.Eaten.g, "lnorm")# Isnt Log normal, but closer

#Gamma fits the dependent best

gamma <- fitdistr(data$Amt.Eaten.g, "gamma")

qqp(data$Amt.Eaten.g, "gamma", shape = gamma$estimate[1], rate=gamma$estimate[2])

m4 <- glmer(Amt.Eaten.g ~ Wk1A.BodySizePC1+Week*Diet + (1|ID), data=data, family=Gamma(link="log"), control=glmerControl((optimizer="bobyqa"),optCtrl=list(maxfun=20000)))

m5 <- glmer(Amt.Eaten.g ~ Wk1A.BodySizePC1+(I(Week^2))*Diet + (1|ID), data=data, family=Gamma(link="log"), control=glmerControl((optimizer="bobyqa"),optCtrl=list(maxfun=20000)))

anova(m4,m5) #m4 has the higher loglik, so include week as linear

summary(m4)

m6 <- glmer(Amt.Eaten.g ~ Wk1A.BodySizePC1+Week+Diet+Sex+ (1|ID), data=data,family=Gamma(link="log"), control=glmerControl((optitimizer="bobyqa"),optCtrl=list(maxfun=20000)))

m7 <- glmer(Amt.Eaten.g ~ Wk1A.BodySizePC1+(I(Week^2))+Diet+Sex+ (1|ID), data=data,family=Gamma(link="log"), control=glmerControl((optitimizer="bobyqa"),optCtrl=list(maxfun=20000)))

anova(m6,m7)#m6 has higher loglik - include week as linear

m8 <- glmer(Amt.Eaten.g ~ Wk1A.BodySizePC1+Week*Diet*Sex + (1|ID), data=data,family=Gamma(link="log"), control=glmerControl((optitimizer="bobyqa"),optCtrl=list(maxfun=20000)))

summary(m8)

library(MuMIn)

options(na.action= "na.fail")

m8.Global.mod.eval<-dredge(m8,rank=AIC,evaluate = TRUE,fixed=~Wk1A.BodySizePC1+Week+Diet+Sex) # n = 446 obs, 9 terms in m8, 446/9=49. >40 so can use AIC, not AICc

m8.Global.mod.eval

m8.subset4<-subset(m8.Global.mod.eval, delta < 4,recalc.weights = TRUE,recalc.delta=TRUE)# Weight column recalculated to sum to 1

m8.subset4

nested(m8.subset4) # gives a T/F list whether model X has other models 'nested' within it

nst <- nested(m8.subset4, indices = "row")

nst # print model "x" and all models nested within it

m8.subset4$nested <- sapply(nst, paste, collapse = ",")

m8.subset4 #Original model selection table with the nested models indicated

m8.nest.reduced.subset4<-subset(m8.subset4, !nested(.)) # Automatically filter out overly complex models according to the "nesting selection rule"

m8.nest.reduced.subset4 # One model has full support

FinalMod<- (get.models(m8.nest.reduced.subset4, 1)[[1]])

summary(FinalMod)

Anova(FinalMod, type = 3)# Use type 3 sums of squares because there is a significant interaction

FinalMod<- glmer(Amt.Eaten.g ~ Wk1A.BodySizePC1+Week+Diet+Sex+Diet:Sex + (1|ID), data=data,family=Gamma(link="log"), control=glmerControl((optitimizer="bobyqa"),optCtrl=list(maxfun=20000)))

summary(FinalMod)

Anova(FinalMod, type = 3)# Use type 3 sums of squares because there is a significant interaction

m9 <- glmer(Amt.Eaten.g ~ Wk1A.BodySizePC1+Period*Diet*Sex + (1|ID), data=data, family=Gamma(link="log"), control=glmerControl((optimizer="bobyqa"),optCtrl=list(maxfun=20000)))

summary(m9)

library(effects)

Int.effs<-effect(m8, term="Diet:Sex", xlevels=100, se=TRUE)

Int.effs<-as.data.frame(Int.effs)

Int.effs #for some reason the CIs only appear after you make it a data frame

write_xlsx(Int.effs,"...")

df<- read_excel("...")

df<-as.data.frame(df)

str(df)

pd <- position_dodge(0.1) # move them .05 to the left and right

Plot<-ggplot(df, aes(x=Diet,y=Amount.Eaten, colour = Sex, group=Sex))+

geom_errorbar(aes(ymin=lwr, ymax=upr),colour= c("aquamarine3","aquamarine3","darkred","darkred"), width=0.2,lwd=0.9,position=pd)+

geom_line(position=pd, colour= c("aquamarine3","aquamarine3","darkred","darkred"))+

geom_point(position=pd, size = 3, colour= c("aquamarine3","aquamarine3","darkred","darkred"))+

labs(x = "Diet",y = "Amount of Diet Eaten (g)")

Plot+theme(panel.border = element_rect(colour = "black", fill=NA, size=0.5),

panel.grid.major = element_blank(),

panel.grid.minor = element_blank(),

panel.background = element_blank(),

axis.line = element_line(colour = "black"),

axis.text=element_text(size=12),

axis.title=element_text(size=16))

Plot<-ggplot(df, aes(x=Diet,y=fit, colour = Sex, group=Sex))+

geom_errorbar(aes(ymin=lower, ymax=upper),colour= c("aquamarine3","aquamarine3","darkred","darkred"), width=0.2,lwd=0.9,position=pd)+

geom_line(position=pd, colour= c("aquamarine3","aquamarine3","darkred","darkred"))+

geom_point(position=pd, size = 3, colour= c("aquamarine3","aquamarine3","darkred","darkred"))+

labs(x = "Diet",y = "Amount of Diet Eaten (g)")

Plot+theme(panel.border = element_rect(colour = "black", fill=NA, size=0.5),

panel.grid.major = element_blank(),

panel.grid.minor = element_blank(),

panel.background = element_blank(),

axis.line = element_line(colour = "black"),

axis.text=element_text(size=12),

axis.title=element_text(size=16))

Plot <- ggplot(df, aes(x = Diet, y = fit, group = Sex)) +

geom_errorbar(aes(ymin = fit - se, ymax = fit + se, colour = Sex), width = 0.2, lwd = 0.9, position = pd) +

geom_point(aes(colour = Sex), position = pd, size = 3) +

labs(x = "", y = "Amount of Diet Eaten (g)") +

theme(panel.border = element_rect(colour = "black", fill = NA, size = 0.5),

panel.grid.major = element_blank(),

panel.grid.minor = element_blank(),

panel.background = element_blank(),

axis.line = element_line(colour = "black"),

axis.text = element_text(size = 12),

axis.title = element_text(size = 16)) +

scale_color_manual(values = c("aquamarine3", "darkred"), labels = c("Male", "Female")) +

scale_x_discrete(labels = c("Carbohydrate", "Protein"))

Plot

library(car)

library(MASS)

require(car)

require(MASS)

library(ggplot2)

library(sjPlot)

library(mlogit)

library(nnet)

library(ggplot2)##This package needs to be called first before sjplot

library(lme4)

require(lme4)#THis is where glmer is

library(cAIC4)

library(MuMIn)

library(lmerTest)# does the same thing as lme4 (so loading it masks lme4), but gives denominator df and p values

library(sjPlot)

library(multcomp)

library(effects)

library(xlsx)

library(nlme)

library(readxl)

library(writexl)

library(ggpubr)

library(splines)

library(ggpubr)

library(xlsx)

library(readxl)

library(ggplot2)

#make the "Data" tab the active sheet before opening

data <- read_xlsx(".../Muzzatti.et.al.2024.Data.xlsx", sheet = "Choice_ttest")

data$Sex=as.factor(data$Sex)

str(data)

#Split the data into males and females

mdata <- data[ which(data$Sex=='M'), ]

str(mdata)

fdata <- data[ which(data$Sex=='F'), ]

str(fdata)

mdata$Sex=as.factor(mdata$Sex)

str(mdata)

fdata$Sex=as.factor(fdata$Sex)

str(fdata)

shapiro.test(mdata$PC.Ratio)# P is less than 0.05, so significantly different than normal

ggqqplot(mdata$PC.Ratio, ggtheme = theme_minimal())#Not Normal (positive skew)

#Sqrt transformed -best

shapiro.test(mdata$Sqrt.PC.Ratio)# P is greater than 0.05, so not significantly different than normal

ggqqplot(mdata$Sqrt.PC.Ratio, ggtheme = theme_minimal())#Normal

#Females

#Untransformed

shapiro.test(fdata$PC.Ratio)# P is less than 0.05, so significantly different than normal

ggqqplot(fdata$PC.Ratio, ggtheme = theme_minimal())#Not Normal (positive skew)

#Sqrt Transformed

shapiro.test(fdata$Sqrt.PC.Ratio)# P is less than 0.05, so significantly different than normal

ggqqplot(fdata$Sqrt.PC.Ratio, ggtheme = theme_minimal())#Not Normal

#Log10 Transformed

shapiro.test(fdata$Log10.PC.Ratio)# P is less than 0.05, so significantly different than normal

ggqqplot(fdata$Log10.PC.Ratio, ggtheme = theme_minimal())#Not Normal

#Inverse Transformed - Best

shapiro.test(fdata$Inv.PC.Ratio)# P is greater than 0.05, so not significantly different than normal

ggqqplot(fdata$Inv.PC.Ratio, ggtheme = theme_minimal())# Normal

#All Individuals

#Untransformed

shapiro.test(data$PC.Ratio)# P is less than 0.05, so significantly different than normal

ggqqplot(data$PC.Ratio, ggtheme = theme_minimal())#Not Normal (positive skew)

#Sqrt Transformed

shapiro.test(data$Sqrt.PC.Ratio)# P is less than 0.05, so significantly different than normal

ggqqplot(data$Sqrt.PC.Ratio, ggtheme = theme_minimal())#Not Normal

#Log10 Transformed -Best

shapiro.test(data$Log10.PC.Ratio)# P is greater than 0.05, so not significantly different than normal

ggqqplot(data$Log10.PC.Ratio, ggtheme = theme_minimal())#Normal

#Males - Sqrt Transformed

Males <- t.test(mdata$Sqrt.PC.Ratio, mu = 1,alternative = "two.sided")

Males #Back-transform for Sqrt with (mean of x)^2

#Females - Inverse Transformed

Females <- t.test(fdata$Inv.PC.Ratio, mu = 1,alternative = "two.sided")

Females #Back-transform for inverse with 1/(mean of x)

#All - Log 10 Transformed

All <- t.test(data$Log10.PC.Ratio, mu = 1,alternative = "two.sided")

All #Back-transform for Log10 with 10^(mean of x)
